# Evolution of a chordate-specific mechanism for myoblast fusion

**DOI:** 10.1101/2021.07.24.453587

**Authors:** Haifeng Zhang, Renjie Shang, Kwantae Kim, Wei Zheng, Christopher J. Johnson, Lei Sun, Xiang Niu, Liang Liu, Theodore A. Uyeno, Jingqi Zhou, Lingshu Liu, Jimin Pei, Skye D. Fissette, Stephen A. Green, Sukhada P. Samudra, Junfei Wen, Jianli Zhang, Jonathan Eggenschwiler, Doug Menke, Marianne E. Bronner, Nick V. Grishin, Weiming Li, Kaixiong Ye, Yang Zhang, Alberto Stolfi, Pengpeng Bi

**Author notes:** These authors contributed equally (Haifeng Zhang, Renjie Shang). Corresponding Author: Pengpeng Bi; Alberto Stolfi.

## Abstract

The size of an animal is determined by the size of its musculoskeletal system. Myoblast fusion is an innovative mechanism that allows for multinucleated muscle fibers to compound the size and strength of individual mononucleated cells. However, the evolutionary history of the control mechanism underlying this important process is currently unknown. The phylum Chordata hosts closely related groups that span distinct myoblast fusion states: no fusion in cephalochordates, restricted fusion and multinucleation in tunicates, and extensive, obligatory fusion in vertebrates. To elucidate how these differences may have evolved, we studied the evolutionary origins and function of membrane-coalescing agents Myomaker and Myomixer in various groups of chordates. Here we report that *Myomaker* likely arose through gene duplication in the last common ancestor of tunicates and vertebrates, while *Myomixer* appears to have evolved *de novo* in early vertebrates. Functional tests revealed an unexpectedly complex evolutionary history of myoblast fusion in chordates. A pre-vertebrate phase of muscle multinucleation driven by Myomaker was followed by the later emergence of Myomixer that enables the highly efficient fusion system of vertebrates. Thus, our findings reveal the evolutionary origins of chordate-specific fusogens and illustrate how new genes can shape the emergence of novel morphogenetic traits and mechanisms.

## Introduction

The robustness of the musculoskeletal system is crucial for an animal’s fitness in its challenging environment. Vertebrate myogenesis involves a series of events that begins with the specification of muscle lineage precursors by transcriptional regulators like Pax7 and MyoD, followed by the expression of a vast number of genes that establish muscle structure and function^1, 2^. A fundamental step in this process is the fusion of mononucleated myoblasts to form multinucleated myofibers^3–7^. Generation of syncytial myofibers allows concerted power outputs to fulfill complex locomotor functions, and therefore was likely instrumental for the adaptive radiation of vertebrates. However, the evolutionary origins of vertebrate myoblast fusion are currently unknown.

Recent genetic studies in mice uncovered two muscle-specific fusogens, Myomaker (MymK) and Myomixer (MymX) that jointly drive myoblast fusion^8–11^. Deletion of either gene causes perinatal lethality of mice due to fusion defects that result in muscle malfunction^8, 9^. The expression of this duo is tightly controlled by MyoD and restricted to the precise time window of fusion during muscle development and regeneration^12–14^. *MymK* encodes a 7-pass transmembrane protein, with a topology similar to that of G protein-coupled receptors^15^. In contrast, *MymX* encodes a small single-pass membrane protein that requires MymK to induce fusion of myoblasts^9^. Reconstitution experiments established the model in which MymK can induce formation of small myotubes, whereas adding MymX boosts the fusion efficiency of MymK and the generation of large muscle syncytia^12^. Moreover, forced expression of this duo confers fusogenic activity even onto fibroblasts, which are not normally capable of undergoing cell fusion^9^.

Despite this emerging knowledge of MymX and MymK function, the evolutionary origins of these crucial agents remain elusive. Uncovering their evolutionary histories holds the promise of providing long-sought insights into vertebrate evolution and the molecular mechanisms underlying vertebrate myoblast fusion. Here, we report the identification and characterization of MymX and MymK orthologs from lamprey and the tunicate *Ciona robusta,* a non-vertebrate chordate. Despite low sequence similarity, these distantly related proteins can replace the function of their mammalian and reptilian orthologs in driving myoblast fusion. Furthermore, we demonstrate that the fusogenic activity of MymK likely evolved in the last common ancestor of tunicates and vertebrates (Olfactores), and therefore predates the origin of MymX, which appears to have evolved *de novo* specifically in the vertebrate lineage. Unexpectedly, we find evidence that MymK has undergone extensive functional co-evolution with MymX and other, as of yet unknown factors. We propose that vertebrate-specific co-evolution of MymK and MymX has afforded a highly efficient mechanism for myoblast fusion that drives the massive multinucleation of skeletal muscle cells observed in vertebrates. These results present the first definitive genetic evidence for the evolutionary underpinnings of a chordate-specific mechanism for myoblast fusion.

## Results

### Evolutionary origins of MymK

The phylum Chordata is comprised of vertebrates together with two non-vertebrate subphyla: Tunicata and Cephalochordata (**Fig. 1a**). Importantly, each of these extant groups occupies a unique position with regard to the evolution of myoblast fusion and multinucleated muscles: lancelets (cephalochordates) have mononucleated muscles indicating no myoblast fusion (Extended Data Fig. **1a**)^16^, whereas tunicates exhibit limited multinucleation of certain muscles^17^ and vertebrates have extensive, obligatory multinucleation (Extended Data Fig. **1b**). Therefore, comparative gene function studies of these closely related animal groups might shed insights into the evolutionary history and cellular mechanisms of myoblast fusion.

**Figure 1:**
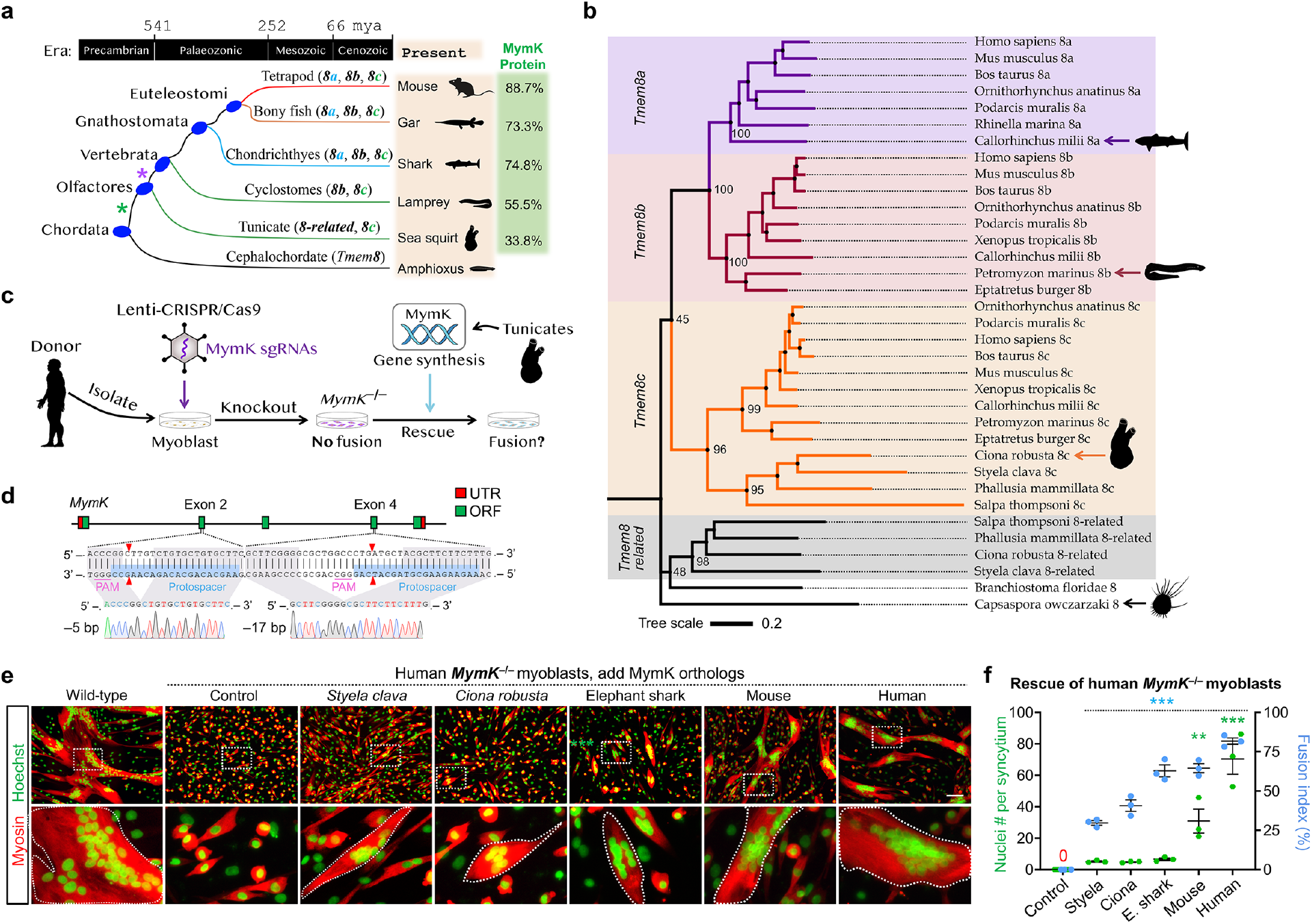
Tunicate MymK orthologs possess weak fusogenic function in human myoblasts. **a**, Phylogenetic relationships of various chordate clades used to deduce the evolutionary origins of the *MymK* gene (also known as *Tmem8c*). Asterisks represent two potential duplication events of *Tmem8* genes that give rise to *8-related*, *8a*, *8b* and *8c* members. **b**, Phylogeny of the Tmem8 gene family inferred by a distance-based method (neighbour joining). The bootstrap percentages obtained from 1,000 replicates were shown for four major clades in the cladogram. Extended phylogenetic analysis is seen in Extended Data. Fig. 3a. **c**, Schematic of experimental design to test the fusogenic function of tunicate MymK proteins in human *MymK*-deficient myoblasts. **d**, Human *MymK* gene structure, sgRNA positions and genotyping results that showed biallelic frameshift mutations induced by CRISPR/Cas9. **e**, Myosin immunostaining of human *MymK*^−/−^ myoblasts transfected with MymK orthologs. Muscle syncytia (outlined) were observed in *Styela* and *Ciona* MymK expression groups, though smaller than the syncytia induced by vertebrate MymK proteins. Scale bar, 100 µm. **f**, Measurements of myoblast fusion in **e** after 4 days of myogenic differentiation. E. shark: elephant shark. Data are means ± SEM. ** *P* < 0.01; *** *P* < 0.001, compared to control group, one-way ANOVA.

Originally known as *Tmem8c*, *MymK* belongs to a gene family that in vertebrates also contains two other paralogs^8^: *Tmem8a* and *Tmem8b*. The evolutionary origins of *MymK* within this gene family are poorly understood. Homology guided searches revealed that multiple tunicate species possess both a *Tmem8c* (*MymK*) gene and a *Tmem8a/b-*like gene (herein named *Tmem8-related*) (**Fig. 1a**; Extended Data Fig. 2a). In the cephalochordate amphioxus (e.g. *Branchiostoma floridae*), only a single *Tmem8* family gene can be identified (**Fig. 1a**). In fact, *Tmem8* sequences are found in diverse eukaryotes, including insects, worms, plants, fungi, and the unicellular filasterean *Capsaspora owczarzaki* (**Fig. 1b**; Extended Data Fig. **3a**). Like vertebrate Tmem8a and Tmem8b, most invertebrate and unicellular Tmem8 orthologs contain an EGF (epidermal growth factor-like) domain that is lacking from vertebrate and tunicate MymK (Extended Data Fig. **2b**). *Tmem8* orthologs are not found outside eukaryotes. Finally, although multinucleation is also a prominent feature of arthropod musculature^18, 19^, the *MymK* gene was not found in this phylum (Extended Data Fig. **3a**).

Phylogenetic analysis suggested that duplication of an ancestral *Tmem8* gene gave rise to *Tmem8a/b* and *MymK* before tunicates and vertebrates diverged. After vertebrates split from tunicates, *Tmem8a/b* underwent gene duplication again giving rise to *Tmem8a* and *Tmem8b*. Whereas most extant vertebrates retained all three Tmem8 family members, cyclostomes (lampreys and hagfishes) appeared to have lost the *Tmem8a* gene (**Fig. 1b**). The exact timing of the duplication event that gave rise to *MymK* and *Tmem8a/b* remains unknown, because phylogenetic analyses were not able to confidently place cephalochordate or tunicate *Tmem8-*related genes (Extended Data Fig. **3a**). Thus, the duplication may have occurred after Olfactores (tunicates + vertebrates) and Cephalochordata diverged, or cephalochordates may have lost *MymK* (Extended Data Fig. **3b**). In either scenario, the lack of multinucleated muscles in cephalochordates suggested a functional link between myoblast fusion and the presence of *MymK* in olfactorians.

### Functional comparisons of tunicate versus vertebrate MymK proteins

We identified *MymK* orthologs in several tunicate species, including the benthic ascidians (e.g. *Styela clava*, *Ciona robusta*) and the pelagic thaliaceans (e.g*. Salpa thompsoni*, *Pyrosomella verticillata*). Tunicate MymK sequences show ∼26–38% amino-acid identity with human MymK (**Fig. 1a**; Extended Data Fig. **2a**). Moreover, tunicate MymK proteins are predicted to have a similar protein domain topology as mammalian MymK (Extended Data Fig. **2b**).

To examine the functional conservation of tunicate MymK as a fusogen, we devised a complementation approach whereby the human *MymK* gene was replaced by the expression of various *MymK* orthologs from different tunicates (**Fig. 1c**). Genotyping of CRISPR–treated human myoblasts revealed biallelic frameshift mutations in *MymK* (**Fig. 1d**), which completely abolished syncytializations (**Fig. 1e**). We then expressed tunicate MymK proteins in these cells and detected a band of predicted size in the membrane compartments as confirmed by Western blot (Extended Data Fig. **2c, d**). Interestingly, tunicate MymK proteins, from either ascidians (**Fig. 1e, f**) or thaliaceans (Extended Data Fig. **4a, b**), consistently rescued the fusion of human *MymK*^−/−^ myoblasts, albeit with lower levels of efficiency than vertebrate proteins. Consistent with the neofunctionalization of *MymK*, tunicate Tmem8-related and cephalochordate Tmem8 proteins did not elicit any fusogenic activity (Extended Data Fig. **4c, d**). We also observed fusogenic activity of tunicate MymK proteins in mouse and brown anole (lizard) *MymK*^−/−^ myoblasts that we generated by CRISPR mutagenesis, indicating broad compatibility with a variety of vertebrate species (Extended Data Fig. **5**).

Human myoblast fusion involves MymK and its function catalyzer MymX^12^. Although MymK protein alone can moderately induce myoblast fusion, co-expression with MymX, as seen during normal myogenesis, enhances fusion^12^. Despite extensive searching, we were not able to identify MymX homologs in tunicates or any other non-vertebrate species. We postulated that fusogenic activity of tunicate MymK is independent of MymX. To test this idea, we generated human *MymX* and *MymK* double knockout (dKO) myoblasts by CRISPR mutagenesis. Indeed, without MymX, tunicate MymK still induced myoblast fusion at a level roughly comparable to human MymK, supporting its conserved function (Extended Data Fig. **6**). However, a functional difference between human and tunicate MymK was unmasked by resupplying MymX protein. Specifically, co-expression of human MymX and human MymK induced massive fusion (Extended Data Fig. **6**). Such synergy was not observed when human MymX was paired with tunicate MymK (Extended Data Fig. **6**). These results suggest that, although fusogenic capability of MymK predates the emergence of vertebrates and the MymX gene, vertebrate-specific changes to MymK were essential for the evolution of functional synergy with MymX.

### Temporally and spatially restricted expression of *MymK* drives multinucleation program of *Ciona* **muscle**

Having established the fusogenic activity of tunicate MymK in vertebrate cells, we next asked whether this fusogen plays a role in the development of multinucleated myofibers in tunicates as well. The presence of *MymK* in tunicates was intriguing because these non-vertebrate chordates also have multinucleated muscles that, like vertebrate skeletal muscles, formed by myoblast fusion^20^. While the tail muscles from tunicate larvae are mononucleated, the siphon and body-wall muscles of post-metamorphic juveniles and adults are formed by a series of multinucleated fibers^21, 22^ (**Fig. 2a**). Moreover, within the tunicate clade, the presence of multinucleated siphon muscles correlates with presence of the *MymK* gene as well, as both were secondarily lost from a unique tunicate group, the neotenic Appendicularians^23^ (**Fig. 2b**).

**Figure 2:**
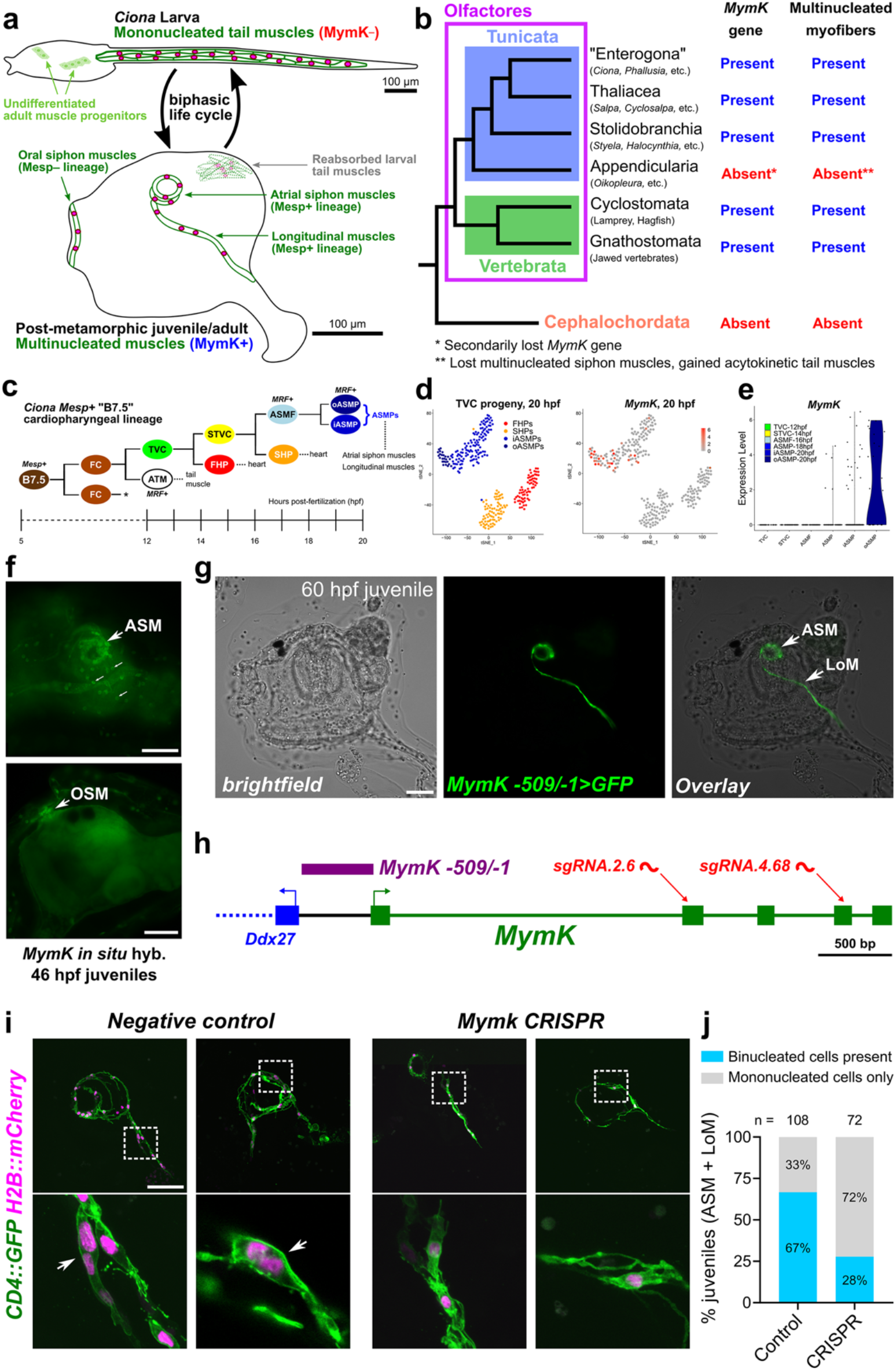
MymK is required for multinucleation of post-metamorphic muscles in the tunicate *Ciona*. **a**, Diagram of biphasic life cycle of ascidians (sessile tunicates) like *Ciona.* The motile larvae have strictly mononucleated tail muscles during the dispersal phase. After settlement and metamorphosis, tail muscle cells undergo programmed cell death and are reabsorbed, while dedicated muscle progenitors set aside in the larva differentiate to form the multinucleated siphon and body-wall muscles of the juvenile. Muscles surrounding and emanating from the oral and atrial siphons are derived from distinct cell lineages in the larva. Only those from the atrial siphon are derived from the Mesp+ B7.5 lineage (in **c**). **b**, Cladogram of extant chordates showing correlation between presence of *MymK* gene and muscle multinucleation in different clades. **c**, Diagram of the B7.5 lineage in *Ciona robusta,* adapted from Razy-Krajka et al^27^. FC: Founder Cell, TVC: Trunk Ventral Cell, ATM: Anterior Tail Muscle Cell, STVC: Secondary TVC, FHP: First Heart Precursor, SHP: Second Heart Precursor, ASMF: Atrial Siphon Muscle Founder Cell, ASMP: Atrial Siphon Muscle Precursor, oASMP: Outer ASMP, iASMP: Inner ASMP. Asterisk indicates that both Founder Cells give rise to identical lineages. *MRF: Myogenic Regulatory Factor* (*MyoD* ortholog). Hpf: hours post-fertilization. **d**, tSNE plots adapted from Wang et al^25^. showing *MymK* expression mapped onto TVC progeny clusters at 20 hpf. *MymK* is expressed exclusively in ASMPs, and especially enriched in Outer ASMPs. Abbreviations same in **c**. **e**, Violin plot comparing *MymK* expression levels in selected TVC derivatives (see panel **c**) at different developmental stages, showing initial infrequent expression starting in ASMFs, increasing in ASMPs at 18 hpf, and enriched primarily in oASMPs at 20 hpf. See Extended Data Fig. 7b for corresponding pseudotemporal expression profile plot. **f**, Whole-mount mRNA *in situ* hybridization showing *MymK* expression in developing atrial siphon muscle (ASM) and oral siphon muscle (OSM) cells in metamorphosing juveniles. Small arrows indicate autofluorescent tunic cells. **g**, Post-metamorphic *C. robusta* juvenile developed from a zygote transfected with a *MymK -509/-1>GFP* reporter plasmid, labeling ASMs and longitudinal body-wall muscles (LoM). **h**, Diagram of *MymK* locus in *C. robusta,* showing location of the *MymK -509/-1* promoter fragment and target sites of single guide RNAs (sgRNAs) used for CRISPR mutagenesis. **i**, Representative Z-projection confocal fluorescence images of 84 hpf negative control (transfected with *Mesp>Cas9* only, no sgRNAs) juveniles alongside same-age juveniles in which *MymK* was targeted for mutagenesis specifically in the B7.5 lineage. *MymK CRISPR:* zygotes transfected with *Mesp>Cas9* and *U6*>*MymK-*sgRNA vectors. Muscle plasma membranes and nuclei labeled by *MRF>CD4::GFP* and *MRF>H2B::mCherry*, respectively. Arrows in negative control panels showing development of typical binucleated myofibers that is inhibited upon *MymK* CRISPR. **j**, Data from scoring of juveniles represented in panel **i** showing reduced frequency of binucleated atrial siphon/longitudinal myofibers in *MymK CRISPR* juveniles. N, numbers of juveniles assayed for each condition. Scale bars, 50 µm.

Previous bioinformatic analyses of myogenic gene expression in the model tunicate *Ciona robusta* had missed *MymK* due to its absence from the most commonly used gene set^24^. By re-analyzing raw single-cell transcriptome data^25, 26^ at different developmental stages of *C. robusta*, we detected *MymK* reads specifically in siphon muscle precursor cells (**Fig. 2c–e**; Extended Data Fig. **7b**) but not mononucleated tail muscle cells from larvae (Extended Data Fig. **7a**). This was confirmed by whole mount *in situ* hybridization (**Fig. 2f**), and by transfection of a *MymK* promoter reporter plasmid (*MymK -509/-1>GFP*) that exclusively labeled multinucleated juvenile muscles (**Fig. 2g**; Extended Data Fig. **7c, d**), but not in any other cell type including mononucleated larval tail muscle cells. Taken together, these results suggest that expression of the *MymK* gene is tightly correlated with muscle multinucleation in tunicates.

To test the requirement of *MymK* for multinucleation of tunicate siphon muscles, we performed *MymK* loss-of-function experiments in *Ciona.* We inactivated the *MymK* gene by performing tissue-specific CRISPR/Cas9-mediated mutagenesis^27^ in the cardiopharyngeal mesoderm lineage (Mesp+, **Fig. 2c**) that gives rise to the atrial siphon and associated longitudinal muscles of the *Ciona* juvenile^28^. Animals were transfected with *Mesp>Cas9* and validated single guide RNA (sgRNA) expression vectors to induce double-stranded breaks in the 2^nd^ and 3^rd^ exons of *MymK* (**Fig. 2h**; Extended Data Fig. **8a**) specifically in Mesp+ progenitors. In control juveniles transfected with *Mesp>Cas9* alone, circular atrial siphon myofibers invariably formed as orderly rings with occasional longitudinal myofibers emanating from the siphon region (**Fig. 2i**, Extended data Fig. **8c**; Supplementary Video **1**). In contrast, *MymK* CRISPR resulted in highly disorganized atrial siphon muscles (**Fig. 2i**; Extended Data Fig. **8b, d**; Supplementary Video **2**), while oral siphon myofibers (which do not express *Mesp>Cas9*) from the same animals were still formed normally (Extended data Fig. **8d**). Moreover, there was a reduction in the frequency of binucleated atrial siphon/longitudinal myofibers in *MymK* CRISPR juveniles (**Fig. 2i, j**), suggesting that *MymK* is required for myoblast fusion in *Ciona.* We also tested whether *MymK* is sufficient to promote fusion of normally mononucleated muscle cells of the larval tail, by overexpressing it in tail muscle progenitors using the *Myogenic Regulatory Factor* (*MRF,* also known as *MyoD*) promoter. Although cell morphology was altered, we did not detect the clear presence of multinucleation (Extended Data Fig. **9**; 16 hpf control, Supplementary Video **3**; 16 hpf *MRF*>*MymK*, Supplementary Video **4**). This suggests that, as in vertebrates^12^, MymK activity in *Ciona* likely requires other factor(s), which in *Ciona* should be expressed in juvenile but not larval tail muscle cells.

### A distantly related MymX sequence from lamprey genomes

Lampreys are descended from an ancient cyclostome vertebrate lineage that diverged from jawed vertebrates (gnathostomes) ∼500 million years ago^29^. Histological analysis revealed extensive multinucleation of lamprey muscle cells (**Fig. 3a**), which can host up to several hundred myonuclei per fiber, a level of fusion comparable to those in jawed vertebrates including shark (Extended Data Fig. **1b**). We hypothesized that a protein with MymX function exists in lamprey to robustly induce myoblast fusion in cooperation with MymK.

**Figure 3:**
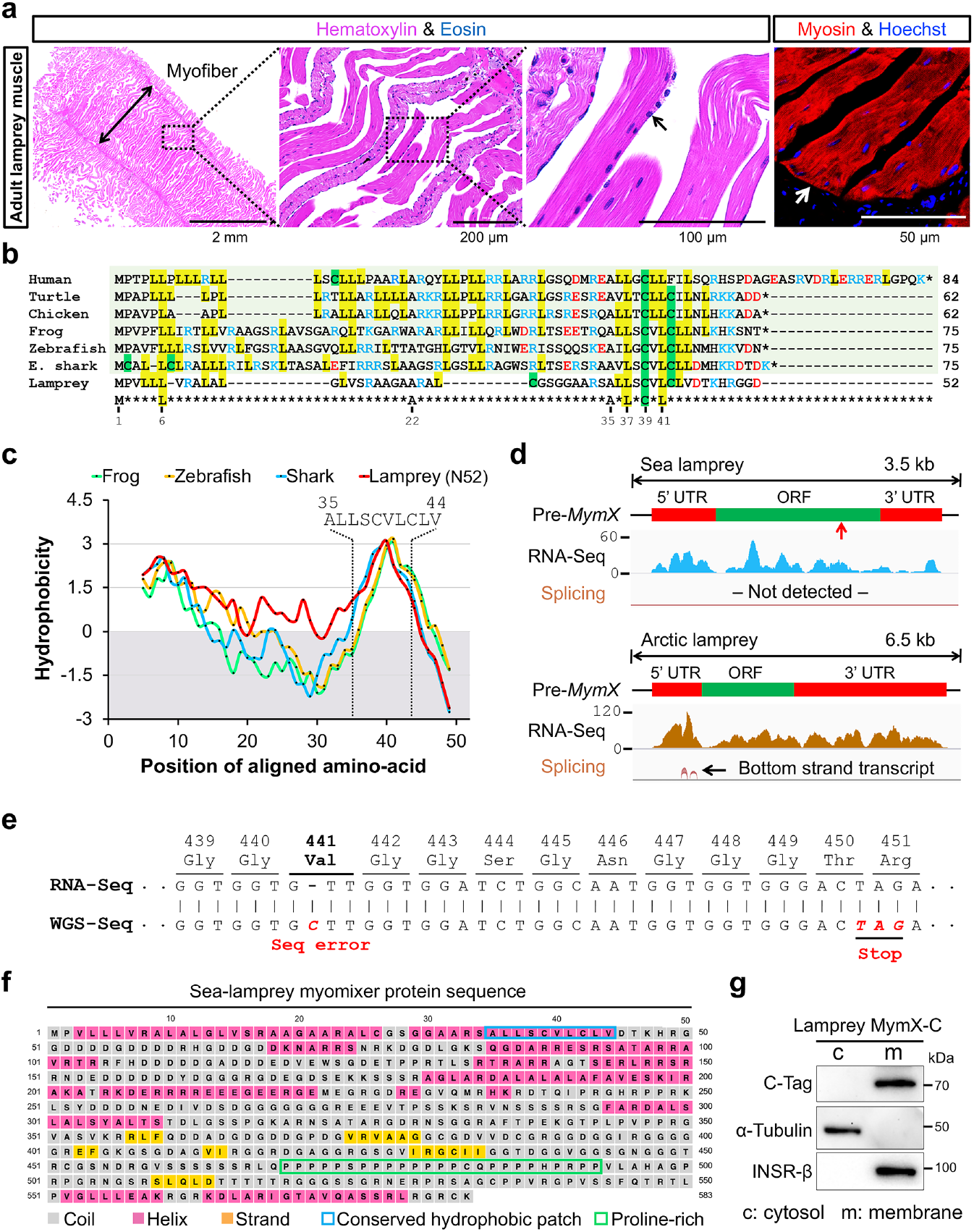
Discovery of the unusual *MymX* genes from lampreys. **a**, Histological staining and immunofluorescence of muscle tissues dissected from adult sea lamprey (*Petromyzon marinus*). Multinucleated myofibers (arrows) are observed from the longitudinal sections. **b**, Cross-species homology of lamprey MymX aligned with its orthologs from jawed vertebrates. Only a few residues from the AxLyCxL motif can be aligned. x denotes leucine, valine, or isoleucine, and y denotes serine, threonine, or glycine. The numbers below the consensus sequence refer to the positions in sea lamprey MymX (only the N-terminal 52 amino acids are shown). **c**, Hydrophobicity signatures of MymX proteins for the aligned regions in **b**. The conserved hydrophobic patch (Ala35–Val44) from lamprey MymX is indicated. **d**, RNA sequencing tracks that confirmed transcription of *MymX* genes in embryos of two closely related lamprey species. SRA accession: PRJNA497902 for sea lamprey and PRJNA371391 for arctic lamprey (*Lethenteron camtschaticum*). No splicing junction was detected in these hypothetical ORFs. **e**, Alignment of sequencing (seq) reads of RNA (SRA accession: PRJNA50489) with genome data (GenBank: AEFG01021847). A single-nucleotide gap of a cytosine insertion is highlighted (red). Cloning and sequencing of this region validated the correct reading frame shown in RNA-seq reads. WGS: whole genome shotgun. **f**, Prediction of the secondary structure for full-length sea lamprey MymX (583 amino acids). **g**, Western blot analyses of cytosolic (c) and membrane (m) fractions of human myoblasts transfected with C-tagged lamprey MymX. α-Tubulin blot was used as a positive control of cytosolic proteins. Insulin receptor β (INSR-β) blot was used as a positive control of membrane proteins.

The search for MymX orthologs is intrinsically challenging due to the small size of MymX proteins (< 100 residues) and a high frequency of amino-acid substitutions along its entire sequence (**Fig. 3b**). Therefore, we iteratively BLAST searched a large pool of lamprey genome and transcriptome data for a sequence related to any known MymX proteins. These attempts were unsuccessful even when the search algorithm threshold was relaxed (E value 100). However, using the short C-terminus region of elephant shark (*Callorhinchus milii*) MymX, one hit was discovered from the whole-genome shotgun (WGS) sequence (GenBank: AEFG01021847.1) of the sea lamprey (*Petromyzon marinus*). This distantly related sequence only aligned with a few hydrophobic residues that constituted an AxLyCxL motif (**Fig. 3b**), which is essential for mammalian MymX function^30^. Intriguingly, despite its highly diverged sequence, one part of this lamprey MymX-related sequence shared a hydrophobicity signature with known MymX proteins (**Fig. 3c**), hinting at structural and functional conservation.

Next, we set out to annotate this gene by defining its expression, exon structure and coding sequences (**Fig. 3d**). Sequence discrepancy of RNA-seq and WGS reads was noted in one site of the ORF (**Fig. 3e**). Targeted sequencing of this region identified a complete ORF that encodes 583 amino acids (**Fig. 3f**), a number that is almost ten times larger than known MymX proteins, e.g. 60 amino acids from reptiles (e.g. *Anolis carolinensis*). The extended length of lamprey MymX appears to be due primarily to its C-terminus, which contains largely unstructured and repetitive sequences (**Fig. 3f**). A homologous sequence was also discovered from arctic lamprey (*Lethenteron camtschaticum*, APJL01015224) that is composed of 595 amino acids (Extended Data Fig. **10**), sharing 93% identity with sea lamprey MymX. The complete ORF of sea lamprey MymX was codon-optimized, cloned by gene synthesis and expressed in human myoblasts. Western blot readily detected a 70 kDa band specifically from the membrane fraction (**Fig. 3g**), suggesting a function in this compartment.

### MymX protein from lamprey can replace its mammalian orthologs in enhancing myoblast fusion

We investigated the function of sea lamprey MymX in promoting myoblast fusion using a similar rescue experiment as for MymK above. Of note, human myoblasts with deletion of *MymX* are weakly fusogenic, due to the activity of MymK in these cells^12^. Strikingly, the expression of lamprey MymX in human (**Fig. 4a**) and mouse *MymX*^−/−^ myoblasts (Extended Data Fig. **11**) rescued their fusion deficiency. The large human muscle syncytia induced by lamprey MymX contained an average of 17 nuclei, a stark contrast to 4 nuclei in the control group (**Fig. 4b**). Using the same assay, even larger myotubes were formed when expressing shark, zebrafish or human MymX (**Fig. 4a, b**), indicating higher activity of more closely related orthologs, as expected. Similar to human MymX, lamprey MymX strictly required human MymK for function because it failed to induce fusion when *MymK* was deleted from human myoblasts (Extended Data Fig. **12**). Therefore, this distantly related lamprey sequence is a functional and authentic ortholog of MymX and capable of synergizing with human MymK to promote myoblast fusion, in spite of sequence divergence and a many-fold difference in protein length (**Fig. 4c**). Because MymX is not found in any non-vertebrate chordate groups and does not share homology with any other proteins, we conclude that *MymX* is a vertebrate-specific orphan gene encoding a core molecular component of myoblast fusion that is conserved from lampreys to humans.

**Figure 4:**
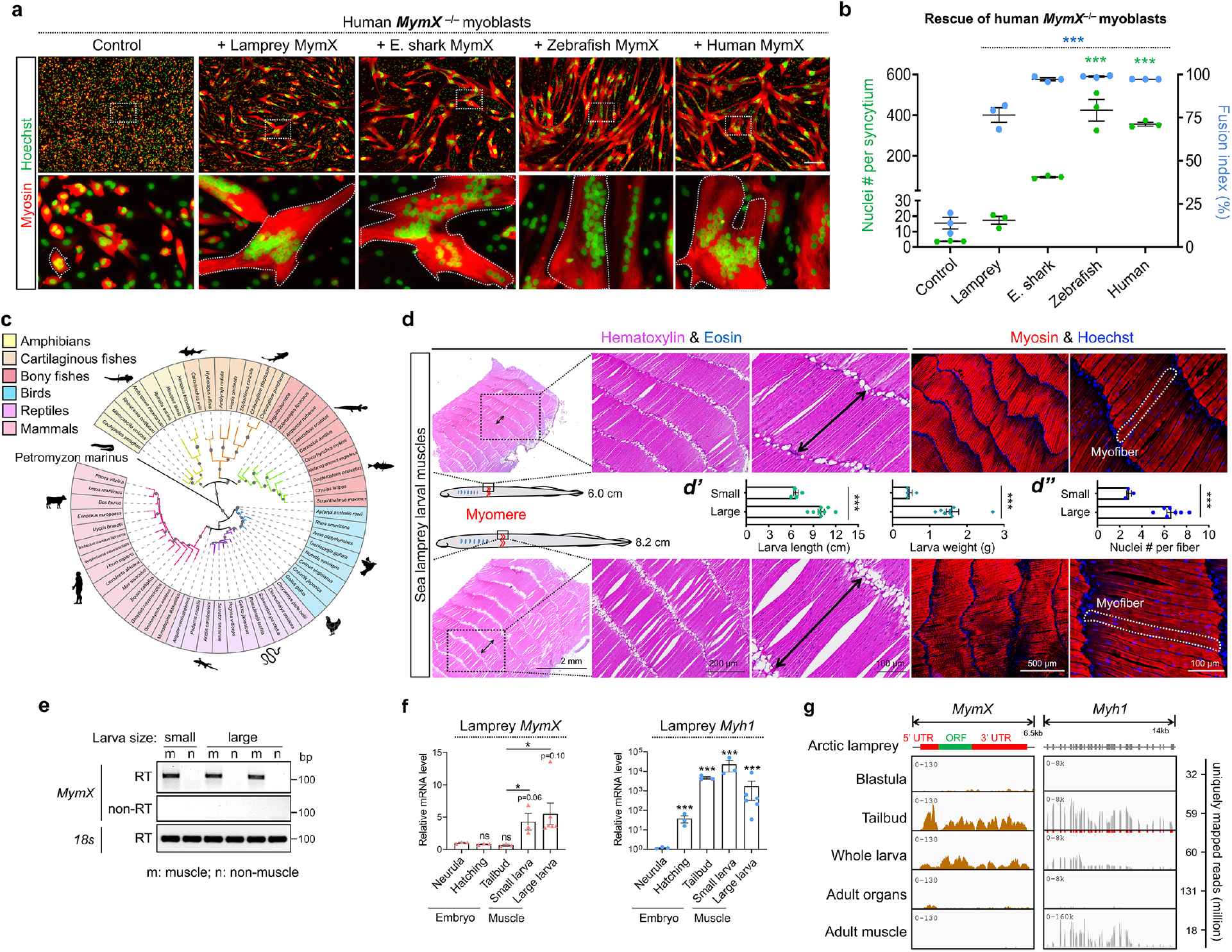
Lamprey *MymX* is specifically expressed in fusing larval muscles and can replace its human ortholog in driving myoblast fusion. **a**, Myosin immunostaining of human *MymX* ^−/−^ myoblasts transfected with MymX orthologs. Note that sea lamprey MymX can rescue fusogenic defects of human *MymX* ^−/−^ cells that formed larger muscle syncytia (outlined) than control (empty vector). E. shark: elephant shark. Scale bar, 100 µm. **b**, Measurement of myoblast fusion in **a** after 4 days of differentiation. **c**, Phylogenetic tree of *MymX* orthologs. *MymX* genes are only found in vertebrate species. **d**, Staining of longitudinal sections of muscle tissues dissected from sea lamprey larvae. Larvae of two different size-groups were analyzed to identify muscle fusion stage. Measurements of lamprey body length and weight (***d’***), and nuclei number per myofiber (***d’’***) are presented. Considering adult lamprey myofibers contain much larger numbers of myonuclei, these larvae represent an early myonuclei accretion stage. **e**, Reverse transcription PCR results that validated the muscle-specific expression pattern of *MymX* gene in sea lamprey larval muscle tissues. m, muscle cDNA; n, non-muscle (intestine and liver) cDNA. **f**, qPCR results that measured the expression levels of sea lamprey *MymX* and myosin heavy chain 1 (*Myh1*) genes at various embryonic development and larval growth stages. Larva sizes refer to the grouping in **d**. **g**, RNA-seq results of *MymX* and *Myh1* genes from arctic lamprey (*Lethenteron camtschaticum*) embryos (SRA accession: PRJNA371391), whole larva (17-day old, SRA accession: PRJNA553689), adult organs (notochord, ovary, testis, kidney and heart, SRA accession: PRJNA354821) and muscle tissues (SRA accession: PRJNA354821). Note that *MymX* gene in arctic lamprey is specifically expressed during muscle development and growing stages. Data are means ± SEM. * *P* < 0.05, *** *P* < 0.001. Panel b: compared to control group, one-way ANOVA; panel f: Student’s *t* test.

### Expression of *MymX* coincides with lamprey muscle multinucleation *in vivo*

We next investigated the expression pattern of MymX during lamprey muscle development. Sea lampreys have a complex life cycle that involves a larval period of 2–10 years, followed by metamorphosis into an adult stage. It is unclear when myoblast fusion occurs in lampreys, though it was reported that muscle cells from young larvae remained mononucleated^31^. To identify the temporal window of active myoblast fusion for sea lamprey, we examined two larval groups of distinct sizes that are estimated of 2–3.5 years of age (**Fig. 4d**), considering that multinucleation is a major cellular mechanism of muscle growth^32, 33^. Indeed, moderate multinucleations were consistently observed in both groups (**Fig. 4d**), while an increase of nuclei number per myofiber was associated with a larger size of muscle and animal (**Fig. 4d’, d’’**). Consistent with a role for MymX in sea lamprey myogenesis, its expression was readily detected in larval muscles but not other tissues (**Fig. 4e**). Although *MymX* was also expressed in sea lamprey embryos when myoblast fusion does not occur, its expression was significantly upregulated in larval muscle tissues (**Fig. 4f**). Expression of the *MymX* gene in arctic lamprey was also confirmed by RNA-seq data (**Fig. 4g**). Thus, lamprey *MymX* is a muscle-specific gene and its abundant expression coincides with syncytialization of lamprey myofibers.

### Lamprey-specific carboxyl terminus from MymX is indispensable for optimal fusogenic activity

For lamprey MymX, only a short region of 52 amino acids at the N-terminus (N52) can be aligned to conventional orthologs (**Fig. 3b**). However, expression of the N52 polypeptide failed to induce myoblast fusion (**Fig. 5a–c**). A similarly deleterious effect was observed when the conserved AxLyCxL motif was removed (**Fig. 5a–c**). We continued to dissect the function of the non-conserved yet long C-terminus sequence by generating a series of sea lamprey *MymX* mutants (**Fig. 5a**). Interestingly, as the region of deletions enlarged, MymX function, quantified as nuclei number per syncytium, gradually diminished (**Fig. 5c**; Extended Data Fig. **13a, b**). Therefore, the optimal activity of sea lamprey MymX requires its large C-terminal structure.

**Figure 5:**
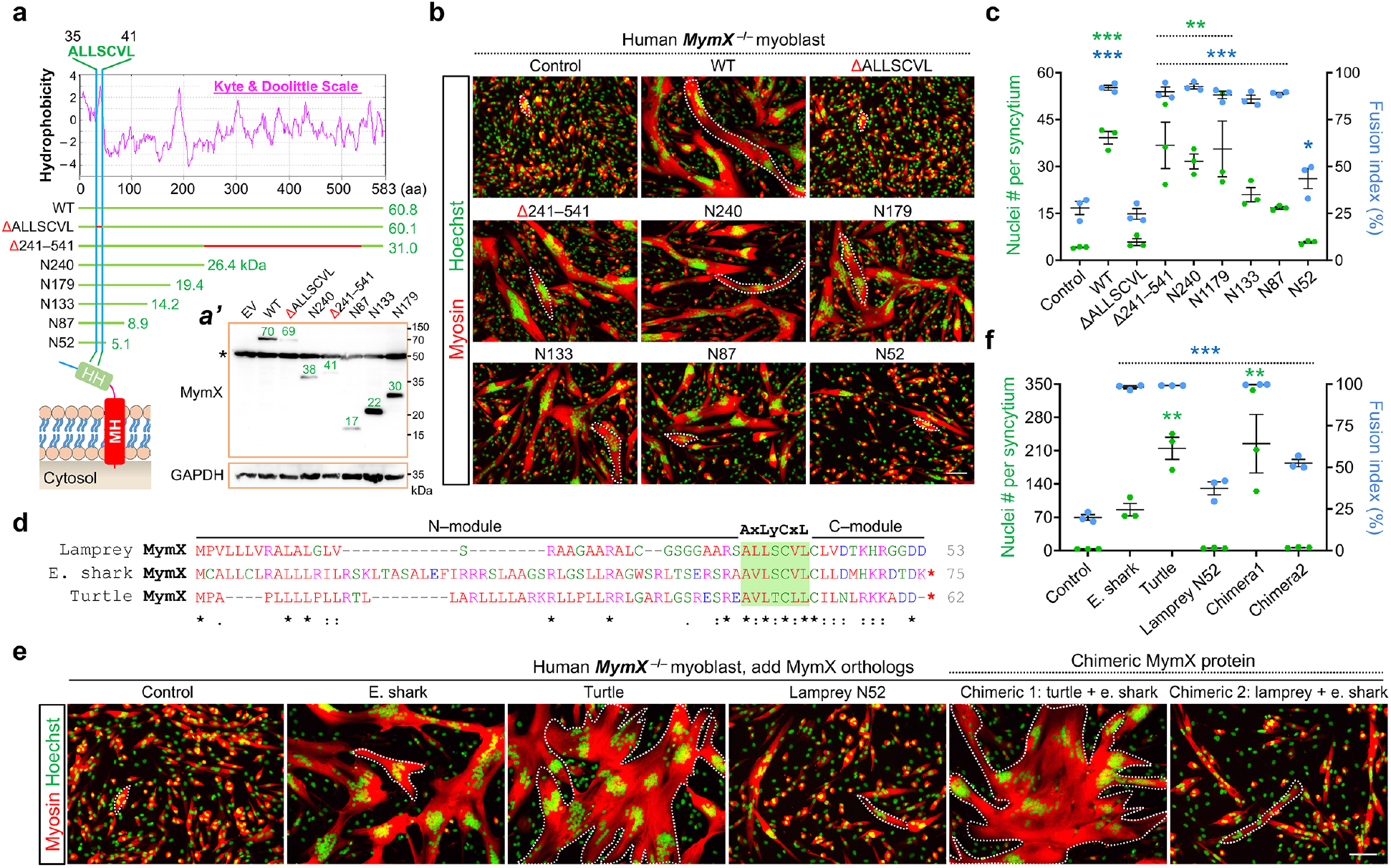
Structure–function analysis of lamprey MymX protein. **a**, Hydrophobicity map of sea lamprey MymX and a schematic of mutants. Red lines highlight deleted regions. HH, hydrophobic helix; MH, membrane-anchor helix. ***a’***, Western blot results that confirmed expression of lamprey MymX mutants in human myoblasts. MymX was detected by blotting a diminutive C-tag fused at the C-terminus of target. The predicted and detected molecular weights are labelled on the schematics and Western blots, respectively. The 4-amino-acid epitope tag (EPEA) is 0.4 kDa. Replicates are shown in Extended Data Fig. 13b. **b**, Myosin immunostaining of human *MymX* ^−/−^ myoblasts transfected with full length (WT) or truncated lamprey MymX proteins. The size of myotubes (outlined) gradually reduced as the length of C terminus shortens. **c**, Measurement of myoblast fusion in **b** after 4 days of myogenic differentiation. **d**, Amino acid sequences of MymX and cross-species homology that guides the design of chimeric proteins shown in following panels. E. shark: elephant shark. **e**, Myosin immunostaining of *MymX* ^−/−^ myoblasts transfected with MymX orthologs or chimeric proteins. Multinucleated myotubes are outlined. The first chimera comprised N-terminal region of turtle MymX and C-terminal region of elephant shark MymX in **d**; the second chimera comprised N-terminal region of lamprey MymX with C-terminal region of elephant shark MymX in **d**. **f**, Measurement of myoblast fusion in **e** after 4 days of myogenic differentiation. Scale bars, 100 µm. Data are means ± SEM. * *P* < 0.05, ** *P* < 0.01, *** *P* < 0.001, compared to control group, one-way ANOVA.

In parallel, we also tested whether the C-terminus of sea lamprey MymX can be functionally replaced by a shorter version from other MymX proteins, e.g. elephant shark (*Callorhinchus milii*) (**Fig. 5d**). Corroborating the modular function of this domain in jawed vertebrates, we show that the C-terminus of turtle MymX (*Chrysemys picta bellii*) can be replaced with that of the elephant shark MymX without losing function (**Fig. 5e, f**; Extended Data Fig. **13c**). However, replacing the lamprey C-tail with that of shark completely abolished lamprey MymX function (**Fig. 5e, f**). These results suggest that the evolutionary adaption of MymX proteins did not simply shorten C-tail length, but also involved major compensatory changes along the entire length of the protein.

### Lamprey MymK is a muscle-specific protein yet does not elicit fusogenic activity in gnathostome cells

Based on the synergy observed between lamprey MymX and human MymK proteins (**Fig. 4a**), we initially expected similar cooperativity of lamprey MymX with its putative endogenous partner, lamprey MymK. Analyses of sea lamprey RNA-seq data revealed the conserved intron/exon structure of *MymK*, which in sea lamprey codes for a protein of 220 amino acids with a 56% identity to human MymK (**Fig. 6a**; Extended Data Fig. **14**). The expression of this ORF in lamprey larval muscles was also confirmed by reverse transcription PCR (**Fig. 6b**) and validated by Sanger sequencing (Extended Data Fig. **15**). Uniquely for the lamprey MymK gene, two additional exons (named as exon 1’ and 1) were discovered that can produce three splicing variants (**Fig. 6a**), two of which encode the same protein while the third can encode 78 additional amino acids (**Fig. 6a**), depending on the start site of transcription (Extended Data Fig. **15a**). By PCR profiling of transcript species, we show that only the short isoform (220 amino acids) is expressed in lamprey larval muscles that show active myoblast fusion (**Fig. 6c**; Extended Data Fig. **15b–e**).

**Figure 6:**
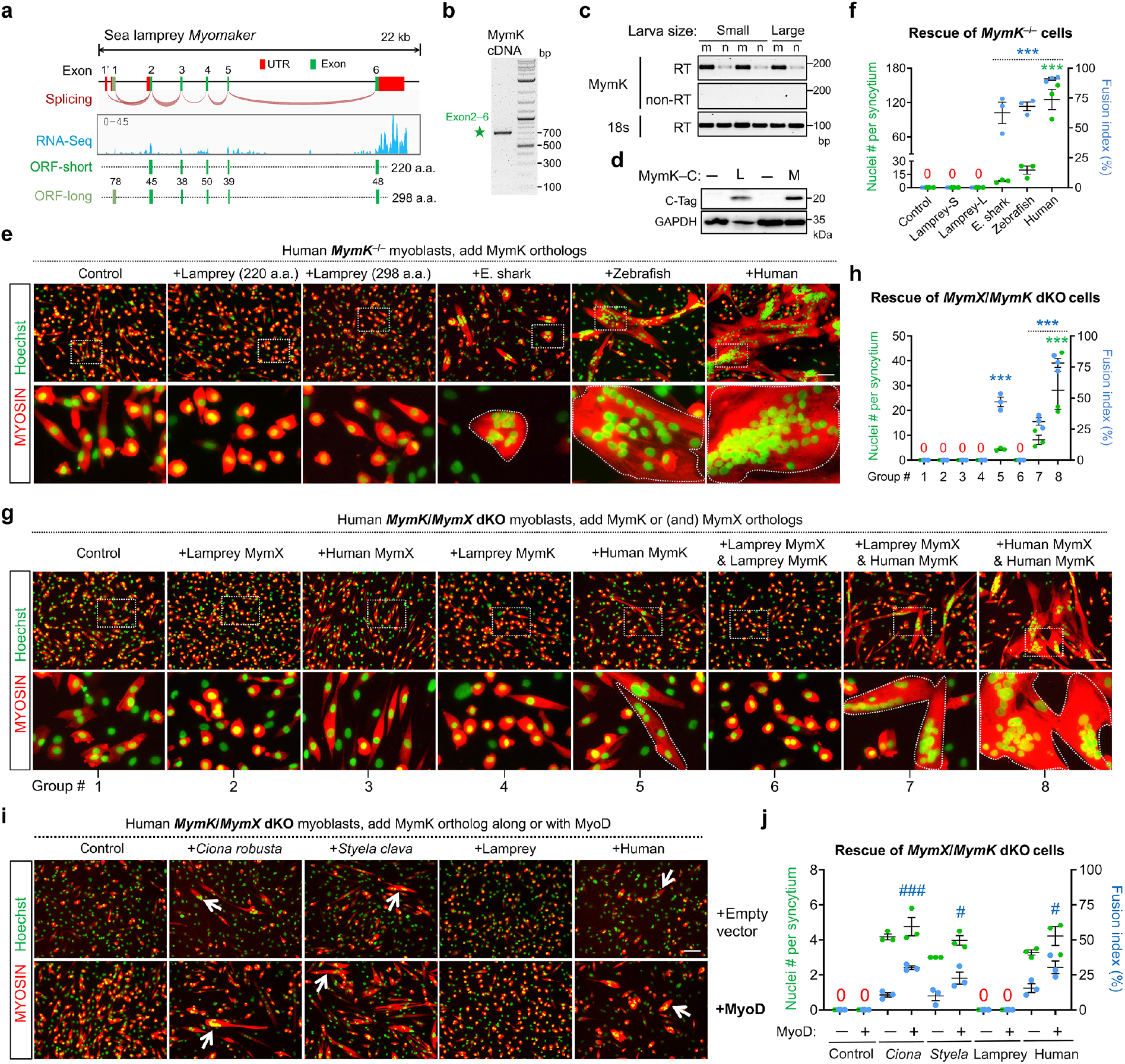
Cyclostome MymK proteins cannot activate fusion of human *MymK* ^−/−^ myoblasts. **a**, Lamprey *MymK* gene structure and RNA sequencing tracks that confirmed its expression in embryos of sea lamprey (SRA accession: PRJNA497902). Note that joining of exon 1 with exons 2–6 can produce a longer isoform that encodes 298 amino acids (a.a.), though this mRNA isoform was not detected by RT-PCR amplification of developing muscle tissues (see Extended Data Fig. 15). **b**, RT-PCR results that confirmed the expression of exons 2–6 in fusion-stage muscle tissues of lamprey larvae. **c**, RT-PCR results that showed muscle-specific expression pattern of *MymK*. m, muscle cDNA; n, non-muscle (intestine and liver) cDNA. **d**, Western blot results that validated the overexpression of C-tagged sea lamprey (L) and mouse (M) MymK proteins in human myoblasts. The predicted molecular weight is 24.8 kDa for lamprey MymK. **e**, Myosin immunostaining of human *MymK* ^−/−^ myoblasts transfected with MymK orthologs. Muscle syncytia are outlined. Note that lamprey MymK, both short and long isoforms, failed to induce human myoblast fusion. E. shark: elephant shark. **f**, Measurement of myoblast fusion in **e** after 5 days of differentiation. S, short isoform; L, long isoform. **g**, Myosin immunostaining of human *MymK*/*MymX* dKO myoblasts that tested the function synergy between MymK and MymX orthologs. Muscle syncytia are outlined. Note that sea lamprey MymK does not possess any fusogenic function in human myoblasts even when sea lamprey MymX was provided. **h**, Measurement of myoblast fusion for groups in **g** after 5 days of differentiation. **i**, **j**, Myosin immunostaining of human *MymK/MymX* dKO myoblasts transfected with MymK orthologs individually (upper panels) or together with mouse MyoD protein (lower panels). Muscle syncytia are pointed by arrows. MyoD promotes myoblast fusion when tunicate or human MymK protein is co-expressed, yet co-expression of MyoD with sea lamprey MymK fails to induce fusion. Scale bars, 100 µm. Data are means ± SEM. ***, ### *P* < 0.001; # *P* < 0.05. * compared to control group, one-way ANOVA; # compared effect of MyoD expression, one-way ANOVA.

To test its potential fusogenic activity, we codon-optimized the lamprey MymK ORF and generated an expression vector by gene synthesis. After transfection into human myoblasts, its expression was confirmed by Western blot that detected a similar size with that of human MymK (**Fig. 6d**). The plasma membrane localization of lamprey MymK protein was also validated by live cell staining using an epitope tag fused to its N-terminus extracellular region (Extended Data Fig. **16**). Surprisingly, lamprey MymK, either short or long isoform, did not elicit any fusogenic activity in either human (**Fig. 6e, f**), mouse (Extended Data Fig. **17a**), or lizard (Extended Data Fig. **17b**) *MymK*^−/−^ cells, which remained mononucleated after full-term myogenic differentiation. As positive controls, elephant shark, zebrafish and human MymK proteins consistently induced fusion of myoblasts and the formation of large myotubes (Extended Data Fig. **5**). In addition to the sea lamprey MymK, we also tested the function of another closely-related ortholog from arctic lamprey (Extended Data Fig. **18a**), which similarly failed to induce fusion of *MymK*^−/−^ myoblasts generated from human (Extended Data Fig. **18b, c**), mouse (Extended Data Fig. **17a**), or lizard (Extended Data Fig. **17b**).

We also identified and tested a third cyclostome MymK from the hagfish (*Eptatretus burgeri*), which shares 67% protein identity with sea lamprey MymK. By histological analysis, we first confirmed the presence of heavily multinucleated myofibers in adult hagfish muscles (Extended Data Fig. **19a**). Interestingly, hagfish MymK also failed to elicit any fusogenic activity in the three jawed vertebrate species myoblasts (Extended Data Fig. **17a, b**; Extended Data Fig. **19b**), though its expression could be detected by Western blot (Extended Data Fig. **19c**). Therefore, despite having high sequence similarity, MymK proteins from jawless vertebrates consistently fail to elicit any fusogenic activity in mammalian or reptilian myoblasts. This was unexpected given that distinct tunicate MymK proteins, which have less sequence similarity, consistently show fusogenic activity in all these cells under the same conditions.

We reasoned that the absence of activity for cyclostome MymK might reflect a strict requirement for cooperation with cyclostome MymX but also an incompatibility with MymX proteins existing in these jawed species myoblasts. Therefore, we co-expressed lamprey MymK and MymX in human *MymK/MymX* dKO cells. However, fusion was still not observed (**Fig. 6g, h**). As positive controls, myotubes were formed when human MymK was expressed alone or together with human or lamprey MymX proteins (**Fig. 6g, h**). Thus, even though lamprey MymX can synergize with mammalian MymK to promote myoblast fusion, it is still unable to elicit any fusogenic activity from lamprey MymK.

Membrane coalescence between two cells requires MymK function in both cells^8^. For instance, exogeneous expression of mammalian MymK in fibroblasts can drive fusion with myoblasts that normally express MymK^8^, even when MymK function is sub-optimal on the fibroblast side^12, 15^. Thus, we tested the activity of lamprey MymK in this heterologous fusion system, in which GFP+ fibroblasts are mixed with mCherry+ myoblasts (Extended Data Fig. **20a**). Consistently, both tunicate and human MymK conferred fusogenic activity to fibroblasts, by which large dual-labelled fibroblast–myoblast syncytia were generated (Extended Data Fig. **20b**). In contrast, lamprey MymK protein still could not induce fusion (Extended Data Fig. **20b**). Recent fusion reconstitution experiments suggested that MymK requires as-of-yet unidentified MyoD-dependent permissive factor(s) to induce myoblast fusion^12^, since MymK expression is not sufficient to induce fusion between *MyoD*^−/−^ myoblasts^12^. Similar results were observed with tunicate MymK proteins in human *MyoD*^−/−^ myoblasts (Extended Data Fig. **21**). We found that prolonged expression of *MyoD* in human myoblasts significantly boosted tunicate and vertebrate MymK function and induced higher level of multinucleations (**Fig. 6i, j**; Extended Data Fig. **22d, e**), suggesting the dose-dependent activity of a rate-limiting factor(s). However, sea lamprey MymK failed to elicit any fusion even with MyoD overexpression. This suggests that, while tunicate and vertebrate MymK proteins likely interact with the same permissive factor(s), sea lamprey MymK does not. In summary, the marked distinction of activity of cyclostome MymK stands in stark contrast to the highly conserved function of MymX proteins, which nonetheless show much greater sequence divergence.

### Structural modeling analyses of MymK evolution

Because the function of a protein depends on its structure, we sought to understand the structural changes of MymK proteins during olfactorian evolution, which might better explain the differential fusogenic activity of tunicate and cyclostome MymK proteins in mammalian or reptilian cells. Deep learning emerged as a valuable approach to predict structures for proteins that would otherwise be difficult to test experimentally^34^. Furthermore, computational modeling also offers unparalleled power for batch analysis. By applying a new deep neural network-based structure assembly method^35, 36^, we obtained nine structural models for MymK proteins from representative species of vertebrates and tunicates (Extended Data Fig. **23a**). Template modelling score (TM-score) is a standard metric to assess the similarity of computational models relative to experimentally determined structures^37, 38^. TM-score ranges between 0 and 1 with >0.5 reflecting a correct global topology^37^, and >0.914 as equivalent to the experimentally determined structure^34^. As the native MymK structure is not available to assess the prediction accuracy, we assessed the accuracy of MymK structural models through a well-established confidence scoring system built on large-scale benchmark tests^35^. Reflecting the high confidence of accuracy, the estimated TM-scores for the MymK models are all around 0.8 (Extended Data Fig. **23a**). Of note, our MymK model shares high similarity with that predicted by AlphaFold2 (AF-A6NI61-F1-model_v1)^39^ and RoseTTAFold^40^, two other neural network-based methods.

MymK proteins from all taxonomic groups share a similar structure, and the similarity score between any MymK models readily exceeds sequence similarity (**Fig. 7a**). All MymK proteins contain seven transmembrane (TM) helices arranged in an anti-clockwise manner from TM1 to TM7 when viewed from extracellular space (**Fig. 7b**). As part of TM1, the N-terminal residues 1–7 protrude toward outside of the cell and form the extracellular face together with three extracellular loops (Extended Data Fig. **23b, c**). The structure of C-terminal residues 202–221 is disordered, which forms the intracellular face together with three intracellular loops (Extended Data Fig. **23b, c**). The TM helices of MymK enclose an internal cavity that goes through the entire structure with a small intracellular opening and a larger extracellular opening (Extended Data Fig. **23d**).

**Figure 7:**
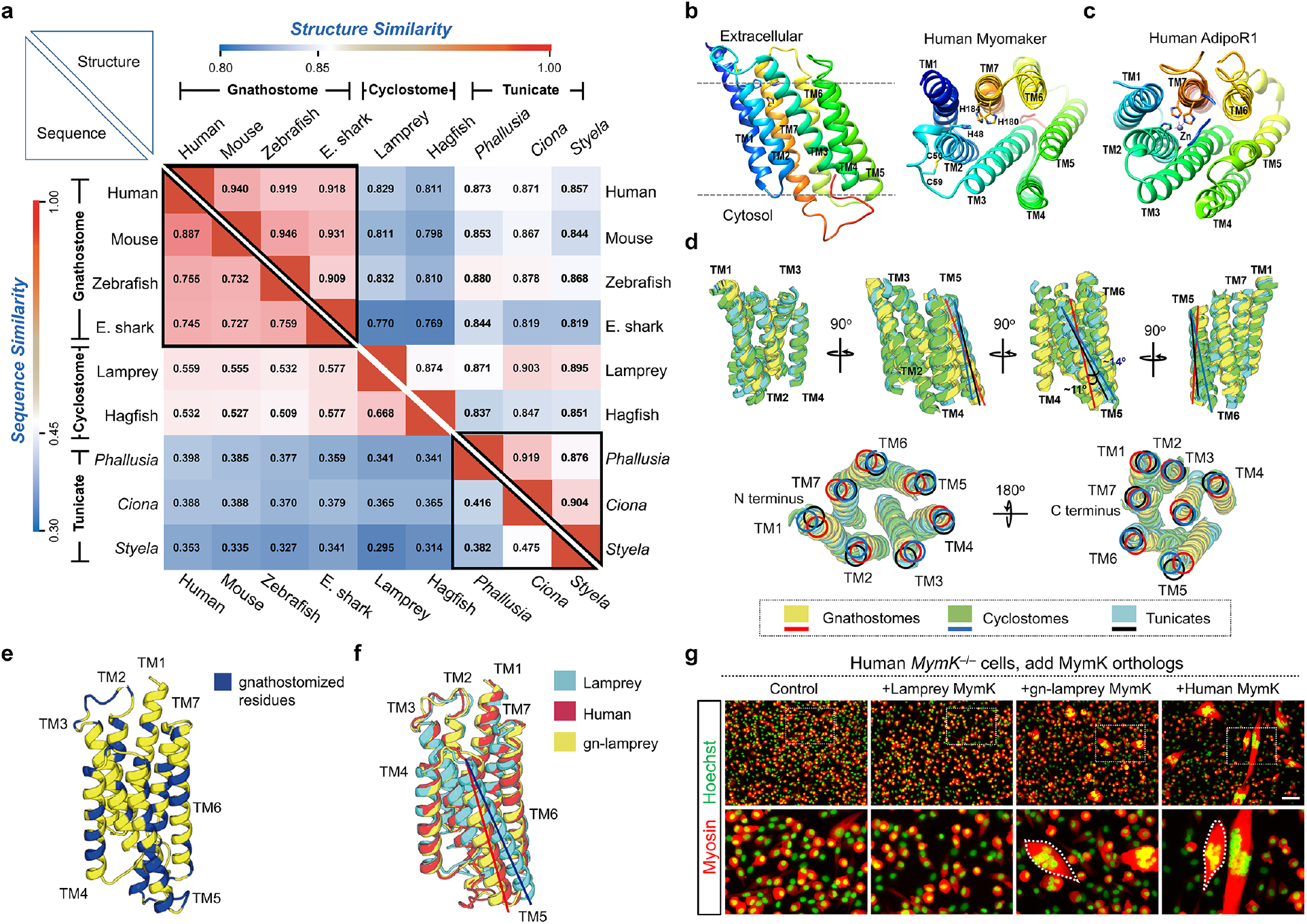
Modeling the structural evolution of MymK proteins. **a**, Similarities of amino-acid sequence (lower left triangle) and predicted structure (upper right triangle) between MymK proteins. **b**, Ribbon representation of the predicted human MymK structure. TM: transmembrane helix. The conserved histidine and cysteine residues on human MymK model are highlighted. **c**, Zinc-binding motif of adiponectin receptor 1 (AdipoR1, PDB ID: 6KRZ). **d**, Superimpositions of the overall structural models for MymK proteins from gnathostomes, cyclostomes and tunicates. The orientations of TM helix 5 show obvious shifts between taxonomic groups. **e**, Positions of mutated residues on gnathostomized lamprey MymK (gn-lamprey MymK). **f**, Superimpositions of MymK structure models. **g**, Myosin immunostaining of human *MymK*^−/−^ myoblasts that revealed the fusogenic activity of gn-lamprey MymK. Cells were differentiated for 4 days. Scale bar, 100 µm. See Extended Data Fig. 23h for quantification results.

MymK models resemble the structure of adiponectin receptor (AdipoR) (**Fig. 7c**; Extended Data Fig. **23e**), though the N-terminus-out topology of MymK is the opposite of the C-terminus-out topology of AdipoR^15, 41^. Stabilization of AdipoR structures requires a zinc ion coordinated by three Histidine (His) residues (**Fig. 7c**). Of note, MymK models contain a similar motif which is located in the outer lipid-layer of the membrane and composed of His48 from TM2, His180 and His184 from TM7 (**Fig. 7b**, **right**). In addition, two cysteine (Cys) residues from the TM2 (Cys50) and extracellular loop 1 (Cys59) are predicted to form a disulfide bond (**Fig. 7b**, **right**). These histidine and cysteine residues are highly conserved in MymK orthologs from all taxonomic groups, suggesting a crucial contribution to the structure and function of MymK proteins.

Unsupervised comparison of the MymK models clustered them into groups that are consistent with their taxonomic identities (**Fig. 7a**). Interestingly, the structural models of gnathostome MymK are more closely related to tunicate MymK than to cyclostome MymK, though the sequence similarity showed an opposite trend (**Fig. 7a**). The major difference in MymK models among taxonomic groups is the orientation of TM5 helix. TM5 in gnathostomes is tilted by 14° and 11° relative to TM5 in cyclostomes and tunicates, respectively (**Fig. 7d**). Thus, perhaps the conservation of fusogenic activity of tunicate MymK in jawed vertebrates might be due to this closer structural similarity, in particular the more subtle shift in TM5 orientation.

To test whether we could elicit fusogenic activity of lamprey MymK in mammalian cells by altering its predicted structure, we “gnathostomized” lamprey MymK by changing those protein residues that are conserved in gnathostomes yet divergent in cyclostomes (**Fig. 7e**; Extended Data Fig. **23f, g**). This gnathostomized lamprey MymK protein, referred to as “gn-lamprey MymK”, differs from human MymK by 32 amino acids but is predicted to adopt a structure that is nearly identical to human MymK (**Fig. 7f**). As expected, gn-lamprey MymK showed fusogenic activity in human *MymK*^−/−^ myoblasts (**Fig. 7g**; Extended Data Fig. **23h**). Together, these results suggest that, although MymK-driven myoblast fusion likely emerged in the last common tunicate-vertebrate ancestor, MymK may have undergone structural refinements (**Fig. 7d**), possibly concomitantly with its interacting partners, after the split between jawed and jawless vertebrates. While tunicate MymK has retained a relatively conservative structure, possibly explaining its modest fusogenic activity in mammalian cells, cyclostome MymK structure has diverged more substantially. We propose that such structural and functional adaptations facilitated the elaboration of multinucleated myofibers in different vertebrate groups (**Fig. 8**).

**Figure 8:**
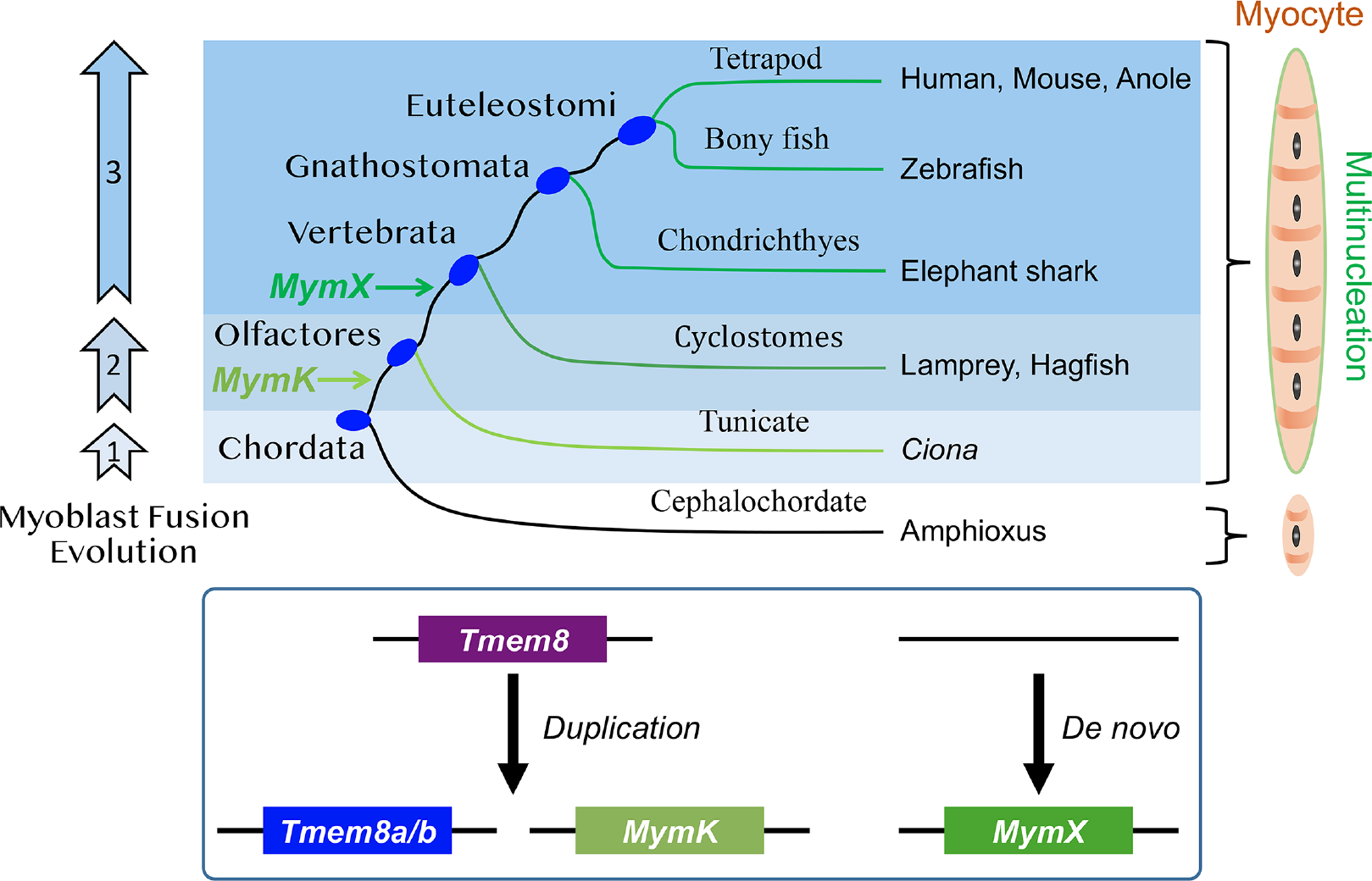
Evolution of chordate-specific control system of myoblast fusion. The emergence of *MymK* gene after duplication of the eukaryotic *Tmem8* gene allowed the multinucleation of muscle cells for common ancestors of tunicates and vertebrates. The emergence of MymX and structural adaptions of MymK proteins drove the extensive and obligatory fusion in vertebrates. Note that Tmem8a/b is called Tmem8-related in tunicates simply due to poor resolution of phylogenetic classification. In vertebrates, this gene became duplicated again to give rise to Tmem8a and Tmem8b.

## Discussion

Myoblast fusion is a prominent feature of vertebrate muscle morphogenesis. Here we have uncovered the evolutionary history of the genetic mechanism of chordate myoblast fusion. We have identified the evolutionary origins of MymK and MymX, and shown that their functions appear to be conserved even between distantly related species. Novel protein-coding genes can arise either through horizontal transfer, duplication and neofunctionalization of existing genes, or *de novo*^42–44^. Whereas the MymK gene was certainly generated through duplication of an ancestral *Tmem8* gene, MymX, as an orphan gene, might have arisen *de novo* (**Fig. 8**). One scenario for the birth of the *MymX* gene could be through a transitory proto-gene that produced a short polypeptide, given the short length (<100 amino acids) of MymX proteins in most vertebrates. After the cyclostome/gnathostome split, MymX may have been secondarily elongated in lampreys (583 amino acids in sea lamprey, 595 amino acids in arctic lamprey). Alternatively, the ancestral MymX protein was closer in size to that of extant lampreys, but was secondarily reduced in length in jawed vertebrates. Of note, possibly attributed to the fact that the hagfish (*Eptatretus burgeri*) genome is not complete^45^, a *MymX* ortholog has yet to be identified in this species.

The emergence of *MymK* and *MymX* genes provides mechanistic explanations for the evolution of myoblast fusion in chordates. First, the lack of *MymK* orthologs outside of Olfactores is consistent with the notion that myoblast fusion evolved independently in chordates and arthropods (e.g. *Drosophila*)^46^. Within the chordates, only tunicates and vertebrates are known to have both *MymK* and multinucleated muscles, while cephalochordates have neither. In theory, multinucleation can be caused by either cell fusion or acytokinetic mitosis, i.e. nuclei divide but cytokinesis is not complete. Development of tunicate multinucleated muscles has been studied by electron microscopy in the thaliacean *Cyclosalpa affinis* and ascidian *Halocynthia roretzi*, which favored myoblast fusion as the cause of tunicate myofiber multinucleation^20, 47^. As in vertebrates, nuclei from developing *Cyclosalpa* and *Halocynthia* myofibers were never seen in mitosis, thus ruling out acytokinetic mitosis^20, 47^. While we found *MymK* in representative genomes from most tunicate groups, including both sessile ascidians and pelagic thaliaceans, we did not find this gene in another group of pelagic tunicates, the appendicularians. Appendicularians are neotenic and retain the larval form throughout their life. As such, they have secondarily lost the multinucleated, post-metamorphic muscles of the siphon or body wall seen in all other tunicates^23^. Like all tunicates, appendicularians have a *Tmem8-related* gene, suggesting they have specifically lost *MymK.* Therefore, throughout the chordates, the presence of the MymK gene correlates with the presence of myoblast fusion (vertebrates, ascidians and thaliaceans), while its absence correlates with lack of myoblast fusion (cephalochordates and appendicularians).

While most tunicates possess MymK, and this MymK promotes mammalian and reptilian myoblast fusion, it is unable to synergize with vertebrate MymX proteins to augment fusion levels. However, we cannot rule out that tunicate MymK can synergize with a distinct endogenous partner(s), which could conceivably play a role similar to MymX in vertebrates yet is so unlike any MymX sequence thus escaping identifications by BLAST. Nonetheless, the absence of *MymX* in tunicates might better explain why their myofibers typically contain a relatively small number of nuclei^17^. By comparison, myoblast fusion in vertebrate species is more pronounced, resulting in the formation of large myofibers that host hundreds of nuclei each. Indeed, fusogenic activity of tunicate MymK in human myoblasts was comparable to that of vertebrate MymK in the absence of MymX. Thus, the appearance of MymX in vertebrates appears to correlate with this increased reliance on myoblast fusion, as evidenced by the synergy of lamprey MymX with mammalian MymK. Considering gene loss is a pervasive source of genetic variation that drives evolution in all life kingdoms^48, 49^, it is also possible that *MymX* (and by extension, MymK–MymX synergy) arose earlier in chordate evolution but was later lost in tunicates, as this group secondarily evolved a biphasic life cycle with a sessile adult phase.

The evolutionary history of myoblast fusion is complicated by the fact that lamprey and hagfish MymK proteins did not elicit fusogenic activity in mammalian or reptilian myoblasts. This was especially curious given the clear fusogenic activity of different tunicate MymK proteins in the same cells. It is possible the action mechanism of MymK diverged specifically in cyclostomes since the last pan-vertebrate ancestor, while both tunicate and gnathostome MymK proteins might be more mechanistically conservative in comparison. Consistent with this possibility, we observed a greater structural divergence of gnathostome MymK proteins from its cyclostome counterparts, relative to tunicate MymK. Indeed, altering the structure of lamprey MymK to more closely resemble that of gnathostomes (gn-lamprey MymK) was sufficient to confer fusogenic activity in mammalian cells. Although we never observed fusogenic activity of any wild-type cyclostome MymK proteins, the ability of lamprey MymX to enhance mammalian MymK function also supports the existence of a conserved MymK–MymX axis for myoblast fusion in cyclostomes. Consistently, the expression of both lamprey MymX and MymK coincides with myoblast fusion during muscle development and is not detected in any adult organs including post-metamorphic muscle tissues. We speculate that the lack of fusogenic ability of cyclostome MymK proteins in gnathostome cells is due to differences of protein structures and how they interact with permissive effector(s)^12^ of downstream processes such as cytoskeleton reorganization, which is an essential step for cell fusion^3^. Therefore, a complete understanding of the evolution and developmental regulation of myoblast fusion will require a thorough characterization of the biochemical mechanism of MymK function in diverse vertebrate and tunicate species. From this regard, the structural models of MymK proteins that we obtained by computational approach in turn affords a solid biochemical basis to identify the unknown permissive factor(s) that may have co-evolved with MymK.

Another untold part of this story is the regulatory evolution of *MymX* and *MymK* genes. The expression of these genes exhibits a high degree of specificity in muscle cells of mice and humans^8, 9, 12^. It appears that the expression of muscle fusogens in both lampreys and tunicates is also restricted to the time-window of myoblast fusion. The evolutionary interpretation of our functional data of diverse MymK and MymX orthologs does not take into account potential differences in the timing, duration and levels of expression in these very different species (e.g. tunicates, lamprey, mammals). Gene regulatory mechanisms might have co-evolved with the differences in protein activity revealed by our heterologous mammalian expression system. For instance, while mammals have four distinct Myogenic Regulatory Factors (MRFs): MyoD, MyoG, Myf5, and Myf6, the lamprey only has two MRF genes and tunicates have a single MRF gene^50, 51^. While the sole tunicate MRF ortholog is expressed in all muscles including mononucleated tail muscles, its partner EBF is required specifically for multinucleated muscle differentiation^27^. Thus, tunicates might selectively activate MymK expression and myoblast fusion through the cooperative activity of MRF and EBF, which are only co-expressed in multinucleated muscles. Given that the duplication and subfunctionalization of MRFs throughout chordate evolution are thought to have allowed for elaboration of different muscle types and greater control over the temporal dynamics of myogenesis^50^, compensatory changes to MymK/MymX function may have evolved to accommodate this changing regulatory landscape.

## Acknowledgements

We thank trainees G. Gopu, A. Baiju, and E. M. Hicks in Bi laboratory and A. L. Womble from Valdosta State University for technical help. We are grateful to E. N. Olson from University of Texas Southwestern Medical Center for critical reading of the manuscript. We thank H. Li from Ocean University of China, C. Cañestro from University of Barcelona, S. Kuraku, R. Kusakabe and S. Kuratani from RIKEN, M. Cui from University of Texas Southwestern Medical Center, S. Du from University of Maryland School of Medicine, J. Ziermann from Howard University, Z. Yang from University of College London, Michael Coates and Tetsuto Miyashita from University of Chicago, and F. Razy-Krajka for advice; A. Bigot and V. Mouly from the Myoline platform of the Myology Institute for myoblast cell lines; X. Li from University of Texas Southwestern Medical Center, N. S. Johnson from United States Geological Survey, M. Brindley from University of Georgia for providing materials and reagents. **Funding**: This work was supported by the starting up fund from the University of Georgia to P.B., NIH R00 award HD084814 and NSF award 1940743 to A.S., an NSF Graduate Research Fellowship to C.J.J., Great Lakes Fishery Commission (540810) to S.D.F. and W.L., and NSF award 1354788 to T.A.U.

## Author contributions

H.Z., R.S., A.S., P.B. designed research; H.Z., R.S., K.K., W.Z., C.J.J., S.L., X.N., L.L., T.A.U., J.Z., L.L., J.P., S.D.F., S.A.G., S.P.S., J.W., J.Z., J.E., D.M., M.E.B., N.V.G., W.L., K.Y., Z.Y., A.S. and P.B. performed research; H.Z., R.S., K.K., W.Z., C.J.J., L.S., X.N., L.L., J.Z., L.L., J.W., W.L., K.Y., Z.Y., A.S. and P.B. analyzed data; A.S. and P.B. wrote the paper.

## Competing interests

The authors declare that they have no competing interests.

## Data and materials availability

All data needed to evaluate the conclusions in the paper are present in the paper and/or the Supplementary Materials. Additional data related to this paper may be requested from the authors.

## Methods

### Human and mouse cell cultures

Human myoblasts (hSkMC-AB1190) were isolated and immortalized as previously published^1, 2^. These cells were cultured in 15% FBS (GemCell, 100-500) and 5% Growth Medium Supplement Mix (PromoCell, C-39365) in Skeletal Muscle Cell Basal Medium (PromoCell, C-23260) with GlutaMAX and 1% Gentamicin Sulfate. Mouse 10T1/2 fibroblasts (ATCC, CCL-226) and C2C12 myoblasts (ATCC, CRL-1772) were maintained in 10% FBS with 1% penicillin/streptomycin (Gibco, 15140122) in DMEM (Dulbecco’s Modified Eagle’s Medium-high glucose, D5796). Myoblast differentiation medium contained 2% horse serum in DMEM with 1% penicillin/streptomycin. Cells have passed mycoplasma test by using the Universal Mycoplasma Detection Kit (American Type Culture Collection, 30-1012K).

### Lizard cell culture and CRISPR experiments in lizard myoblasts

Myogenic single clones (myosin heavy chain+) were isolated from immortalized *Anolis sagrei* embryonic cells ASEC-1 (*Anolis sagrei* embryonic cell line 1; to be described in detail elsewhere). ASEC-1 and clonally derived myoblasts were cultured in DMEM supplemented with glutamine and 10% FBS (with penicillin/streptomycin and amphotericin B) and cultured at 29°C and 5% CO2.

For CRISPR/Cas9-mediated *MymK* knockout experiment, Cas9 and gRNA were transfected to lizard myoblasts using lipofectamine LTX Plus kit (Thermo Fisher Scientific, A12621). pSpCas9(BB) plasmid was a gift from Feng Zhang (Addgene plasmid # 62988)^3^. Puromycin (25 ug/ml) selection was performed for 24 hours starting from 48 hours post transfection. Single clone was isolated and allowed to expand. *MymK* genotypes for each clone were analyzed by PCR followed by Sanger sequencing. Sequences for sgRNAs and genotyping PCR primers are provided in **Supplemental Table 1.**

### Animal husbandry

Standard operating procedures for transporting, maintaining, handling and euthanizing of sea lamprey and hagfish were approved by the Institutional Committee on Animal Use and Care of Michigan State University and California Institute of Technology, Valdosta State University, University of Georgia and in compliance with standards defined by the National Institutes of Health Guide for the Care and Use of Laboratory Animals.

Sea lampreys were trapped in tributaries of Lakes Huron and Michigan by the United States Fish and Wildlife Service and Fisheries and Oceans Canada. Captured lampreys were transported to the United States Geological Survey, Hammond Bay Biological Station (HBBS), Millersburg Michigan and held in 200– 1,000 L tanks that were continually fed with ambient temp, aerated Lake Huron water. To produce sexually mature ovulated females and males for embryo collection, sea lamprey were transferred to the Ocqueoc River, Millersburg Michigan and held in cages (0.5 m^3^) constructed of polyvinyl chloride and polyurethane mesh, allowing natural sexual maturation in a riverine environment. Sea lamprey were checked daily for sexual maturation. Sexually mature males were identified by applying abdominal pressure and checking milt expression^4^. Sexually mature females were identified by applying abdominal pressure and checking for ovulated oocyte expression^5^ along with visual observation of secondary sexual characteristics^6^. Sexually mature males and female lampreys were returned to HBBS where they were held until used for collecting and culturing lamprey embryos as previously outlined^7^. Embryo viability was determined using techniques established for evaluation of the sterile male release program in the Laurentian Great Lakes^8^. Embryos were checked daily for viability and dead embryos were removed from holding containers. Embryos were pooled together for individual samples according to Piavis stages^9^.

Female Atlantic hagfishes (*Myxine glutinosa*, Linnaeus, 1758) were used in this study (Specimen/mass/length; #1/64g/45cm; #2/57g/41cm; #3/55g/43cm). Live specimens were collected at Shoals Marine Lab (Appledore Island, ME) and transported to Valdosta State University (VSU). Specimens were euthanized using 400mg MS222 (Finquel anaesthetic, Argent Chemicals, Redmond WA) and 200mg NaHCO_3_ (pH buffer) mixed in one liter of filtered artificial seawater. An incision was then made along the ventral midline in order to collect tissue specimens for the histological analysis. Preserved amphioxus and shark specimens were obtained from VWR (470001-802, 470001-486). Subsequent paraffin processing, embedding, sectioning and H&E staining were performed by standard procedures^10, 11^.

### Lentivirus preparation and CRISPR experiments in human and mouse myoblasts

sgRNAs that target the coding regions of human and mouse *MymX* and *MymK* genes were individually cloned into the Lenti-CRISPR v2 vector and validated by Sanger sequencing. Lenti-CRISPR v2 vector was a gift from Feng Zhang (Addgene plasmid # 52961)^12^. sgRNA sequences are provided in **Supplemental Table 1.**

For lentivirus production, Lenti-X 293T cells (Clontech, 632180) were cultured in DMEM (containing 1% penicillin/streptomycin, 10% FBS). Transfection was performed using FuGENE6 (Promega, E2692) with psPAX2 and pMD2.G plasmids. At 48 hours post transfection, lentivirus supernatant was collected, filtered and concentrated by Lenti-X Concentrator (Clontech, PT4421-2) following the manufacturer’s protocol. psPAX2 vector was a gift from Didier Trono (Addgene plasmid # 12260). pMD2.G vector was a gift from Didier Trono (Addgene plasmid # 12259). Human and mouse myoblasts were infected by lentivirus in growth medium. Human *MymX*/*MymK* double knockout myoblast line was generated from a *MymX*^KO^ clone^1^ by infecting lenti-CRISPR MymK sgRNAs. Single clone was isolated, expanded and genotyped by PCR and Sanger sequencing. Sequences for genotyping PCR primers are provided in **Supplemental Table 1**. Human *MyoD*^−/−^ myoblasts were generated and authenticated in previous study^1^.

### Retroviral vector preparations and gene expression

Retroviral expression vector pMXs-Puro (Cell Biolabs, RTV-012) was used for cloning and expressing MymX and MymK orthologs. Open reading frame (ORF) inserts were codon optimized and synthesized by IDT. The DNA sequences were verified by Sanger sequencing. For rescue experiments, the sgRNA insensitive DNA cassettes were used. MyoD-pCLBabe was a gift from Stephen Tapscott (Addgene plasmid # 20917)^13^. pLOVE-GFP plasmid was a gift from Miguel Ramalho-Santos (Addgene plasmid # 15949)^14^. pMXs-Cherry plasmid was generated and described previously^1^.

To produce retrovirus, retroviral plasmid was transfected to HEK293 cells using FuGENE 6 (Promega, E2692). Two days after transfection, viral medium was collected, filtered and used to infect cells assisted by polybrene (Sigma-Aldrich, TR-1003-G). One day after viral infection, cells were switched to growth medium. To induce myogenic differentiation, cells were switched to myoblast differentiation medium (2% horse serum in DMEM with 1% penicillin/streptomycin). Human myoblasts can be fully differentiated three days after switching to differentiation medium. Mouse and lizard myoblasts were differentiated by switching to differentiation medium for at least seven and nine days, respectively.

### Differentiation index and fusion index measurements

Differentiation index was measured as the percentage of nuclei in MF20+ cells in relative to total number of nuclei. Fusion index was measured as the percentage of nuclei number in myotubes (≥ 3 nuclei) in relative to total number of muscle nuclei. Differentiation and fusion indexes were calculated basing on the result of manual counting while treatment information was blinded.

### Quantification and statistical analysis

Quantification results for each experiment were based on at least three independent experiments. For image analysis, randomly chosen views were analyzed. Statistical analyses were carried out with GraphPad Prism 8.3.0. Data are presented as mean ± SEM (standard error of mean). For experiments involving multiple groups, one-way ANOVA with Tukey’s multiple comparison test was performed. For experiments involving only two treatment groups, Student’s *t* test with a two-tail distribution was performed. *P* values < 0.05 were considered statistically significant.

### RNA extraction, cDNA synthesis and real-time PCR

Total RNA was extracted from cells, tissues or embryos using Trizol Reagent (Thermo Fisher Scientific, 15-596-018) according to the manufacturer’s instructions. The RNA quality and concentration were assessed by a spectrophotometer (NanoDrop, Thermo Fisher Scientific) for absorbance at 260 nm and 280 nm. cDNA was synthesized from 2 µg total RNA by reverse transcription using random primers with M-MLV reverse transcriptase (Thermo Fisher Scientific, 28025013). Real-time PCR was performed on QuantStudio 3 Real-Time PCR System (Thermo Fisher Scientific) using SYBR Green Master Mix (Roche) and gene-specific primers. The 2^ΔΔCt^ method was used to compare gene expression levels after normalization to 18S rRNA. Primer sequences are listed in **Supplemental Table 1**.

### Membrane fractionation

Membrane fractionations were performed using the Mem-PERTM Plus Membrane Protein Extraction Kit (Thermo Fisher Scientific, 89842). Briefly, human myoblasts were scrapped off the culture dish into ice-cold PBS with a cell scraper. After centrifugation, cell pellets were washed twice in PBS and permeabilized in cytosol fraction buffer with constant mixing for 10 minutes at 4°C. After centrifugation at 16,000 x g for 15 minutes, the cytosol protein fraction was collected as the supernatant. The pellet was resuspended in membrane protein solubilization buffer and incubated at 4°C for 30 minutes with constant mixing. The membrane protein fraction was collected as the supernatant after 16,000 x g centrifugation for 15 minutes at 4°C.

### Western blotting analyses

Cells were lysed in RIPA buffer (Sigma-Aldrich, R0278) supplemented with complete protease inhibitor (Sigma-Aldrich, 04693159001) and incubated on ice for 15 minutes. Lysates were then centrifuged at 16,000 x g for 15 minutes at 4 °C. Protein supernatant was collected and mixed with 4x Laemmli sample buffer (Bio-Rad, 161-0747). Total 20–40 µg protein was loaded and separated by SDS-PAGE gel electrophoresis. The proteins were transferred to a polyvinylidene fluoride (PVDF) membrane (Sigma-Aldrich, ISEQ00010) and blocked in 5% fat-free milk for one hour at room temperature, and then incubated with the following primary antibodies diluted in 5% milk overnight at 4 °C. GAPDH (Santa Cruz Biotechnology, sc-32233); α-Tubulin (Santa Cruz Biotechnology, sc-8035); Biotin Anti-C-tag Conjugate (Thermo Fisher Scientific, 7103252100); insulin receptor β (Cell Signaling Technology, 3020S); Myomixer (Thermo Fisher Scientific, PA5-47639); Myomaker (mouse monoclonal antibody)^1^. After washes in TBST, PVDF membrane was incubated with the following secondary antibody in blocking buffer for one hour at room temperature. HRP Streptavidin (Vector Laboratories, SA-5004); Donkey anti-sheep IgG-HRP Conjugate (Santa Cruz Biotechnology, sc-2473); Goat Anti-Mouse IgG (H+L)-HRP Conjugate (Invitrogen, A28177); Goat Anti-Rabbit IgG (H+L)-HRP Conjugate (Invitrogen, A27036). Immunodetection was performed using Western Blotting Luminol Reagent (Thermo Fisher Scientific, 34075).

### Immunostaining and microscopy of vertebrate cells

Cells were fixed in 4% PFA/PBS for 10 minutes at room temperature, and permeabilized with 0.2% Triton X-100 in PBS and blocked with 3% BSA/PBS for one hour at room temperature. Cells were incubated with the primary antibody overnight at 4 °C, followed by incubation with Alexa Fluor conjugated secondary antibodies. Myosin (Developmental Studies Hybridoma Bank, MF20), MyoD (Santa Cruz Biotechnology, sc-304). Goat anti-Mouse IgG (H+L), Superclonal™ Recombinant Secondary Antibody, Alexa Fluor 555 (Invitrogen, A28180); Goat anti-Mouse IgG (H+L), Superclonal™ Recombinant Secondary Antibody, Alexa Fluor 488 (Invitrogen, A28175); Goat anti-Rabbit IgG (H+L), Superclonal™ Recombinant Secondary Antibody, Alexa Fluor 555 (Invitrogen, A27039); Goat anti-Rabbit IgG (H+L), Superclonal™ Recombinant Secondary Antibody, Alexa Fluor 488 (Invitrogen, A27034). Nucleus was counterstained with Hoechst 33342. Live cell immunostaining of MymK was performed as previously described^15^. Briefly, cells were first washed with PBS and incubated in blocking buffer (3% BSA/PBS) for 15 min. Primary antibody incubation was performed on ice followed by fixation with 4% PFA/PBS and incubation with secondary antibody. The staining was visualized on a BioTek Lionheart FX Automated Microscope. Fluorescence images were collected by camera on the BioTek Microscope System or Olympus FLUOVIEW FV1200 Confocal Laser Scanning Microscope.

### Molecular phylogenetic analysis

Protein sequences were retrieved from the GenBank, Refseq, Ensembl and Aniseed databases or by BLAST search the genome and transcriptome databases. All sequences were provided in **Supplemental File 2**. To construct the phylogenetic tree, protein sequences were first aligned using MUSCLE^16^ with the default setting. Alignment files were provided as **Supplemental File 3**. Maximum number of iterations was set to 8. Neighbor Joining (NJ) trees were reconstructed from the alignments by the software Geneious Prime (https://www.geneious.com/prime/). Maximum likelihood (ML) trees were built from the alignments using RAxML^17^ (version 8.2.11) with either the JTT+GAMMA or LG+GAMMA models. Bootstrap analysis was carried out with 1,000 replicates for both NJ and ML trees. The RAxML command line for the bootstrap analysis is *raxmlHPC -N100 -m PROTGAMMALGF -fa -s tmem8.aln.fasta -n tmem8 -p470940 -x680848*. Bootstrap support values for internal nodes on the ML phylogenetic trees were calculated by a python program sumtrees.py^18^ with the command line *sumtrees.py -f0 -p -t RAxML_bestTree.tre --replace -F newick -o RAxML_bootstrap.con.tre --no-annotations RAxML_bootstrap.tre*.

### Molecular modeling of MymK protein structures

The tertiary structure prediction of the MymK orthologs is based on the D-I-TASSER pipeline, which is an extension of I-TASSER and C-I-TASSER and integrates the deep-learning-based distance and hydrogen-bonding network models with iterative threading assembly simulations^19–23^. The pipeline consists of four consecutive steps: (i) multiple sequence alignment (MSA) generation by DeepMSA^24^, (ii) PDB template detection by LOMETS3^25^ and deep-learning-based residue-residue distance-map/hydrogen-bonding prediction by DeepPotential, (iii) structure conformation (decoy) sampling by Replica Exchange Monte Carlo (REMC) simulation, and (iv) full-length model construction and atomic-level model refinement.

First, starting from the input sequences, DeepMSA is used to create a set of multiple sequence alignments (MSAs) by iteratively searching the query sequence through whole-genome (Uniref90^26^) and metagenome sequence databases (Metaclust^27^, BFD^28^, Mgnify^29^, and IMG/M^30^). The MSA with the highest accumulative probability obtained by the TripletRes-predicted^31^ top 10*L* (*L* is the protein length) contacts is selected. In the second step, the selected MSA is used as the input for template detection by LOMETS3 and distance-map and hydrogen-bonding prediction by DeepPotential. LOMETS3, a newly developed meta-server program combining both profile- and contact-based threading programs, is used to identify structural templates from a non-redundant PDB structural library, while DeepPotential is a newly developed deep residual neural network-based predictor to create multiple spatial restraints, including Cα-Cα and Cβ-Cβ distances and hydrogen-bonding networks. In the third step, the continuous fragments excised from the LOMETS3 templates are used as the initial conformations for full-length structure assembly using replica-exchange Monte Carlo (REMC) simulations under the guidance of a composite force field, including: (i) optimized knowledge-based energy term, (ii) spatial restraints collected from LOMETS3 templates, and (iii) deep-learning distance and hydrogen-bonding restraints obtained from DeepPotential. Finally, at least 10,000 decoys generated by the low-temperature replicas are submitted to SPICKER^32^ for structure clustering and model selection based on the energy and structure similarity. The largest SPICKER cluster is further refined by the atomic-level fragment-guided molecular dynamic (FG-MD^33^) simulations, with the side-chain rotamer structure repacked by FASPR^34^. All MymK structural models are provided in **Supplemental File 4**.

The D-I-TASSER algorithm (named as “Zhang-Server”) has participated in the most recent 14th critical assessment of protein structure prediction experiment (CASP14), which is a blind test to assess the protein folding ability of different participated algorithms, and was ranked as the best automatic protein structure prediction server (https://www.predictioncenter.org/casp14/zscores_final.cgi?gr_type=server_only).

### Estimation of structural model quality and similarity

The global quality of structural model is usually appraised by the TM-score^35^ between model and the experimental determined structure:

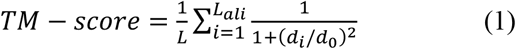

where *L* is the number of residues, *d_i_* is the distance between the *i-*th aligned residue pair between model and experimental structure, and 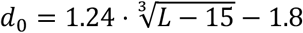 is a scaling factor. TM-score ranges between 0 and 1, and a TM-score greater than 0.5 indicates a structure model of correct global topology^36^.

Because the experimental structure is absent in the present study, instead of actual TM-score, an estimate TM-score (*eTM-score*) was calculated using LOMETS3 threading template quality, contact-map satisfaction rate, and simulation convergence in D-I-TASSER:

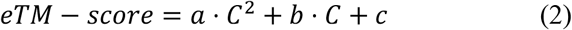

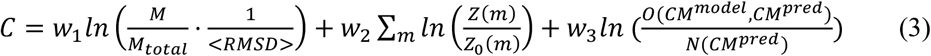

where *a* = 0.00098, *b* = 0.10770, and *c* = 0.79 are fitting parameters retrieved by regression on the large-scale benchmark test data^20–22^, *C* is the overall confidence score of structural assembly, *M_total_* is the total number of decoy conformations submitted to SPICKER clustering, *M* is the number of decoys in the largest cluster, <*RMSD*> is the average RMSD of decoys in the largest cluster, *Z*(*m*) is the significance score of the top template by *m*th threading program, *Z_0_*(*m*) is the cutoff for reliable templates for the *m*th program, *N*(*CM*^*pred*^) is the number of DeepPotential-predicted contacts (predicted distance<8Å) used to guide the REMC simulation, *O*(*CM*^*model*^, *CM*^*pred*^) is the number of overlapping contacts between final models and predicted contacts, and *W*_1_ = 0.77, *W*_2_ = 1.36 and *W*_3_ = 0.67 are free fitting-parameters determined on the large-scale benchmark data. Estimated TM-score (*eTM-score*) highly correlates with the actual TM-score relative to the experimental structures, with a Pearson Correlation Coefficient (PCC) of 0.81 based on a 797 training proteins dataset^20–22^. The similarity among the MymK structural models was calculated as TM-score by superimposing the structural models using TM-align^37^, a sequence-order independent protein structure alignment tool.

### RNA-sequencing data analysis

For bulk RNA-sequencing data, the FASTQ files generated from previous studies^38, 39^ were downloaded from National Center for Biotechnology Information (NCBI) GenBank database with the accession number provided in figure legends. Sequence reads were aligned to genomes by alignment method STAR (Version 2.7.2a) using default setting^40^. Integrative Genomics Viewer (IGV)^41^ was utilized to view the sequencing reads and identify splicing sites and new transcript isoforms.

Single cell RNA-seq data generated from published studies^42, 43^ were reanalyzed for MymK gene expression. Reference genome and annotation files for *C. robusta* were obtained from Aniseed database^44^. MymK locus was manually added to annotation file. Whole larva (18 hpf) single-cell RNA-seq data (GSM3764784, GSM3764785, GSM3764786) were utilized to examine MymK expression in tail muscle cells. Gene-barcode matrices for each sample were generated by 10x Genomics Cell Ranger 3.1.0 using count pipeline under default settings^45^. Downstream analyses were performed by R package Seurat (4.0.)^46^. Cells with fewer than 1,000 expressed genes and genes expressed in fewer than 3 cells were removed. 1,5043 genes across 1,3067 cells were kept in total. Three individual Seurat objects were merged and read counts were normalized and log-transformed for subsequent analysis. Top 1,000 genes with highest standard deviations were selected to exhibit high cell-to-cell variation in the dataset by using FindVariableFeatures function on variance stabilizing transformation method. PCA was performed on the scaled data and statistically significant PCs were determined by heat map pairwise comparison. FindClusters function was used to iteratively group cells by adjusting resolution parameter to 1.4. Expression patterns of genes were visualized by VlnPlot function. Larval tail muscle cell cluster was identified by checking marker genes’ expression.

For re-analysis of the single cell RNA-seq data of the FACS purified cardiopharyngeal-lineage cells (GSE99844), sequence reads of each cell were individually mapped to reference genome using TopHat 2.1.2 with parameter–no-coverage-search^47, 48^. Myomaker FPKM values were calculated by Cufflinks 2.2.1. Clustering results and developmental pseudotime were obtained from the original study^43^. Gene expression patterns were visualized using Seurat R package^46^.

### Software for image and protein sequence analyses

The topology of membrane protein was predicted by TOPCONS^49^. The secondary structure of protein was predicted by PSIPRED^50^. Protein hydrophobicity and similarity were calculated by Expasy^51^. https://web.expasy.org/cgi-bin/sim/sim.pl?prot. Cell and nucleus enumerations for measuring fusion and differentiation indexes were performed using ImageJ (1.52q)^52^.

### *Ciona* embryo handling, electroporation, and immunostaining

Adult *Ciona robusta (intestinalis Type A)* were collected by M-REP (San Diego, USA). Gametes were isolated for *in vitro* fertilization and dechorionation and subsequent electroporation following standard protocols^53, 54^. All plasmid sequences and mixes are described in **Supplemental File 5**. Embryos were raised at 20°C and fixed at the desired stage as calculated by hours post-fertilization. To obtain juveniles, larvae were allowed to metamorphose on (but not attach to) agarose-coated petri dishes in filtered/buffered artificial sea water supplemented with 1X penicillin-streptomycin (Omega Scientific, catalog number PS-20), followed by daily changes of penicillin-streptomycin sea water. For direct visualization of fluorescent proteins, in MEM-FA (3.7% formaldehyde, 0.1M MOPS pH7.4, 0.5M NaCl, 1 mM EGTA, 2 mM MgSO4, 0.05% Triton-X100) for 15 minutes, rinsed in 1X PBS/0.4% Triton-X100/50mM NH4Cl and 1X PBS/0.05% Triton-X100. For immunostaining of CD4::GFP, embryos were fixed and rinsed as above and incubated in mouse anti-GFP (clones 7.1 and 13.1, Roche) at 1:500 dilution for 1 hour, and AlexaFluor 488 goat anti-mouse IgG secondary (ThermoFisher, catalog number A11001) at 1:500 dilution for 1 hour. Both incubations were done in 1X PBS/0.05% Triton-X100/2% Normalized Goat Serum and rinsed in 1X PBS/0.05% Triton-X100. All samples were mounted in 1X PBS/50% Glycerol/2% DABCO.

### In situ hybridization in *Ciona* juveniles

*Ciona* juveniles were raised as described above, fixed in MEM-PFA (4% paraformaldehyde, 0.1M MOPS pH7.4, 0.5M NaCl, 1 mM EGTA, 2mM MgSO4, 0.05% Tween-20) for 2 hours at room temperature or overnight at 4°C, and gradually dehydrated in serial dilutions of ethanol and finally stored in 75% ethanol at –20°C. Whole mount mRNA *in situ* hybridization was carried out as previously described^55, 56^, using the TSA Plus fluorescein detection kit (Akoya Biosciences, catalog number NEL741001KT). Fluorescein-labelled *MymK* riboprobes were prepared by *in vitro* transcription with T7 RNA polymerase from unpurified PCR amplicons of custom-synthesized *MymK* cDNA (Twist Bioscience, see sequence in **Supplemental File 5**).

### Imaging of *Ciona*

Images were acquired on Leica DMi8 and DM IL LED epifluorescence (fig. 2f, g; Extended Data fig. 7c, d; Extended Data **fig. 8b–d**) or Olympus FLUOVIEW FV1200 Confocal Laser Scanning Microscope (fig. 2i; Extended Data fig. 9). The single focal plane images for the representative *Ciona* were used to produce focal plane videos (**Supplemental Videos 1–4**).

### Peakshift assay for sgRNA validation

Embryos were subjected to CRISPR sgRNA validation following the “peakshift” method^57, 58^. Briefly, embryos were electroporated with 25 µg Eef1a>Cas9^59^ and 75 µg of a given U6>sgRNA(F+E)^59^ expression plasmid per 700 µl of electroporation volume. As a negative control for Sanger sequencing chromatogram analysis (see below), U6>Gsx.4(F+E) vector was used instead to drive expression of a sgRNA designed against the unrelated *Gsx* gene instead (see sgRNA sequences below). Embryos were allowed to grow to hatching, then genomic DNA was extracted from each sample of pooled embryos using QIAamp DNA mini kit (Qiagen) following the manufacturers’ recommendations. Purified genomic DNA was then used as template for PCR using Accuprime Pfx (ThermoFisher) following the manufacturer’s recommendations and using a touchdown genomic PCR program as previously described^58^.

Amplicons were PCR-amplified using MymK Peakshift Fwd (CGCGATCACAAATGACGAAAC) and MymK Peakshift Rev (CCCGCAATTACAACATGCTAG) primers. PCR reactions were verified on an agarose gel stained with ethidium bromide, then purified using QIAquick PCR purification kit (Qiagen). Amplicons were sequenced by Sanger sequencing using Exon2seqRev (CCCGCAATTACAACATGCTAG) and Exon4seqFwd (GCATAAGGTGCTGTATGAAACAG) to detect indels in exons 2 and 4, respectively. Sanger sequencing chromatograms were compared between embryos electroporated with MymK sgRNAs and Gsx.4 (negative control) sgRNA using the web application TIDE^60^. Additional sgRNAs targeting exon 3 were tested by sequencing with Exon3seqRev primer (ATTTTGCGTGTCTGAACCTC), but failed to generate any detectable indels.

## Notes

### Competing Interest Statement

The authors have declared no competing interest.

## References

1. Bentzinger, C. F., Wang, Y. X. & Rudnicki, M. A. Building Muscle: Molecular Regulation of Myogenesis. Csh Perspect Biol 4, doi:ARTN a008342 10.1101/cshperspect.a008342 (2012).

2. Buckingham, M. & Rigby, P. W. J. Gene Regulatory Networks and Transcriptional Mechanisms that Control Myogenesis. Dev Cell 28, 225–238, doi:10.1016/j.devcel.2013.12.020 (2014).

3. Rochlin, K., Yu, S., Roy, S. & Baylies, M. K. Myoblast fusion: When it takes more to make one. Developmental Biology 341, 66–83, doi:10.1016/j.ydbio.2009.10.024 (2010).

4. Demonbreun, A. R., Biersmith, B. H. & McNally, E. M. Membrane fusion in muscle development and repair. Semin Cell Dev Biol 45, 48–56, doi:10.1016/j.semcdb.2015.10.026 (2015).

5. Krauss, R. S., Joseph, G. A. & Goel, A. J. Keep Your Friends Close: Cell-Cell Contact and Skeletal Myogenesis. Cold Spring Harb Perspect Biol 9, doi:10.1101/cshperspect.a029298 (2017).

6. Simionescu, A. & Pavlath, G. K. Molecular mechanisms of myoblast fusion across species. Adv Exp Med Biol 713, 113–135, doi:10.1007/978-94-007-0763-4_8 (2011).

7. Abmayr, S. M. & Pavlath, G. K. Myoblast fusion: lessons from flies and mice. Development 139, 641–656, doi:10.1242/dev.068353 (2012).

8. Millay, D. P. et al. Myomaker is a membrane activator of myoblast fusion and muscle formation. Nature 499, 301–305, doi:10.1038/nature12343 (2013).

9. Bi, P. et al. Control of muscle formation by the fusogenic micropeptide myomixer. Science 356, 323–327, doi:10.1126/science.aam9361 (2017).

10. Zhang, Q. et al. The microprotein Minion controls cell fusion and muscle formation. Nat Commun 8, 15664, doi:10.1038/ncomms15664 (2017).

11. Quinn, M. E. et al. Myomerger induces fusion of non-fusogenic cells and is required for skeletal muscle development. Nat Commun 8, 15665, doi:10.1038/ncomms15665 (2017).

12. Zhang, H. et al. Human myotube formation is determined by MyoD–Myomixer/Myomaker axis. Sci Adv In press (2020).

13. Millay, D. P., Sutherland, L. B., Bassel-Duby, R. & Olson, E. N. Myomaker is essential for muscle regeneration. Gene Dev 28, 1641–1646, doi:10.1101/gad.247205.114 (2014).

14. Bi, P. et al. Fusogenic micropeptide Myomixer is essential for satellite cell fusion and muscle regeneration. Proc Natl Acad Sci U S A 115, 3864–3869, doi:10.1073/pnas.1800052115 (2018).

15. Millay, D. P. et al. Structure-function analysis of myomaker domains required for myoblast fusion. Proc Natl Acad Sci U S A 113, 2116–2121, doi:10.1073/pnas.1600101113 (2016).

16. Holland, L. Z. MUSCLE DEVELOPMENT IN AMPHIOXUS: MORPHOLOGY, BIOCHEMISTRY, AND MOLECULAR BIOLOGY. Israel Journal of Zoology 42 (1996).

17. Razy-Krajka, F. & Stolfi, A. Regulation and evolution of muscle development in tunicates. Evodevo 10, 13, doi:10.1186/s13227-019-0125-6 (2019).

18. Kreissl, S., Uber, A. & Harzsch, S. Muscle precursor cells in the developing limbs of two isopods (Crustacea, Peracarida): an immunohistochemical study using a novel monoclonal antibody against myosin heavy chain. Dev Genes Evol 218, 253–265, doi:10.1007/s00427-008-0216-1 (2008).

19. Lee, D. M. & Chen, E. H. Drosophila Myoblast Fusion: Invasion and Resistance for the Ultimate Union. Annu Rev Genet 53, 67–91, doi:10.1146/annurev-genet-120116-024603 (2019).

20. Toselli, P. A. & Harbison, G. R. The fine structure of developing locomotor muscles of the pelagic tunicate, cyclosalpa affinis (Thaliacea: Salpidae). Tissue Cell 9, 137–156, doi:10.1016/0040-8166(77)90055-6 (1977).

21. Sugi, H. & Suzuki, S. Ultrastructural and physiological studies on the longitudinal body wall muscle of Dolabella auricularia. I. Mechanical response and ultrastructure. J Cell Biol 79, 454–466, doi:10.1083/jcb.79.2.454 (1978).

22. K. Terakado, T. O. Structure of multinucleated smooth muscle cells of the ascidian Halocynthia roretzi. Cell and Tissue Research 247, 85–94 (1987).

23. Ferrandez-Roldan, A. et al. Cardiopharyngeal deconstruction and ancestral tunicate sessility. bioRxiv (2021).

24. Satou, Y. et al. Improved genome assembly and evidence-based global gene model set for the chordate Ciona intestinalis: new insight into intron and operon populations. Genome Biology 9, R152, doi:10.1186/gb-2008-9-10-r152 (2008).

25. Wang, W. et al. A single-cell transcriptional roadmap for cardiopharyngeal fate diversification. Nat Cell Biol 21, 674–686, doi:10.1038/s41556-019-0336-z (2019).

26. Cao, C. et al. Comprehensive single-cell transcriptome lineages of a proto-vertebrate. Nature 571, 349–354, doi:10.1038/s41586-019-1385-y (2019).

27. Razy-Krajka, F. et al. Collier/OLF/EBF-dependent transcriptional dynamics control pharyngeal muscle specification from primed cardiopharyngeal progenitors. Dev Cell 29, 263–276, doi:10.1016/j.devcel.2014.04.001 (2014).

28. Stolfi, A. et al. Early chordate origins of the vertebrate second heart field. Science 329, 565–568, doi:10.1126/science.1190181 (2010).

29. Smith, J. J. et al. Sequencing of the sea lamprey (Petromyzon marinus) genome provides insights into vertebrate evolution. Nat Genet 45, 415–421, 421e411-412, doi:10.1038/ng.2568 (2013).

30. Shi, J. et al. Requirement of the fusogenic micropeptide myomixer for muscle formation in zebrafish. Proc Natl Acad Sci U S A 114, 11950–11955, doi:10.1073/pnas.1715229114 (2017).

31. Nakao, T. Electron microscopic studies on the myotomes of larval lamprey, Lampetra japonica. Anat Rec 187, 383–404, doi:10.1002/ar.1091870309 (1977).

32. Fan, C. M., Li, L., Rozo, M. E. & Lepper, C. Making skeletal muscle from progenitor and stem cells: development versus regeneration. Wiley Interdiscip Rev Dev Biol 1, 315–327, doi:10.1002/wdev.30 (2012).

33. Yin, H., Price, F. & Rudnicki, M. A. Satellite Cells and the Muscle Stem Cell Niche. Physiological Reviews 93, 23–67, doi:10.1152/physrev.00043.2011 (2013).

34. Pearce, R. & Zhang, Y. Deep learning techniques have significantly impacted protein structure prediction and protein design. Curr Opin Struct Biol 68, 194–207, doi:10.1016/j.sbi.2021.01.007 (2021).

35. Roy, A., Kucukural, A. & Zhang, Y. I-TASSER: a unified platform for automated protein structure and function prediction. Nat Protoc 5, 725–738, doi:10.1038/nprot.2010.5 (2010).

36. Yang Li, W. Z., Chengxin Zhang, Eric Bell, Xiaoqiang Huang, Robin Pearce, Xiaogen Zhou, Yang Zhang. Protein 3D Structure Prediction by D-I-TASSER in CASP14. CASP14 abstract, 339–341 (2020).

37. Xu, J. & Zhang, Y. How significant is a protein structure similarity with TM-score = 0.5? Bioinformatics 26, 889–895, doi:10.1093/bioinformatics/btq066 (2010).

38. Zhang, Y. & Skolnick, J. Scoring function for automated assessment of protein structure template quality. Proteins 57, 702–710, doi:10.1002/prot.20264 (2004).

39. Jumper, J. et al. Highly accurate protein structure prediction with AlphaFold. Nature, doi:10.1038/s41586-021-03819-2 (2021).

40. Baek, M. et al. Accurate prediction of protein structures and interactions using a three-track neural network. Science, doi:10.1126/science.abj8754 (2021).

41. Tanabe, H. et al. Crystal structures of the human adiponectin receptors. Nature 520, 312–316, doi:10.1038/nature14301 (2015).

42. Carvunis, A. R. et al. Proto-genes and de novo gene birth. Nature 487, 370–374, doi:10.1038/nature11184 (2012).

43. Tautz, D. & Domazet-Loso, T. The evolutionary origin of orphan genes. Nat Rev Genet 12, 692–702, doi:10.1038/nrg3053 (2011).

44. Toll-Riera, M. et al. Origin of primate orphan genes: a comparative genomics approach. Mol Biol Evol 26, 603–612, doi:10.1093/molbev/msn281 (2009).

45. Kazuaki Yamaguchi, Y. H., Kaori Tatsumi, Osamu Nishimura, Jeramiah J. Smith, Mitsutaka Kadota, Shigehiro Kuraku. Inference of a genome-wide protein-coding gene set of the inshore hagfish Eptatretus burgeri. BioRxiv (2020).

46. Paniagua, R., Royuela, M., Garcia-Anchuelo, R. M. & Fraile, B. Ultrastructure of invertebrate muscle cell types. Histol Histopathol 11, 181–201 (1996).

47. Yoshiko Shinohara, K. K. Ultrastructure of the Body-WalI Muscle of the Ascidian Halocynthia roretzi: Smooth Muscle Cell With Multiple Nuclei. The journal of Experimental Zoology 221, 137–142 (1982).

48. Albalat, R. & Canestro, C. Evolution by gene loss. Nat Rev Genet 17, 379–391, doi:10.1038/nrg.2016.39 (2016).

49. Ruddle, F. H., Bentley, K. L., Murtha, M. T. & Risch, N. Gene loss and gain in the evolution of the vertebrates. Dev Suppl, 155–161 (1994).

50. Aase-Remedios, M. E., Coll-Llado, C. & Ferrier, D. E. K. More Than One-to-Four via 2R: Evidence of an Independent Amphioxus Expansion and Two-Gene Ancestral Vertebrate State for MyoD-Related Myogenic Regulatory Factors (MRFs). Mol Biol Evol 37, 2966–2982, doi:10.1093/molbev/msaa147 (2020).

51. Meedel, T. H., Farmer, S. C. & Lee, J. J. The single MyoD family gene of Ciona intestinalis encodes two differentially expressed proteins: implications for the evolution of chordate muscle gene regulation. Development 124, 1711–1721 (1997).

52. Chiba, S. et al. A genomewide survey of developmentally relevant genes in Ciona intestinalis. IX. Genes for muscle structural proteins. Dev Genes Evol 213, 291–302, doi:10.1007/s00427-003-0324-x (2003).

## References

1. Zhang, H. et al. Human myotube formation is determined by MyoD– Myomixer/Myomaker axis. Sci Adv In press (2020).

2. Mamchaoui, K. et al. Immortalized pathological human myoblasts: towards a universal tool for the study of neuromuscular disorders. Skelet Muscle 1, doi:Artn 34 10.1186/2044-5040-1-34 (2011).

3. Ran FA, H. P., Wright J, Agarwala V, Scott DA, Zhang F. Genome engineering using the CRISPR-Cas9 system. Nat Protoc (2013).

4. Siefkes, M. J., Scott, A. P., Zielinski, B., Yun, S. S. & Li, W. Male sea lampreys, Petromyzon marinus L., excrete a sex pheromone from gill epithelia. Biol Reprod 69, 125–132, doi:10.1095/biolreprod.102.014472 (2003).

5. Brant, C. O., Huertas, M., Li, K. & Li, W. Mixtures of Two Bile Alcohol Sulfates Function as a Proximity Pheromone in Sea Lamprey. PLoS One 11, e0149508, doi:10.1371/journal.pone.0149508 (2016).

6. Applegate, V. C. Natural history of the sea lamprey, Petromyzon marinus, in Michigan Doctoral dissertation, University of Michigan (1950).

7. Nikitina, N., Bronner-Fraser, M. & Sauka-Spengler, T. Culturing lamprey embryos. Cold Spring Harb Protoc 2009, pdb prot5122, doi:10.1101/pdb.prot5122 (2009).

8. Campbell, B. Nest Survey Procedures and Examples for the Sterile Male Release Program. (2003).

9. Piavis, G. W. mbryological stages in the sea lamprey and effects of temperature on development *US Government Printing Office*, 111–143 (1961).

10. Bi, P. et al. Control of muscle formation by the fusogenic micropeptide myomixer. Science 356, 323–327, doi:10.1126/science.aam9361 (2017).

11. D. C. Shehan, B. B. H. Theory and Practice of Histotechnology. Battelle Press (1980).

12. Sanjana, N. E., Shalem, O. & Zhang, F. Improved vectors and genome-wide libraries for CRISPR screening. Nat Methods 11, 783–784, doi:10.1038/nmeth.3047 (2014).

13. Yang, Z. et al. MyoD and E-protein heterodimers switch rhabdomyosarcoma cells from an arrested myoblast phase to a differentiated state. Genes Dev 23, 694–707, doi:10.1101/gad.1765109 (2009).

14. Blelloch, R., Venere, M., Yen, J. & Ramalho-Santos, M. Generation of induced pluripotent stem cells in the absence of drug selection. Cell Stem Cell 1, 245–247, doi:10.1016/j.stem.2007.08.008 (2007).

16. Edgar, R. C. MUSCLE: multiple sequence alignment with high accuracy and high throughput. Nucleic Acids Res 32, 1792–1797, doi:10.1093/nar/gkh340 (2004).

17. Stamatakis, A. RAxML version 8: a tool for phylogenetic analysis and post-analysis of large phylogenies. Bioinformatics 30, 1312–1313, doi:10.1093/bioinformatics/btu033 (2014).

18. Sukumaran, J. & Holder, M. T. DendroPy: a Python library for phylogenetic computing. Bioinformatics 26, 1569–1571, doi:10.1093/bioinformatics/btq228 (2010).

19. Yang Li, W. Z., Chengxin Zhang, Eric Bell, Xiaoqiang Huang, Robin Pearce, Xiaogen Zhou, Yang Zhang. Protein 3D Structure Prediction by D-I-TASSER in CASP14. CASP14 abstract, 339–341 (2020).

20. Roy, A., Kucukural, A. & Zhang, Y. I-TASSER: a unified platform for automated protein structure and function prediction. Nature Protocols 5, 725–738, doi:10.1038/nprot.2010.5 (2010).

21. Yang, J. et al. The I-TASSER Suite: protein structure and function prediction. Nature Methods 12, 7–8, doi:10.1038/nmeth.3213 (2015).

22. Zheng, W. et al. Deep-learning contact-map guided protein structure prediction in CASP13. *Proteins: Structure*, Function, and Bioinformatics 87, 1149–1164, doi:10.1002/prot.25792 (2019).

23. Yang Li, C. Z., Wei Zheng, Xiaogen Zhou, Eric W. Bell, Dong-Jun Yu, Yang Zhang. Learning deep statistical potentials for protein folding. CASP14 abstract, 72–73 (2020).

24. Zhang, C., Zheng, W., Mortuza, S. M., Li, Y. & Zhang, Y. DeepMSA: constructing deep multiple sequence alignment to improve contact prediction and fold-recognition for distant-homology proteins. Bioinformatics 36, 2105–2112, doi:10.1093/bioinformatics/btz863 (2020).

25. Wei Zheng, Y. L., Xiaogen Zhou, Chengxin Zhang, Robin Pearce, Yang Zhang. Template-based protein folding guided by residue-residue distance and hydrogen-bond network prediction from deep-learning. CAS*P*14 *abstract*, 342-344 (2020).

26. Suzek, B. E. et al. UniRef clusters: a comprehensive and scalable alternative for improving sequence similarity searches. Bioinformatics 31, 926–932, doi:10.1093/bioinformatics/btu739 (2015).

27. Steinegger, M. & Söding, J. Clustering huge protein sequence sets in linear time. Nature Communications 9, 2542, doi:10.1038/s41467-018-04964-5 (2018).

28. Steinegger, M., Mirdita, M. & Söding, J. Protein-level assembly increases protein sequence recovery from metagenomic samples manyfold. Nature Methods 16, 603–606, doi:10.1038/s41592-019-0437-4 (2019).

29. Mitchell, A. L. et al. MGnify: the microbiome analysis resource in 2020. Nucleic Acids Research 48, D570–D578, doi:10.1093/nar/gkz1035 (2020).

30. Chen, I. M. A. et al. IMG/M v.5.0: an integrated data management and comparative analysis system for microbial genomes and microbiomes. Nucleic Acids Research 47, D666–D677, doi:10.1093/nar/gky901 (2019).

31. Li, Y., Zhang, C., Bell, E. W., Yu, D.-J. & Zhang, Y. Ensembling multiple raw coevolutionary features with deep residual neural networks for contact-map prediction in CASP13. *Proteins: Structure*, Function, and Bioinformatics 87, 1082–1091, doi:10.1002/prot.25798 (2019).

32. Zhang, Y. & Skolnick, J. SPICKER: A clustering approach to identify near-native protein folds. Journal of Computational Chemistry 25, 865–871, doi:https://doi.org/10.1002/jcc.20011 (2004).

33. Zhang, J., Liang, Y. & Zhang, Y. Atomic-Level Protein Structure Refinement Using Fragment-Guided Molecular Dynamics Conformation Sampling. Structure 19, 1784–1795, doi:https://doi.org/10.1016/j.str.2011.09.022 (2011).

34. Huang, X., Pearce, R. & Zhang, Y. FASPR: an open-source tool for fast and accurate protein side-chain packing. Bioinformatics 36, 3758–3765, doi:10.1093/bioinformatics/btaa234 (2020).

35. Zhang, Y. & Skolnick, J. Scoring function for automated assessment of protein structure template quality. *Proteins: Structure*, Function, and Bioinformatics 57, 702–710, doi:10.1002/prot.20264 (2004).

36. Xu, J. & Zhang, Y. How significant is a protein structure similarity with TM-score = 0.5? Bioinformatics 26, 889–895, doi:10.1093/bioinformatics/btq066 (2010).

37. Zhang, Y. & Skolnick, J. TM-align: a protein structure alignment algorithm based on the TM-score. Nucleic Acids Research 33, 2302–2309, doi:10.1093/nar/gki524 (2005).

38. Martik, M. L. et al. Evolution of the new head by gradual acquisition of neural crest regulatory circuits. Nature 574, 675–678, doi:10.1038/s41586-019-1691-4 (2019).

39. Pascual-Anaya, J. et al. Hagfish and lamprey Hox genes reveal conservation of temporal colinearity in vertebrates. Nat Ecol Evol 2, 859–866, doi:10.1038/s41559-018-0526-2 (2018).

40. Dobin, A. et al. STAR: ultrafast universal RNA-seq aligner. Bioinformatics 29, 15–21, doi:10.1093/bioinformatics/bts635 (2013).

41. Robinson, J. T. et al. Integrative genomics viewer. Nat Biotechnol 29, 24–26, doi:10.1038/nbt.1754 (2011).

42. Cao, C. et al. Comprehensive single-cell transcriptome lineages of a proto-vertebrate. Nature 571, 349–354, doi:10.1038/s41586-019-1385-y (2019).

43. Wang, W. et al. A single-cell transcriptional roadmap for cardiopharyngeal fate diversification. Nat Cell Biol 21, 674–686, doi:10.1038/s41556-019-0336-z (2019).

44. Tassy, O. et al. The ANISEED database: digital representation, formalization, and elucidation of a chordate developmental program. Genome Res 20, 1459–1468, doi:10.1101/gr.108175.110 (2010).

45. Zheng, G. X. et al. Massively parallel digital transcriptional profiling of single cells. Nat Commun 8, 14049, doi:10.1038/ncomms14049 (2017).

46. al., Y. H. e. Integrated analysis of multimodal single-cell data . bioRxiv DOI: 10.1101/2020.10.12.335331 (2020).

47. Kim, D. et al. TopHat2: accurate alignment of transcriptomes in the presence of insertions, deletions and gene fusions. Genome Biol 14, R36, doi:10.1186/gb-2013-14-4-r36 (2013).

48. Trapnell, C. et al. Differential gene and transcript expression analysis of RNA-seq experiments with TopHat and Cufflinks. Nat Protoc 7, 562–578, doi:10.1038/nprot.2012.016 (2012).

49. Tsirigos, K. D., Peters, C., Shu, N., Kall, L. & Elofsson, A. The TOPCONS web server for consensus prediction of membrane protein topology and signal peptides. Nucleic Acids Res 43, W401–407, doi:10.1093/nar/gkv485 (2015).

50. Buchan, D. W. A. & Jones, D. T. The PSIPRED Protein Analysis Workbench: 20 years on. Nucleic Acids Res 47, W402–W407, doi:10.1093/nar/gkz297 (2019).

51. Gasteiger E., H. C., Gattiker A., Duvaud S., Wilkins M.R., Appel R.D., Bairoch A. The Proteomics Protocols Handbook. 571–607 (2005).

52. Schneider, C. A., Rasband, W. S. & Eliceiri, K. W. NIH Image to ImageJ: 25 years of image analysis. Nat Methods 9, 671–675, doi:10.1038/nmeth.2089 (2012).

53. Christiaen, L., Wagner, E., Shi, W. & Levine, M. Isolation of sea squirt (Ciona) gametes, fertilization, dechorionation, and development. Cold Spring Harbor Protocols 2009, pdb. prot5344 (2009).

54. Christiaen, L., Wagner, E., Shi, W. & Levine, M. Electroporation of transgenic DNAs in the sea squirt Ciona. Cold Spring Harbor Protocols 2009, pdb. prot5345 (2009).

55. Christiaen, L., Wagner, E., Shi, W. & Levine, M. Whole-mount in situ hybridization on sea squirt (Ciona intestinalis) embryos. Cold Spring Harbor Protocols 2009, pdb. prot5348 (2009).

56. Racioppi, C. et al. Fibroblast growth factor signalling controls nervous system patterning and pigment cell formation in Ciona intestinalis. Nature communications 5, 1–17 (2014).

57. Gandhi, S., Haeussler, M., Razy-Krajka, F., Christiaen, L. & Stolfi, A. Evaluation and rational design of guide RNAs for efficient CRISPR/Cas9-mediated mutagenesis in Ciona. Developmental biology 425, 8–20 (2017).

58. Gandhi, S., Razy-Krajka, F., Christiaen, L. & Stolfi, A. in Transgenic Ascidians 141–152 (Springer, 2018).

59. Stolfi, A., Gandhi, S., Salek, F. & Christiaen, L. Tissue-specific genome editing in Ciona embryos by CRISPR/Cas9. Development 141, 4115–4120 (2014).

60. Brinkman, E. K., Chen, T., Amendola, M. & van Steensel, B. Easy quantitative assessment of genome editing by sequence trace decomposition. Nucleic Acids Research 42, e168–e168, doi:10.1093/nar/gku936 (2014).

