## Extended Data Fig. 1-23 for "Evolution of a chordate-specific mechanism for myoblast fusion"

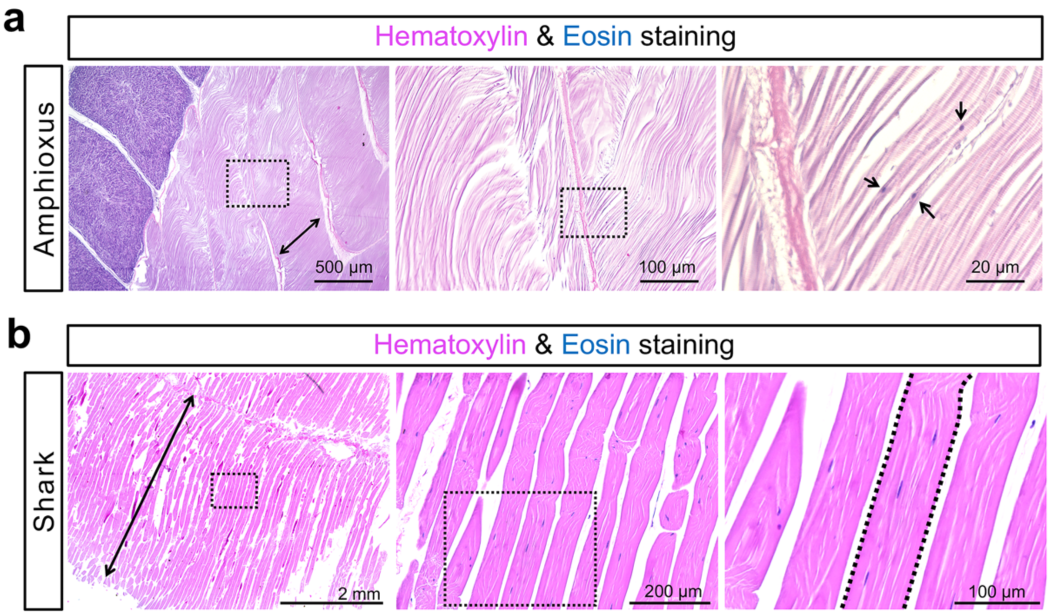


**Extended Data Fig. 1: Muscle histological analyses of adult amphioxus and shark.**

Histology of longitudinal sections of muscle tissues dissected from adult amphioxus (**a**) and shark (*Squalus acanthias*) (**b**). Arrows point to mononucleated myocytes in **a**. A region of multinucleated myofibers is outlined in **b**.

Extended Data Fig. 2

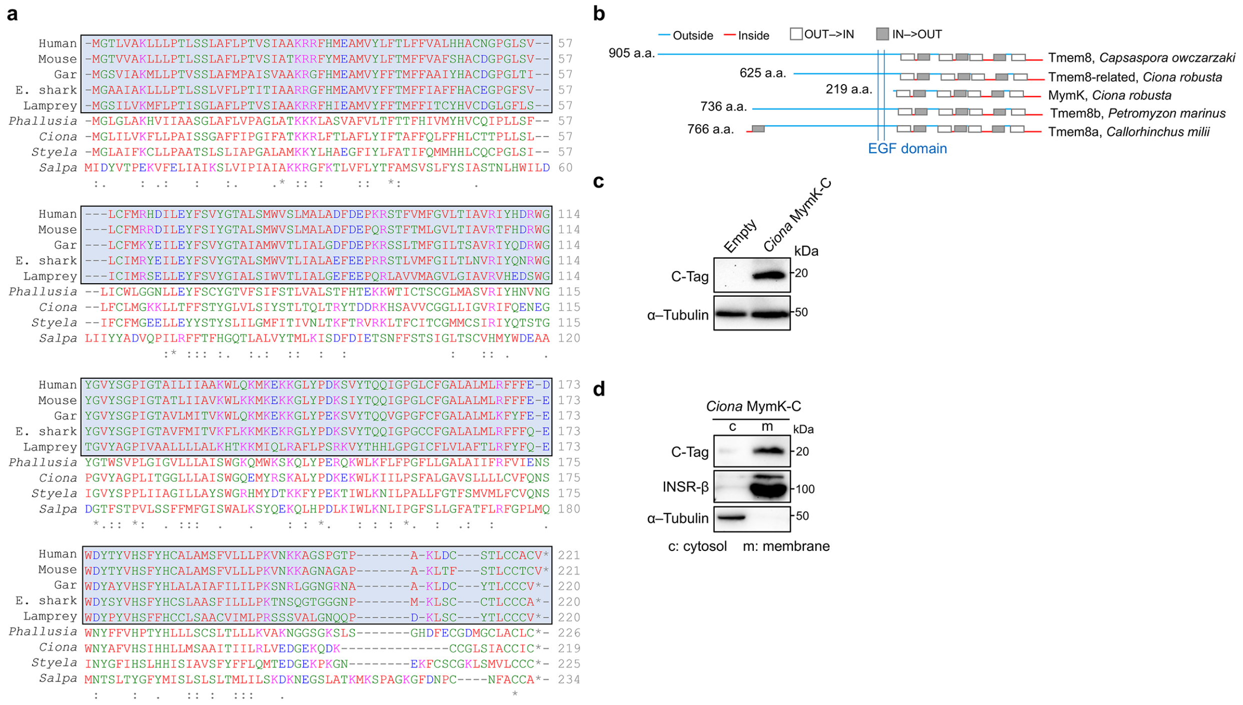


**Extended Data Fig. 2: Sequence alignments and expression of MymK orthologs.**

**a**, Alignment of MymK proteins from eight species: Human (*Homo sapiens*), Mouse (*Mus musculus*), Gar (*Lepisosteus oculatus*), Elephant shark (*Callorhinchus milii*), Lamprey (*Petromyzon marinus*), *Phallusia* (*Phallusia mammillata*), *Ciona* (*Ciona robusta*), *Styela* (*Styela clava*) and *Salpa* (*Salpa thompsoni*). Substantial degree of sequence identity is observed for vertebrate MymK sequences (boxed) but not MymK from tunicates (*Phallusia*, *Ciona*, *Styela* and *Salpa*). **b**, Topology and domain predictions (SCAMPI model) for Tmem8 family transmembrane proteins. EGF domain: epidermal growth factor like domain. **c**, **d**, Western blots confirming the expression of C-tagged *Ciona* MymK protein in human myoblasts (**c**, total protein; **d**, after membrane fractionation). α-Tubulin blot was used as a positive control of cytosolic proteins. Insulin receptor β (INSR-β) blot was used as a positive control of membrane proteins.

Extended Data Fig. 3

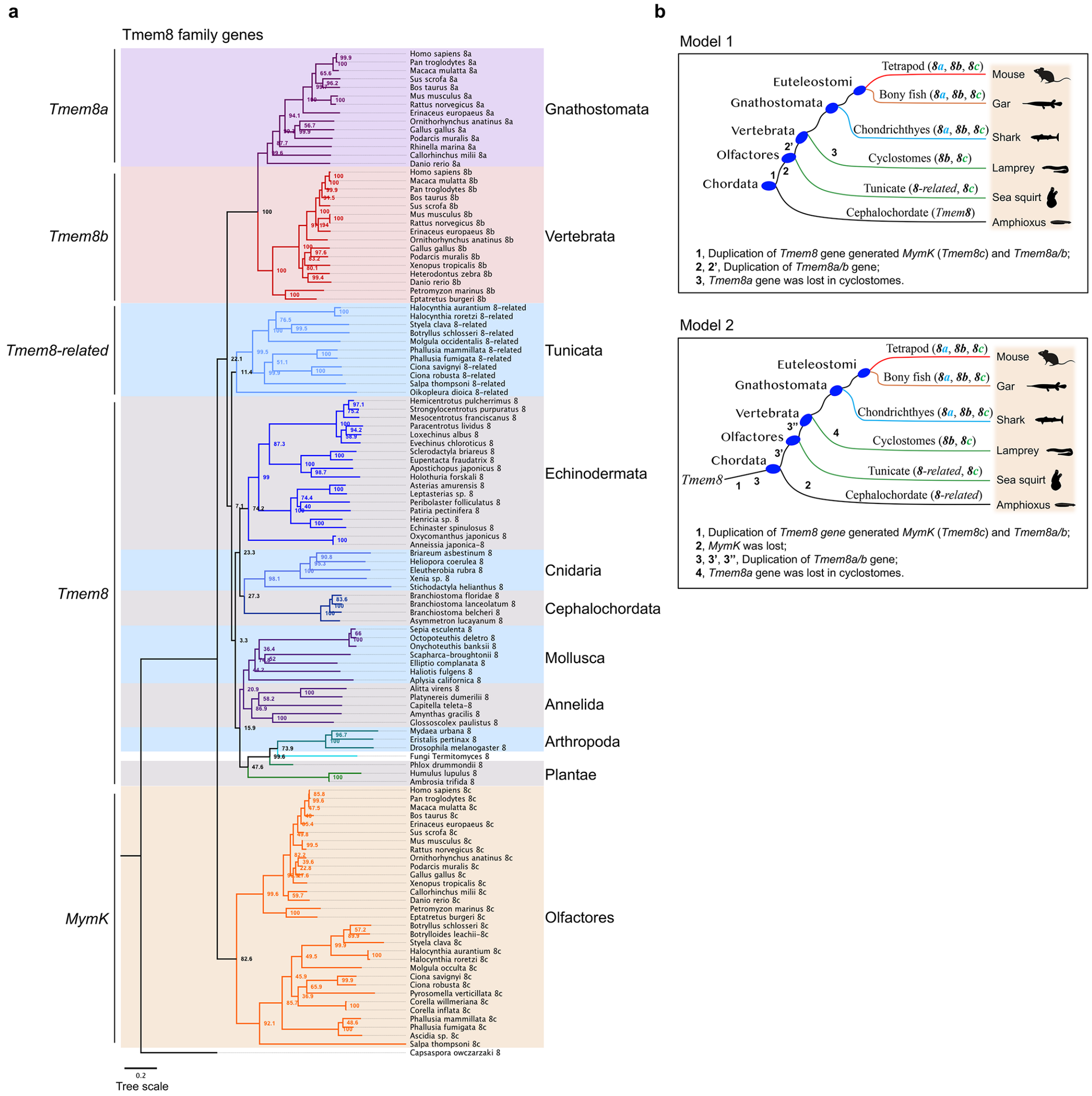


**Extended Data Fig. 3: Phylogenetic analysis of the Tmem8 gene family.**

**a**, Phylogenetic tree of *Tmem8* family genes inferred by a distance-based method (neighbour joining). The bootstrap percentages obtained from 1,000 replicates were shown in the cladogram. **b**, Scenarios of two-round gene duplications that gave rise to various Tmem8 family gene members in Chordata. Model 1 represents a more parsimonious scenario of evolution.

Extended Data Fig. 4

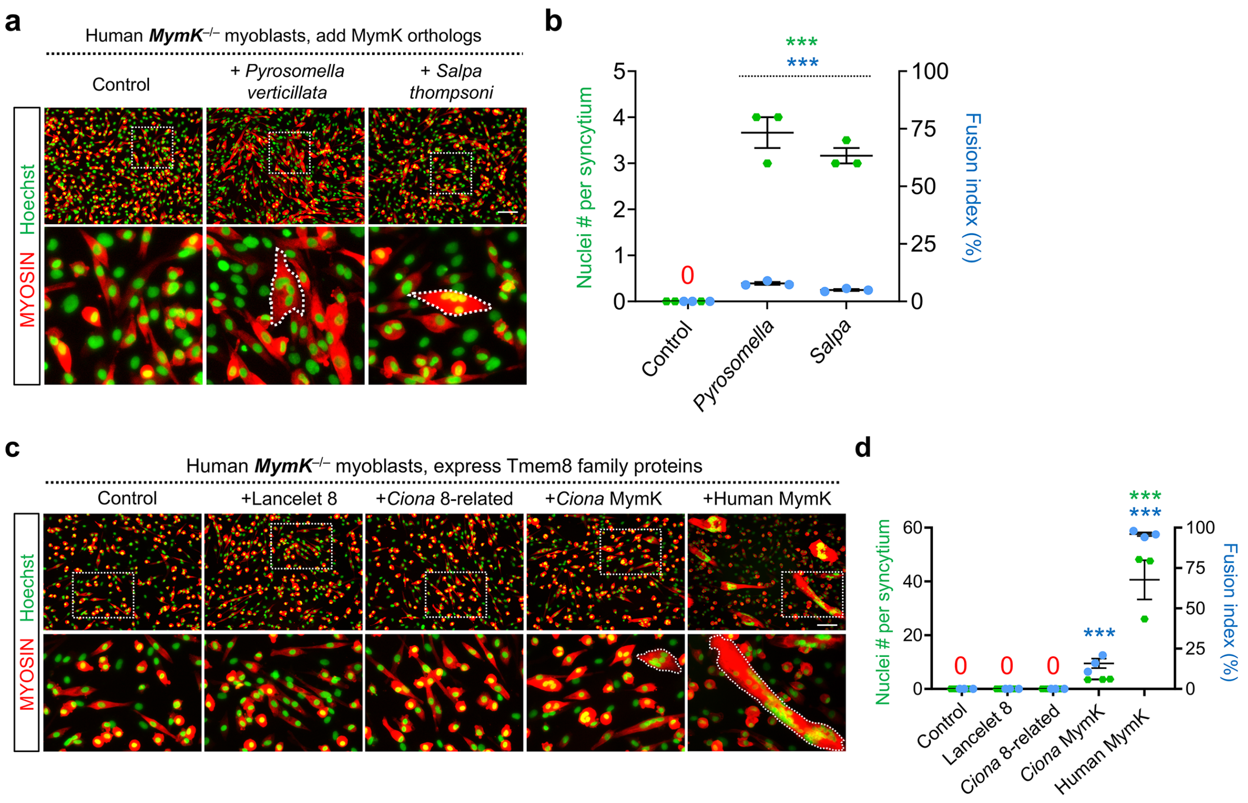


**Extended Data Fig. 4: Expression of *MymK* but not *Tmem8*-related genes induces fusion of human *MymK*$^{-/-}$ myoblasts.**

**a**, Myosin immunostaining of human *MymK*$^{-/-}$ myoblasts transfected with control (empty vector) or expression vectors of MymK orthologs from thaliaceans. **b**, Measurements of myoblast fusion for groups in **a** after 3 days of differentiation. **c**, Myosin immunostaining of human *MymK*$^{-/-}$ myoblasts transfected with control (empty vector) or expression vectors of Tmem8 family genes. Only *MymK* genes induce cell multinucleations (outlined). Lancelet 8: Tmem8 gene from *Branchiostoma floridae*; *Ciona* 8-related: Tmem8-related gene from *Ciona robusta*; *Ciona* MymK: Tmem8c gene from *Ciona robusta*. **d**, Measurements of myoblast fusion in **c** after 5 days of differentiation. Scale bars, 100 µm. Data are means ± SEM. *** $P < 0.001$, compared to control group, one-way ANOVA.

Extended Data Fig. 5

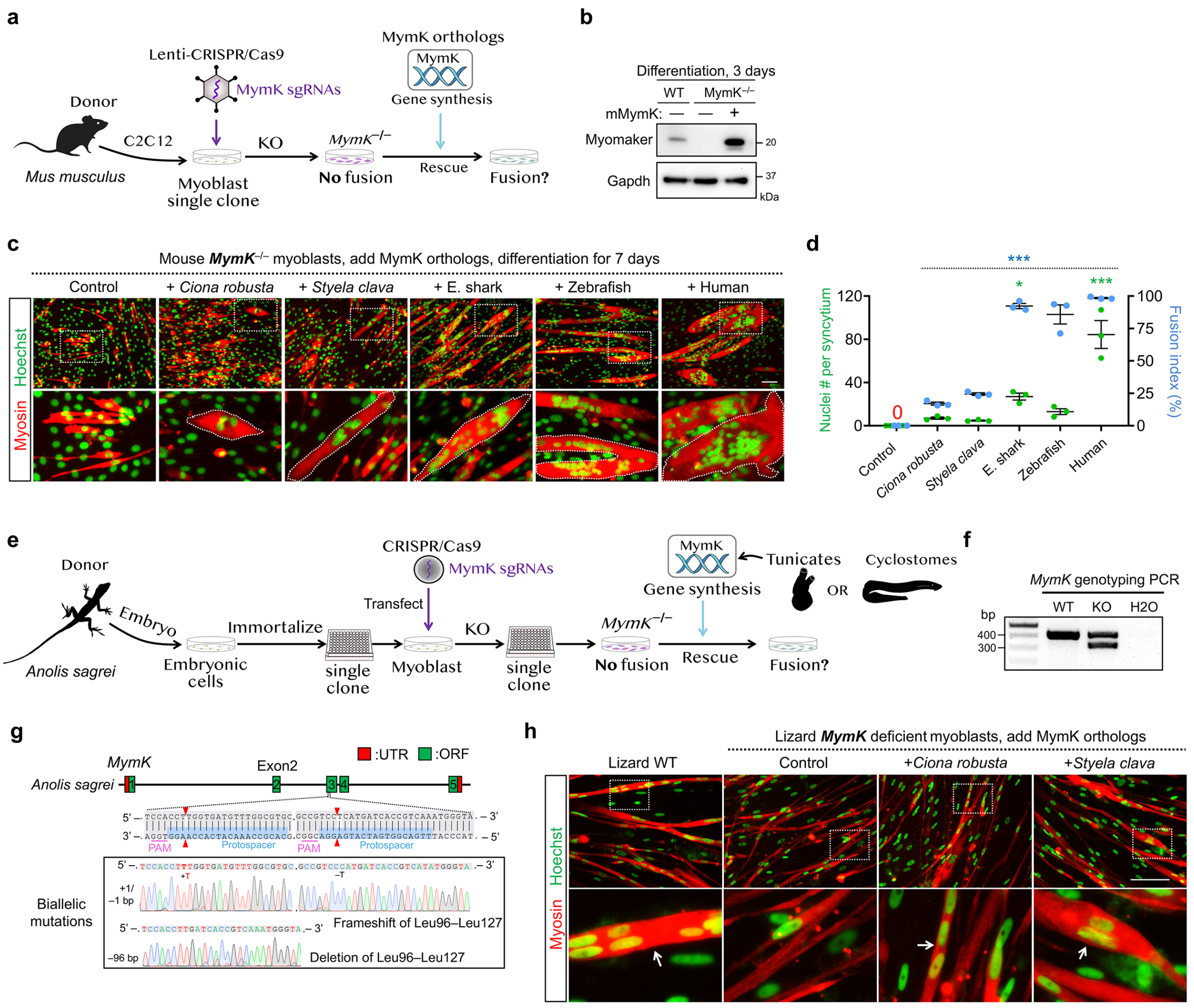


**Extended Data Fig. 5: Tunicate MymK proteins induce fusion of mouse and lizard *MymK*$^{-/-}$ myoblasts.**

**a**, Schematic of experimental design to generate mouse *MymK*-deficient myoblasts and test the fusogenic activity of *MymK* orthologs. Expression of Cas9 and 3 single guide RNA (sgRNAs) that targets exons 1, 2, 3 are delivered by lentiviral infection. **b**, Western blots validated the successful depletions of MymK protein in an isolated single clone of CRISPR treated mouse myoblasts. **c**, Myosin immunostaining of mouse *MymK*$^{-/-}$ myoblasts transfected with MymK orthologs. Tunicate (*Styela* and *Ciona*) MymK genes induced formations of muscle syncytia (outlined), which are smaller compared to syncytia induced by MymK proteins from jawed vertebrates; E. shark: elephant shark. **d**, Measurements of myoblast fusion in **c** after 7 days of differentiation. **e**, Schematic of experimental design to isolate myogenic clone from lizard (*Anolis sagrei*) embryonic cells, inactivate *MymK* gene and finally test the fusogenic activity of tunicate MymK proteins. CRISPR/Cas9-mediated mutagenesis of lizard *MymK* gene was performed using a pair of sgRNAs targeting exon 3. **f**, **g**, Genotyping PCR (**f**) and sanger sequencing (**g**) analyses of lizard *MymK* locus revealed a mutant myoblast clone where both alleles of *MymK* gene are disrupted. The predicted editing sites of sgRNAs are near the codons of Leu96 and Leu127 residues. KO: knockout. **h**, Myosin immunostaining of lizard *MymK*-deficient myoblasts transfected with various *MymK* orthologs. Consistent with the results in human and mouse myoblasts, expression of tunicate MymK proteins induce fusion of lizard myoblasts. Cells are differentiated for 9 days before analysis. Scale bars, 100 μm. Data are means ± SEM. * $P < 0.05$; *** $P < 0.001$, compared to control group, one-way ANOVA.

Extended Data Fig. 6

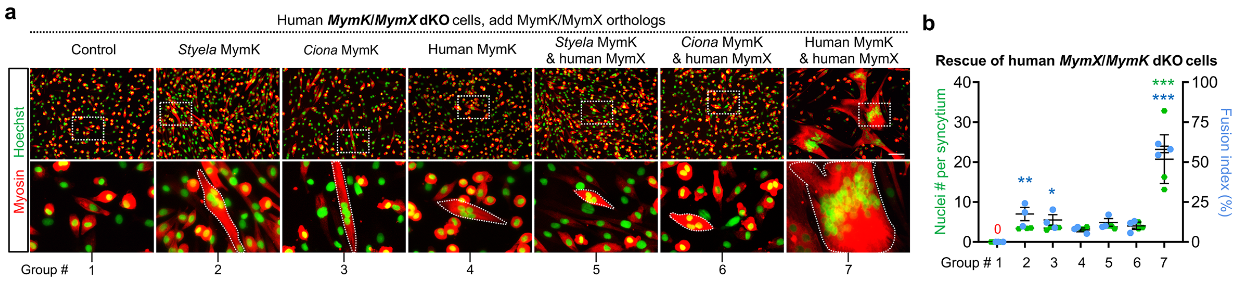


**Extended Data Fig. 6: Fusogenic activity of tunicate MymK proteins is independent of MymX.**

**a**, Myosin immunostaining of human *MymK*/*MymX* dKO myoblasts transfected with MymK orthologs in presence or absence of human MymX re-expression. Muscle syncytia are outlined. Scale bar, 100 µm. **b**, Measurements of myoblast fusion for the expression groups in **a** after 5 days of differentiation. Data are means ± SEM. * $P < 0.05$; ** $P < 0.01$; *** $P < 0.001$, compared to control group, one-way ANOVA.

Extended Data Fig. 7

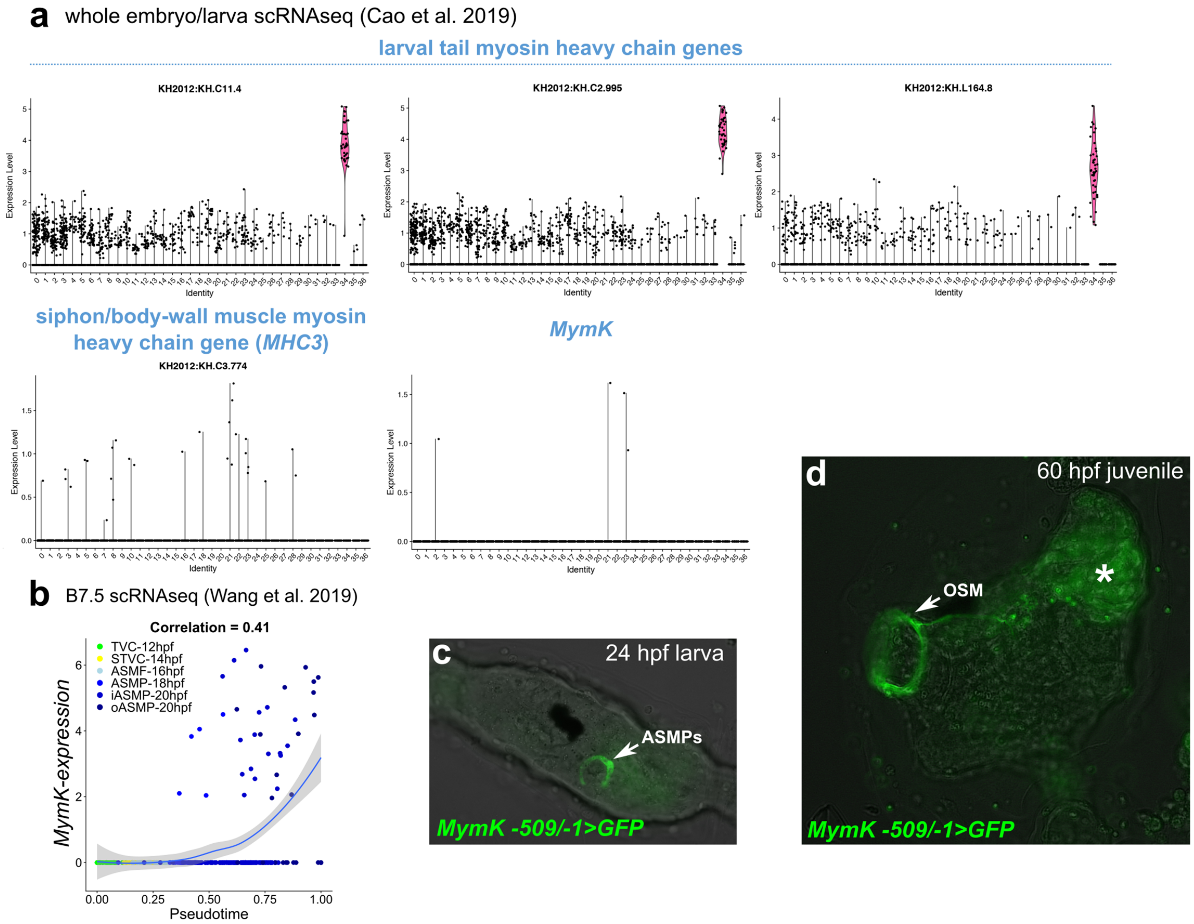


**Extended Data Fig. 7: Mononucleated tail muscle cells from *Ciona robusta* larvae do not express *MymK*.**

**a**, Single-cell RNA sequencing results of *Ciona* larval samples (18 hpf, SRA accession: PRJNA542748) Top row: mononucleated tail muscle cells (cluster 34) identified by the abundant expression of larval tail muscle-specific *Myosin heavy chain* paralogs (*MHC4–6*)[49]. Middle row left: Expression level of post-metamorphic siphon/body-wall muscle marker *MHC3*. Middle row center: *MymK* expression is not detected in the mononucleated tail muscle cells (cluster 34). **b**, Pseudotemporal expression profile of *MymK*, showing upregulation specifically in fate-restricted ASMPs in late larval development. **c**, Larva transfected with *MymK -509/-1>GFP* revealing ASMPs right before settlement and metamorphosis. **d**, Post-metamorphic juvenile transfected with *MymK -509/-1>GFP* revealing expression by oral siphon muscles (OSM) in addition to ASMs (see Fig. 2g). Asterisk indicates autofluorescent reabsorbed larval tail.

Extended Data Fig. 8

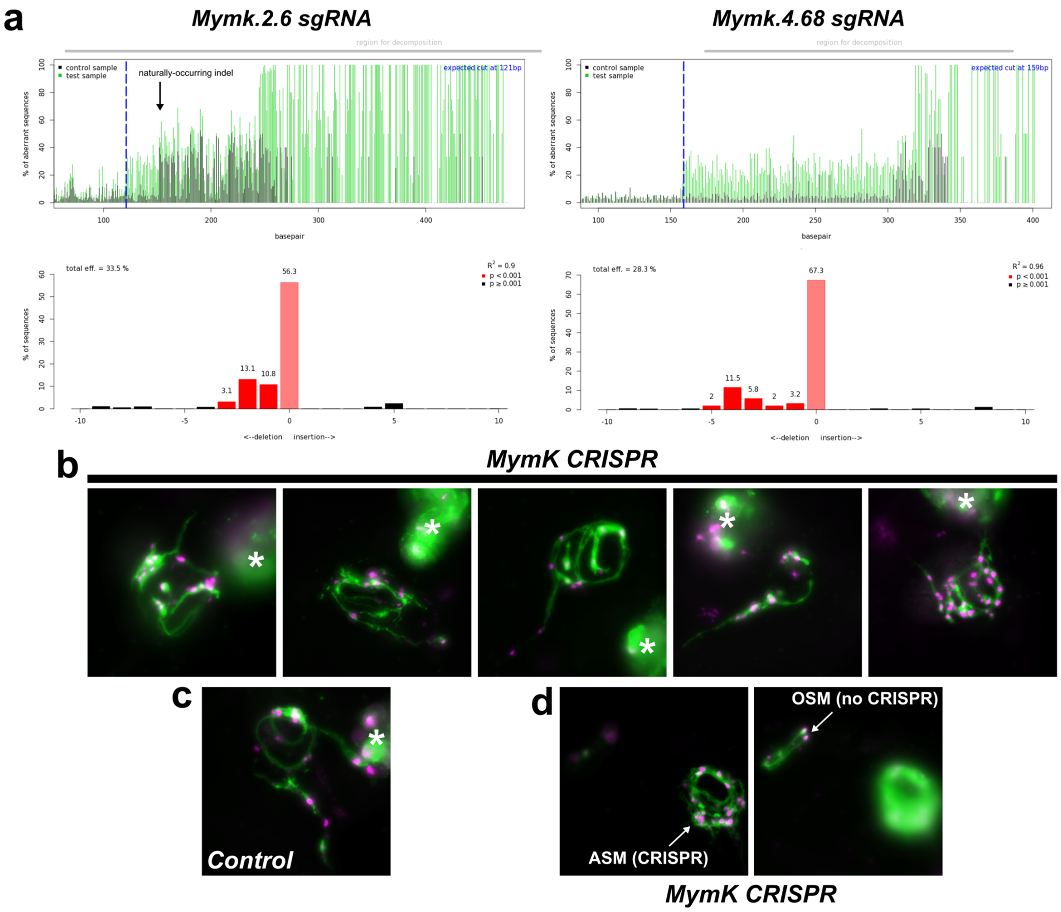


**Extended Data Fig. 8: Knocking out *MymK* from *Ciona robusta* by CRISPR/Cas9.**

**a**, "Peakshift" assay for single guide RNA (sgRNA) validation, performed by Sanger sequencing amplicons from larvae subjected to CRISPR/Cas9-mediated mutagenesis using individual sgRNA designed to target *MymK* in *Ciona robusta.* Two sgRNAs, *MymK.2.6* and *MymK.4.68* were found to generate short indels in the second and fourth exons, respectively. Images generated by TIDE (https://tide.nki.nl/). See Methods for details. **b**, Representative images of developing ASMs in juveniles (60 hpf) subjected to B7.5-specific mutagenesis of *MymK* by CRISPR/Cas9. In 10 of 19 *MymK* CRISPR juveniles, ASMs were disorganized and did not form enclosed rings of circular myofibers around the atrial siphon primordia. Asterisks indicate reabsorbed larval tail muscle cells. **c**, Negative control juvenile (60 hpf, transfected with *Mesp>Cas9* alone) showing clear formation of circular myofibers. **d**, Different focal plane views of a single *MymK* CRISPR juvenile (60 hpf) with both OSMs and ASMs labeled. Defects are only seen in the ASMs, which was expected due to lack of expression of Cas9 in the OSM lineage (*Mesp*–), which in turn serves as an internal negative control. Note that the oral siphon opening is perpendicular to the focal plane (not parallel to the focal plane, like the atrial siphon openings), which means the OSMs are imaged in profile. Muscle plasma membranes and nuclei in panels **b–d** are labeled by *MRF>CD4::GFP* and *MRF>H2B::mCherry,* respectively. All microscopy images were captured on a compound epifluorescence microscope. Images in **b**, **c** are projections of Z stacks, while images in **d** are single focal plane images.

Extended Data Fig. 9

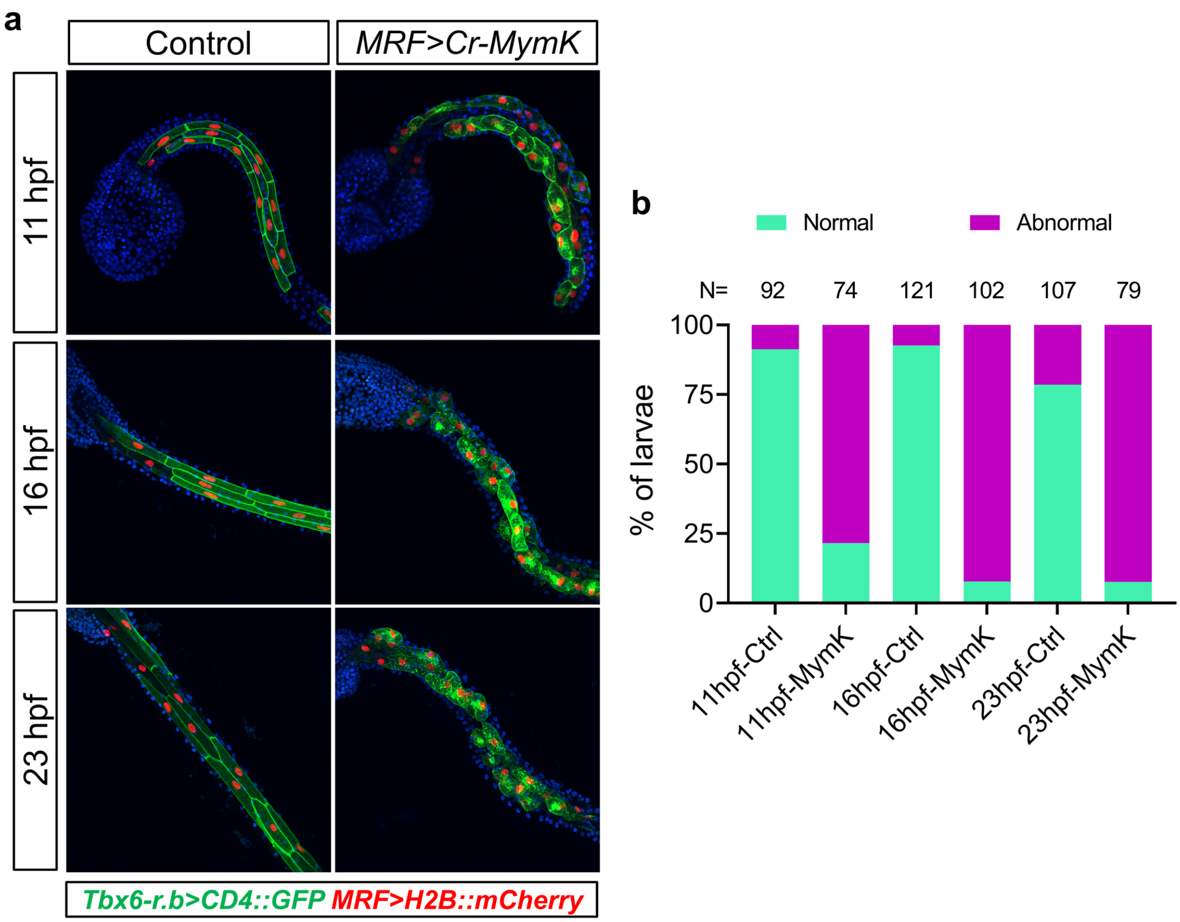


**Extended Data Fig. 9: Overexpression of MymK in mononucleated tail cells perturbs morphology but does not induce cell fusion.**

**a**, Comparison of mononucleated tail muscle cell morphology between negative control embryos/larvae and those overexpressing *Ciona robusta* MymK driven by the MRF promoter (*MRF>Cr-MymK*). Cells lost their characteristic shape and cell-cell junctions, but did not show ectopic bi- or multi-nucleation. Cells were labeled with a combination of *Tbx6-r.b>CD4::GFP* (green) and *MRF>H2B::mCherry* (red) reporter plasmids and counterstained with DAPI (blue). All images are projections of Z stacks acquired on a scanning confocal microscope. **b**, Scoring of "normal" vs. "abnormal" tail muscle morphology in embryos/larvae represented by images in panel **a**. Individuals were scored under confocal microscope as showing "abnormal" morphology if over 50% of labelled tail muscle cells show rounded or other irregular shape other than the regular polygonal shape observed in majority of negative control cells. N, numbers of larvae assayed for each condition.

Extended Data Fig. 10

```
Sea lamprey      MPVLLLVRALALGLVSRAAGAARALCGSGGAARSALLSCVLCLVDTKHRGGDDDD---GD 57
Arctic lamprey   ***************************************************G***DGG** 60

Sea lamprey      DDDRHGDDGDDKNA--RRSNRKDGDLGKSQGDARRESRSATARRAVRTRRFHDDDDDG-- 113
Arctic lamprey   ******V*******SR*******V********D***********************GDGA 120

Sea lamprey      ADDDEDVEWSGDETPPRTLSRTRARRAGTSERLRRSRRNDEDD-DDDDYDGGGRGDEGDS 172
Arctic lamprey   D*****A************************************D*******R*******C 180

Sea lamprey      EKKSSSRAGLARDALALALAFAVESKIRAKATRKDERRRREEEGEER-GEMEGRGDREGV 231
Arctic lamprey   ***********************************************G************ 240

Sea lamprey      QMRHKRDTQIPRGHRPPRKLSYDDD-DNEDIVDSDGGG-GGGGREEEVTPSSKSRVNSSS 289
Arctic lamprey   ******E******************G************G********************* 300

Sea lamprey      SRSGFARDALSLALSYALTSTDLGSSPGKARNSATARGDRNSGGRAFTPEKGTPLPVPPR 349
Arctic lamprey   *********************E************************************** 360

Sea lamprey      GVASVKRRLFQDDADGDGDDGPDGVRVAAGGCGDVVDCGRGGDGGIRGGGGGREFGKGSG 409
Arctic lamprey   *************G*******R*****************************S****R*** 420

Sea lamprey      DAGVIRGGRDGAGGRGSGVIRGCIIGGTDGGVGGSGNGGGTRCGSNDRGVSSSSSSRLQP 469
Arctic lamprey   *V*****************T*************A*******************-****** 479

Sea lamprey      PPPPSPPPPPPPPCQPP-PPHPRPPVLAHAGPRPGRNGSRSLQLDTTTT-TRGGGSSGRN 527
Arctic lamprey   *****************P**********R**************F***I*T*TTK****** 539

Sea lamprey      ERPRSAGCPPVRGPVSSFQTRTLPVGLLLEAKRGRKDLARIGTAVQASSRLRGRCK* 583
Arctic lamprey   ******************************T************************** 595
```

**Extended Data Fig. 10: Conservation of MymX protein sequences in lamprey species.**

Alignment of protein sequences encoded by *MymX* genes identified in sea lamprey and arctic lamprey. Identical amino acids are shown as asterisks. AxLyCxL motif is highlighted.

Extended Data Fig. 11

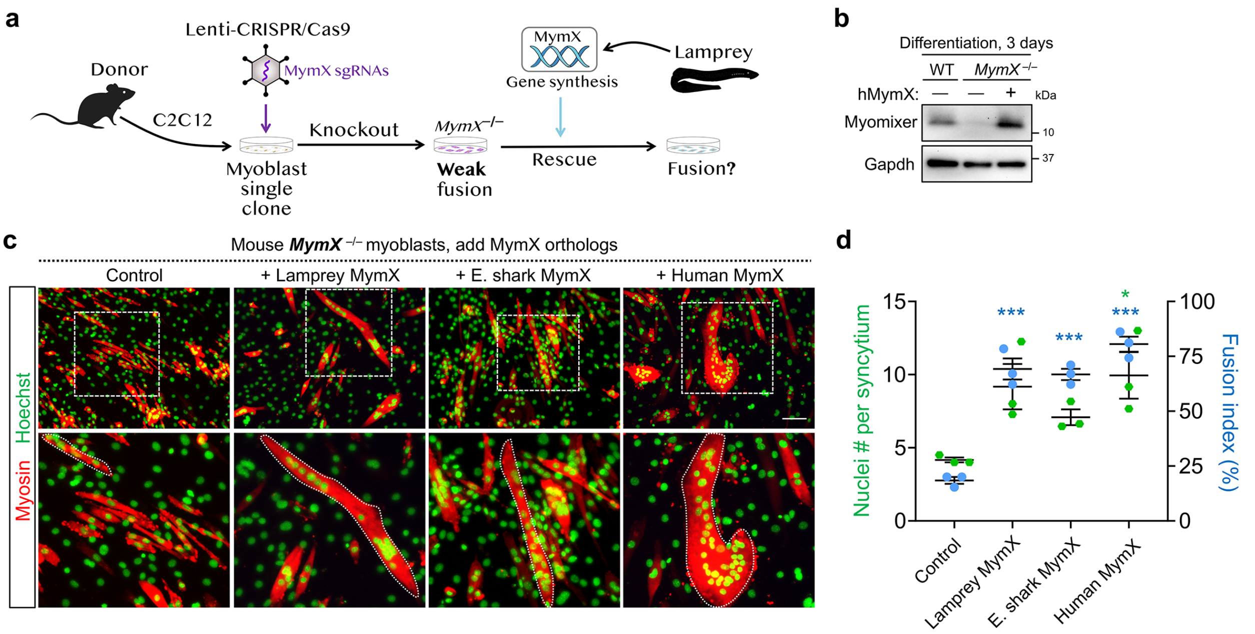


**Extended Data Fig. 11: Lamprey MymX protein can rescue the fusion of mouse *MymX*$^{-/-}$ myoblasts.** **a**, Schematic of experimental design to generate mouse *MymX*-deficient myoblasts by CRISPR/Cas9 and test the fusogenic function of lamprey MymX proteins. A pair of sgRNAs that target the coding exon is applied. **b**, Western blots validating the successful depletions of MymX protein in an isolated single clone of CRISPR treated mouse myoblasts. **c**, Myosin immunostaining of mouse *MymX*$^{-/-}$ myoblasts transfected with control (empty vector) or MymX ortholog expression vectors. Sea lamprey MymX protein can induce myotube formations (outlined). E. shark: elephant shark. Scale bar, 100 µm. **d**, Measurements of myoblast fusion in **c** after 5 days of differentiation. Data are means ± SEM. * $P < 0.05$; *** $P < 0.001$, compared to control group, one-way ANOVA.

Extended Data Fig. 12

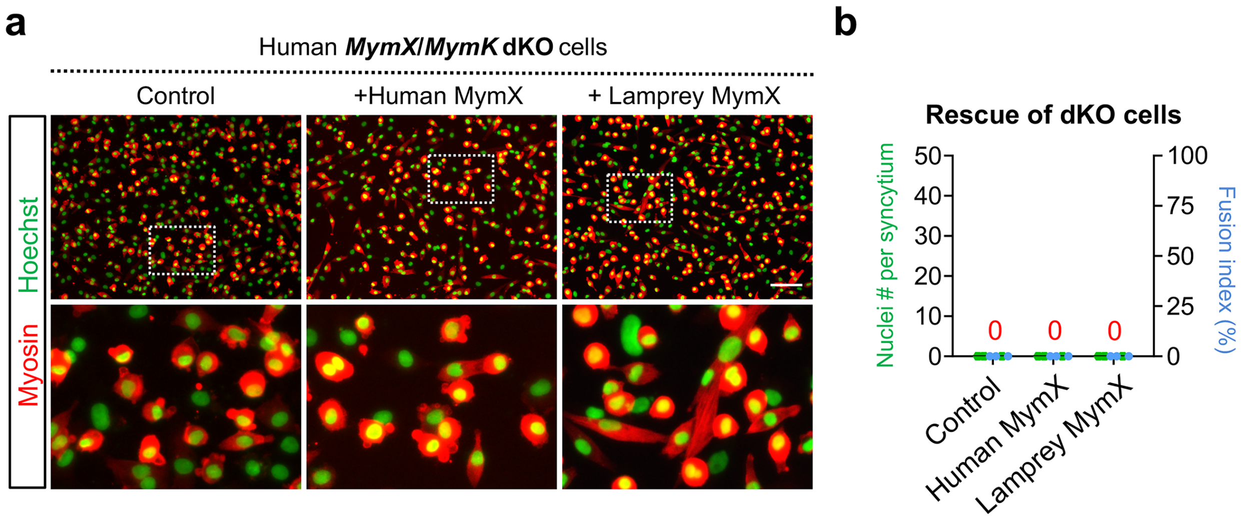


**Extended Data Fig. 12: Lamprey MymX protein requires MymK for function.**

**a**, Myosin immunostaining of human *MymK*/*MymX* dKO myoblasts transfected with human or lamprey MymX expression vector. Scale bar, 100 µm. **b**, Measurements of myoblast fusion in **a** after 5 days of differentiation. Data are means ± SEM.

Extended Data Fig. 13

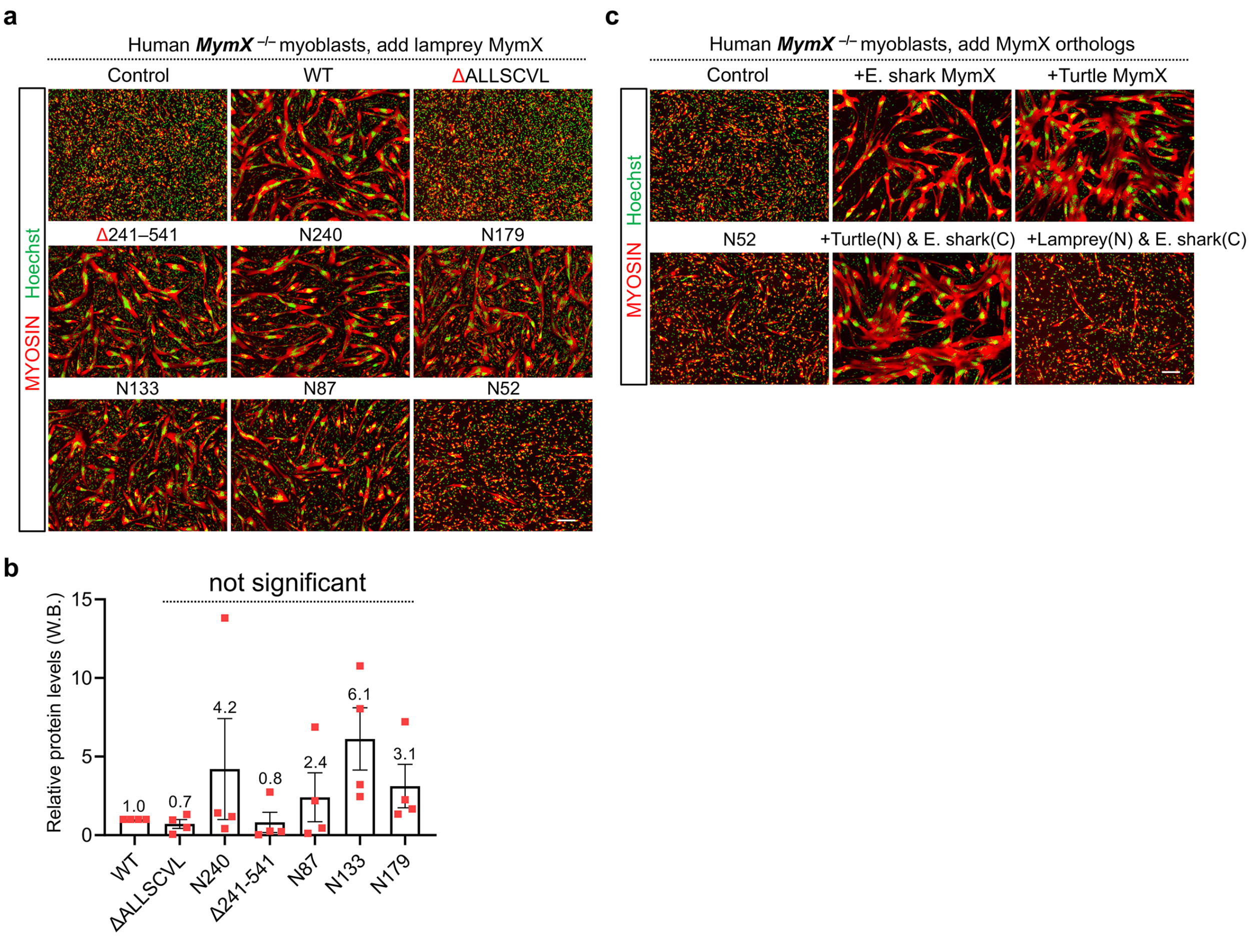


**Extended Data Fig. 13: Structure and function analyses of lamprey MymX mutants.**

**a**, **c**, Macroscopic views of myosin immunostaining of human *MymX* $^{-/-}$ myoblasts transfected with MymX expression or control vector. Cells are differentiated for 4 days. Scale bars, 200 µm. **b**, Measurement of lamprey MymX protein expression levels in human myoblasts by analyzing band intensity from Western blot. Numbers are the average fold-changes normalized to the expression level of wild-type (WT) lamprey MymX protein. Data are means ± SEM. n = 4, compared to WT, one-way ANOVA.

Extended Data Fig. 14

| Clade | Species | Coding Exon 1 (bp) | Coding Exon 2 (bp) | Coding Exon 3 (bp) | Coding Exon 4 (bp) | Coding Exon5 (bp) | ORF (a.a.) | Protein identity vs. human MymK |
|---|---|---|---|---|---|---|---|---|
| Jawed vertebrate | Human | 135 (3N) | 115 (3N+1) | 149 (2+3N) | 117 (3N) | 147 (3N) | 221 | 100% |
| | Zebrafish | | | | | 144 (3N) | 220 | 75.2% |
| | Elephant shark | | | | | 144 (3N) | 220 | 74.8% |
| Jawless vertebrate | Arctic lamprey | 135 (3N) | 115 (3N+1) | 149 (2+3N) | 117 (3N) | 150 (3N) | 222 | 56.1% |
| | Sea lamprey | | | | | 144 (3N) | 220 | 55.5% |
| Ascidian (tunicates) | *P. mammillata* | 135 (3N) | 146 (3N+2) | 121 (1+3N) | 122 (3N+2) | 151 (1+3N) | 225 | 38.1% |
| | *C. robusta* | | | | | 133 (1+3N) | 219 | 36.6% |
| | *H. aurantium* | | | | | 151 (1+3N) | 225 | 36.3% |
| | *H. roretzi* | | | | | 151 (1+3N) | 225 | 35.8% |
| | *B. schlosseri* | | | | | 151 (1+3N) | 225 | 35.1% |
| | *B. leachii* | | | | | 151 (1+3N) | 225 | 34.2% |
| | *M. occulta* | | | | | 160 (1+3N) | 228 | 34.1% |
| | *S. clava* | | | | | 151 (1+3N) | 225 | 33.8% |
| | *C. inflata* | 135 (3N) | 146 (3N+2) | 121 (1+3N) | 122 (3N+2) | 136 (1+3N) | 220 | 33.6% |
| Thaliacean (tunicates) | *P. verticillata* | 135 (3N) | 146 (3N+2) | 121 (1+3N) | 122 (3N+2) | 151 (1+3N) | 225 | 34.9% |
| | *S. thompsoni* | 138 (3N) | 158 (3N+2) | 121 (1+3N) | 122 (3N+2) | 163 (1+3N) | 234 | 26.5% |

Nucleotide number and complementarity of exon: 3N, 3N+1, 2+3N, 3N+2, 1+3N

**Extended Data Fig. 14: Exon size and structure for various MymK orthologs.** Exon structures for MymK orthologs are conserved in the tunicate and vertebrate species groups.

Extended Data Fig. 15

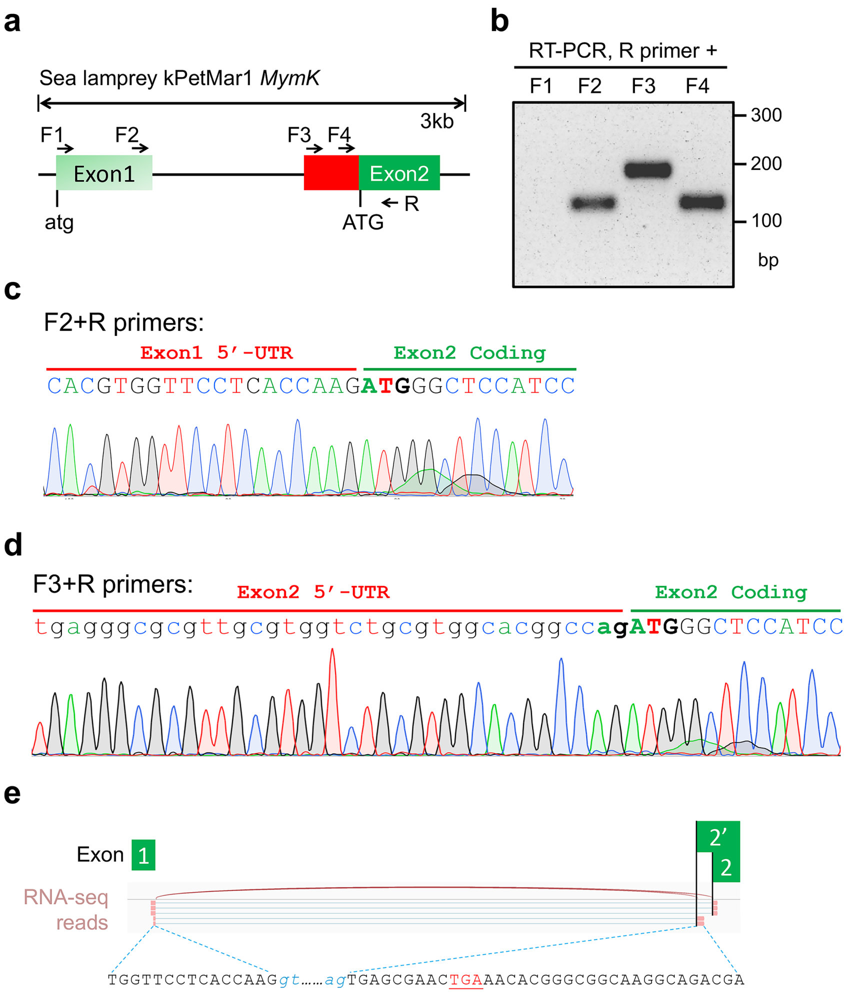


**Extended Data Fig. 15: Transcript isoforms of lamprey *MymK* gene.**

**a**, Schematic of lamprey *MymK* exons and the positions of RT-PCR primers. **b**, Gel electrophoresis results of RT-PCR amplification of lamprey MymK mRNA using various combinations of forward primers and universal reverse primer. **c**, **d**, Sanger sequencing results of RT-PCR products in **b**. These results validate the 5’ UTRs (untranslated region) for two splicing isoforms. The 3′ splice acceptor site (ag) is highlighted. **e**, RNA-seq read (SRA accession: SRR9958780) reveals a cryptic splicing site by which *MymK* expresses the shorter ORF (220 a.a.) in neural crest of lamprey embryos. Splice donor and acceptor sites are highlighted in blue.

Extended Data Fig. 16

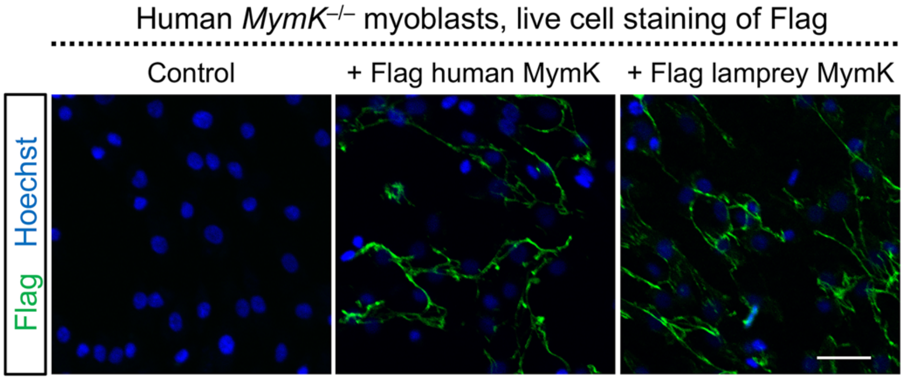


**Extended Data Fig. 16: Confirmation of the plasma membrane expression of lamprey MymK.** Immunostaining of Flag epitope on live myoblasts after transfection with control (empty vector) or Flag-MymK expressing vectors. Flag-MymK constructs contain a synthetic cleavable signal sequence upstream of Flag epitope followed by the full-length MymK protein. By the topology of MymK protein, Flag epitope is located in the extracellular space thus accessible to antibody staining of live cells (without fixation or permeabilization). Scale bar, 40 µm.

Extended Data Fig. 17

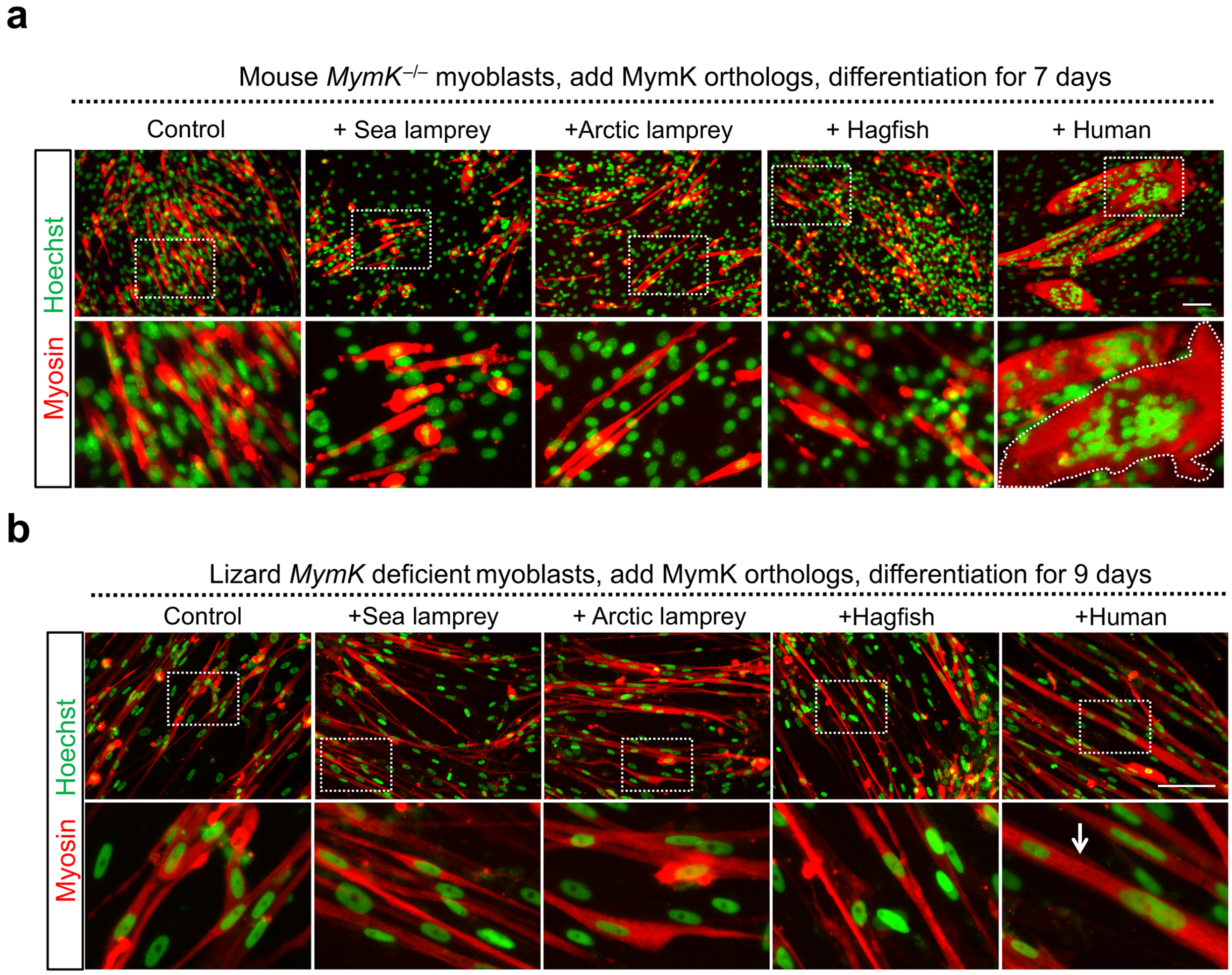


**Extended Data Fig. 17: Cyclostome MymK cannot induce fusion of mouse and lizard *MymK*$^{-/-}$ myoblasts.**

**a**, **b**, Myosin immunostaining of mouse (**a**) and lizard (**b**) *MymK*$^{-/-}$ myoblasts transfected with cyclostome or human MymK orthologs. Cells are differentiated for 7 days for **a** and 9 days for **b**. Scale bars, 100 µm.

Extended Data Fig. 18

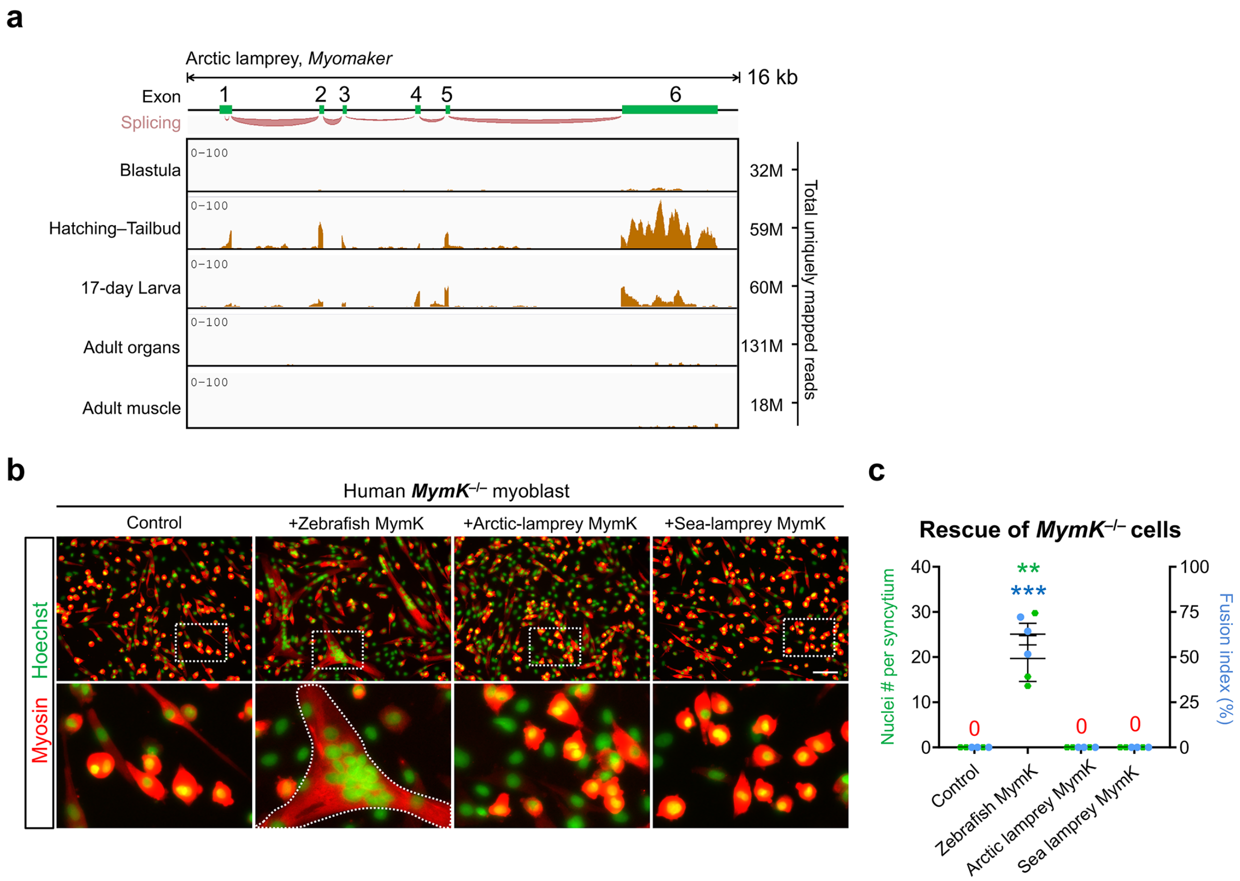


**Extended Data Fig. 18: Arctic lamprey MymK protein is not able to activate fusion of human *MymK*$^{-/-}$ myoblasts.**

**a**, Arctic lamprey *MymK* gene structure and RNA sequencing tracks that show the expression levels of this gene in arctic lamprey embryos (SRA accession: PRJNA371391), whole larva (17 day old, SRA accession: PRJNA553689) and adult organs (notochord, ovary, testis, kidney and heart, SRA accession: PRJNA354821). Arctic lamprey MymK protein contains 222 amino acids that are highly conserved (97% identical to sea lamprey MymK). **b**, Myosin immunostaining of human *MymK*$^{-/-}$ myoblasts transfected with MymK orthologs. Muscle syncytium is outlined. Scale bar, 100 µm. **c**, Measurement of myoblast fusion for groups in **b** after 5 days of differentiation. Data are means ± SEM. ** $P < 0.01$; *** $P < 0.001$, compared to control group, one-way ANOVA.

Extended Data Fig. 19

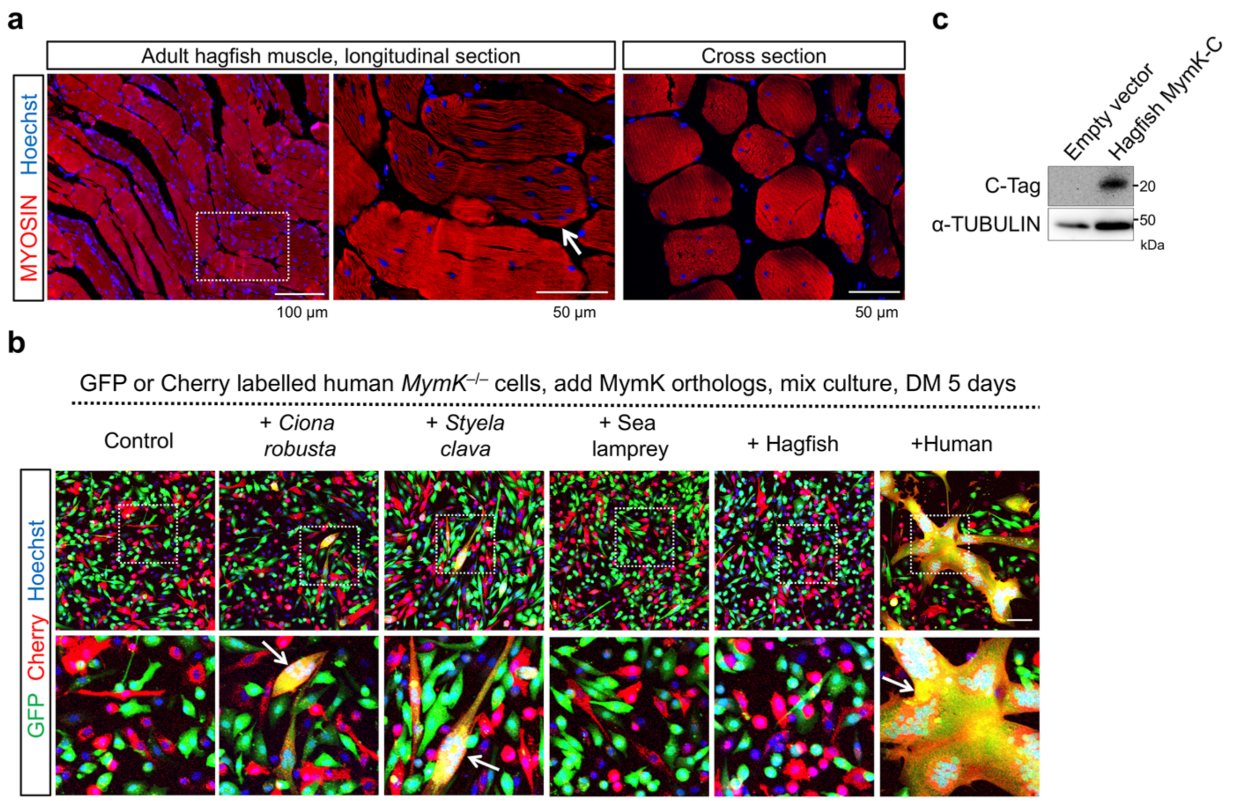


**Extended Data Fig. 19: Hagfish MymK protein fails to activate fusion of human *MymK*$^{-/-}$ myoblasts.**

**a**, Myosin immunostaining of muscle tissues dissected from adult hagfish (*Myxine glutinosa*). Arrow points to a multinucleated myofiber. **b**, Immunofluorescence of human *MymK*$^{-/-}$ myoblasts separately labelled by GFP or Cherry fluorescent proteins and later transfected with MymK orthologs before the mixing culture in differentiation medium for 5 days. Dual-color labelled myotubes (arrows) indicates the multinucleation through fusion. Lamprey and hagfish MymK proteins cannot induce human myoblast fusion. Scale bar, 100 µm. **c**, Western blot results that confirmed expression of hagfish MymK protein (C-tagged) in human myoblasts. The predicted molecular weight for the fusion protein is 25.2 kDa.

Extended Data Fig. 20

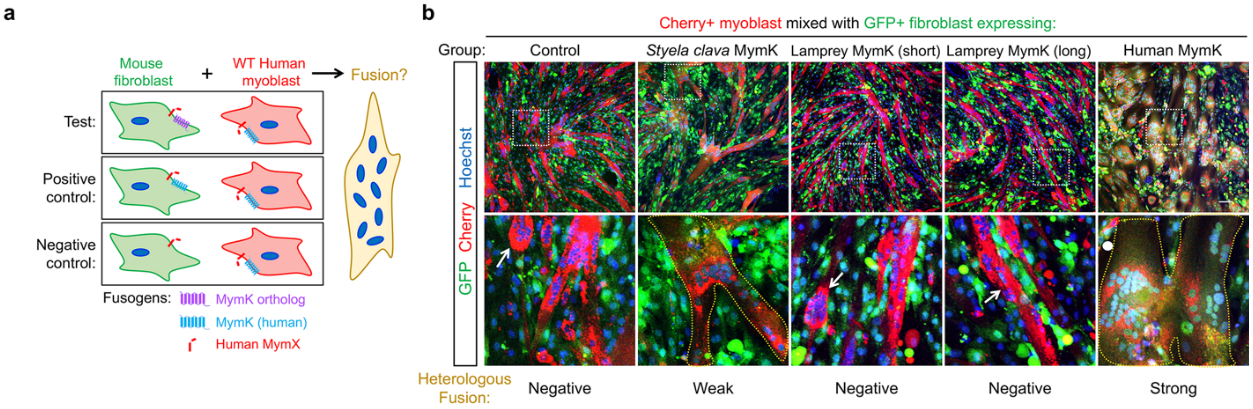


**Extended Data Fig. 20: Lamprey MymK proteins fail to induce fibroblast–myoblast fusion.**

Schematic (**a**) and results (**b**) of cell labeling and mixing experiments that examine the function of MymK orthologs in mouse fibroblasts. Fibroblast-myoblast syncytium is outlined. Arrows point to myoblast-myoblast syncytia. In contrast to sea lamprey MymK, MymK proteins from tunicate (*Styela clava*) and human induced heterologous cell fusion. Cells were differentiated for 7 days after mixing. Scale bar, 100 µm.

Extended Data Fig. 21

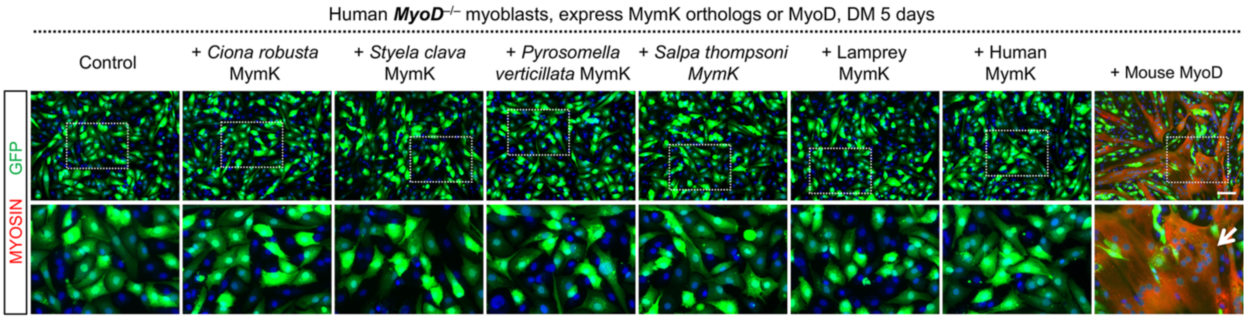


**Extended Data Fig. 21: Fusogenic activity of MymK proteins in human myoblasts depends on the master regulator of myogenesis, MyoD.**

Myosin immunostaining of GFP labelled human *MyoD*$^{-/-}$ myoblasts. Deletion of *MyoD* gene completely abolished myogenic (myosin expression) and fusogenic programs of human myoblasts. Such defects can only be rescued by re-expression of MyoD protein but not MymK orthologs, suggesting the fusogenic activity of MymK proteins depends on the expression of MyoD-responsive gene(s) or permissive factor(s). Note the endogenous expression of *MymK* and *MymX* is abolished in *MyoD*$^{-/-}$ myoblasts[12]. GFP expression was utilized to view cell body. Cells are differentiated for 5 days. Scale bar, 100 µm.

Extended Data Fig. 22

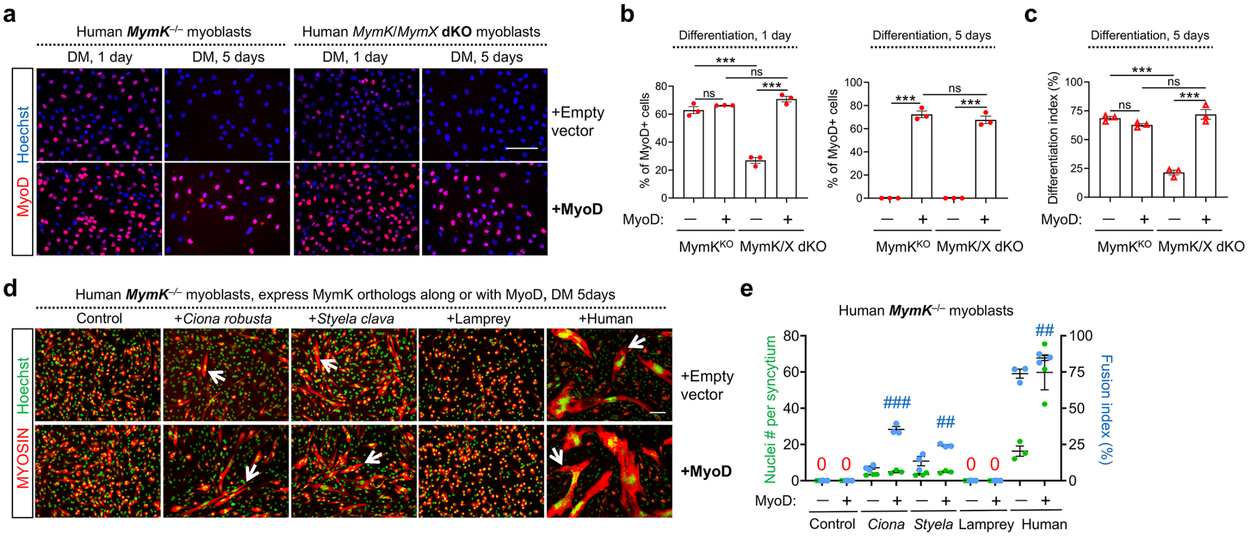


**Extended Data Fig. 22: Prolonged expression of MyoD during human myoblast differentiation boosts the fusogenic activity of MymK orthologs.**

**a**, MyoD immunostaining of human *MymK*$^{-/-}$ and *MymK*/*MymX* dKO myoblasts at different timepoints (day-1 and day-5) of differentiation under normal (top) or MyoD overexpression (bottom) conditions. DM, differentiation medium. MyoD is normally expressed in early (1 day) but not late stage (5 days) differentiation. In addition, the percentage of MyoD$^{+}$ cells is lower in *MymK*/*MymX* dKO compared with *MymK*$^{-/-}$ myoblasts (upper panels), reflecting an inherent difference of myogenic potential in the clonally isolated mutant cells. Such a difference can be normalized by forced expression of mouse MyoD protein (lower panels). Note the endogenous expression of *MyoD* is not affected by deletion of *MymX* or (and) *MymK* gene per se[12]. **b**, Quantification of MyoD staining in **a** after 1 day (left) or 5 days (right) of differentiation. **c**, Measurement of myogenic differentiation in absence or presence of MyoD overexpression after 5 days of differentiation. **d**, **e**, Myosin immunostaining of human *MymK*$^{-/-}$ myoblasts transfected with MymK orthologs individually (upper panels) or together with mouse MyoD protein (lower panels). Muscle syncytia are pointed by arrows. MyoD promoted myoblast fusion when tunicate or human MymK protein is co-expressed, yet co-expression of MyoD with sea lamprey MymK failed to induce fusion. Cells are differentiated for 5 days. Scale bars, 100 µm. Data are means ± SEM. ***, ### $P < 0.001$; ## $P < 0.01$; # $P < 0.05$. * compared to control group, one-way ANOVA; # comparing effect of MyoD expression, one-way ANOVA.

Extended Data Fig. 23

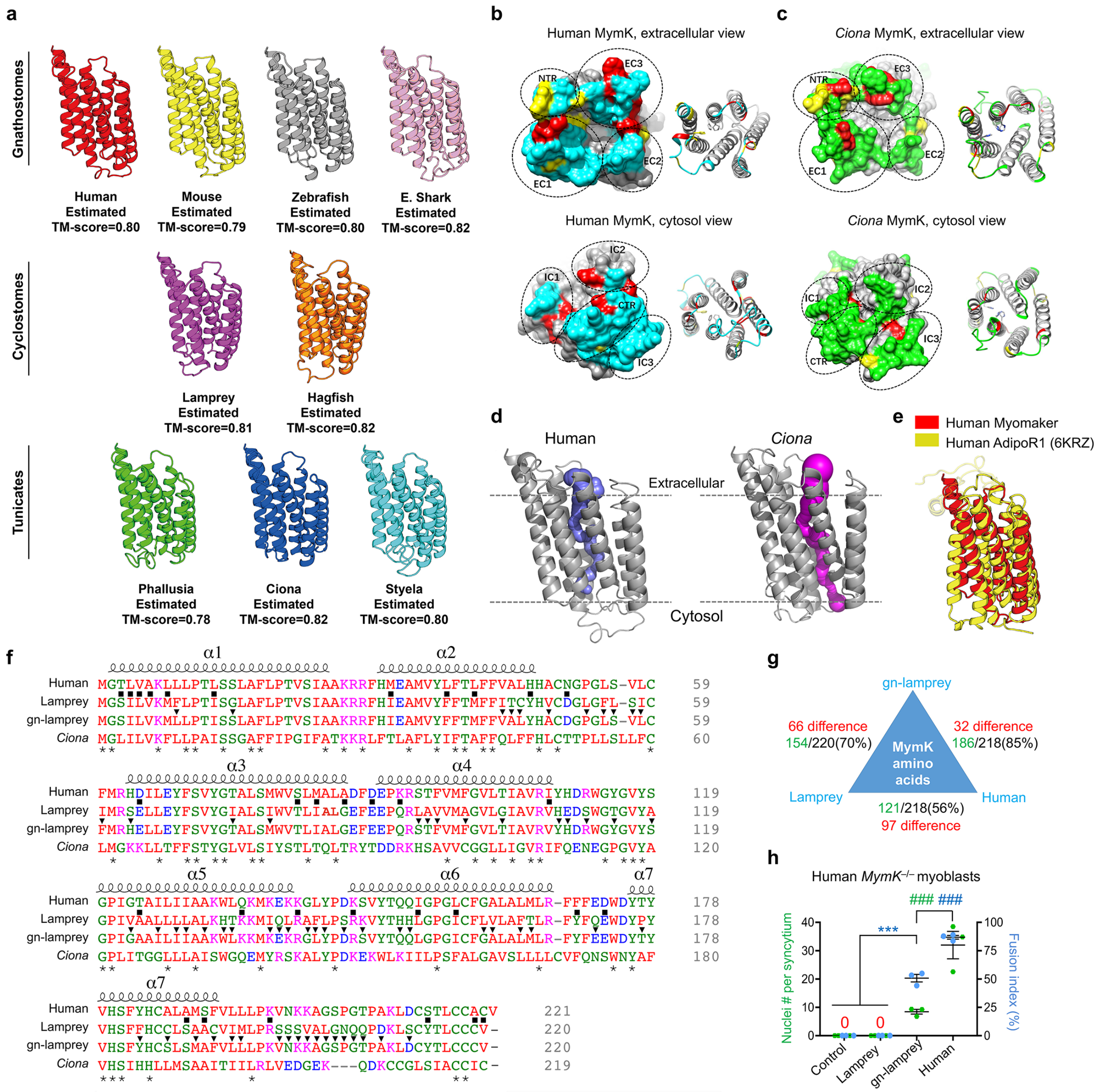


**Extended Data Fig. 23: Structural modeling of MymK proteins.**

**a**, Protein structural models for MymK from jawed vertebrates (top), jawless vertebrates (middle) and tunicates (bottom). TM-score: template modelling score. **b**, The extracellular (top) and intracellular (bottom) faces of human MymK protein model shown by surface (left) and cartoon (right) representations. Cyan highlights the surface residues; red highlights the conserved residues among vertebrates; yellow highlights the conserved residues among all species. **c**, The extracellular (top) and intracellular (bottom) faces of *Ciona* MymK protein model shown by surface (left) and cartoon (right) representations. Green highlights the surface residues; red highlights the conserved residues among tunicates; yellow highlights the conserved residues among all species. EC: extracellular; IC: intracellular; NTR: N-terminus region; CTR:C-terminus region. **d**, Side view of the predicted cavities inside MymK proteins. **e**, Superimposition of human MymK and human adiponectin receptor 1 (AdipoR1, PDB ID: 6KRZ). **f**, Sequence alignment of MymK proteins. Arrowheads indicate different residues between gn-lamprey MymK and native lamprey

MymK; squares indicate different residues between human MymK and gn-lamprey MymK. **g**, Comparisons of human MymK protein sequence identity with that of lamprey MymK before and after (gn-lamprey) mutation. **h**, Measurement of myoblast fusion for the experiment in Fig. 7g. Data are means ± SEM. *** $P < 0.001$, compared to control group, one-way ANOVA. $^{\#\#\#}$ $P < 0.001$ for comparison between gn-lamprey and human MymK groups, one-way ANOVA.
