## Supplemental File 5 for "Evolution of a chordate-specific mechanism for myoblast fusion"

**Electroporation mixes (all per 700 µl of cuvette volume):**

MymK CRISPR
140 µg MRF-2685/-1>CD4::GFP
25 µg MRF-2685/-1>H2B::mCherry
40 µg Mesp-1916/-1>Cas9
40 µg U6>MymK.2.6
40 µg U6>MymK.4.68

Negative Control (for MymK CRISPR)
140 µg MRF-2685/-1>CD4::GFP
25 µg MRF-2685/-1>H2B::mCherry
40 µg M-1>Cas9

MRF>MymK (larval tail overexpression)
70 µg MRF-2685/-1>Ciona robusta MymK
50 µg Tbx6-r.b-1249/+12>CD4::GFP
10 µg MRF-2685/-1>H2B::mCherry

Negative Control for larval tail overexpression
50 µg Tbx6-r.b-1249/+12>CD4::GFP
10 µg MRF-2685/-1>H2B::mCherry

MymK-509/-1>GFP
140 µg MymK-509/-1>Unc-76::GFP

**Novel, unpublished sequences from this study:**

sgRNA target sequences validated by TIDE (N19+**PAM**):
MymK.2.6: CAATGGAGTTGTGCAGAGG**TGG**
MymK.4.68: CAGCTCCAAGTGCAAACGA**TGG**

Peakshift assay negative control sgRNA sequence (N19+**PAM**):
Gsx.4: ACGACAGTGACGAAAGGTC**CGG**

sgRNAs rejected due to insufficient evidence of activity (N19+**PAM**):
MymK.2.22: AAAACAGAAGAGACAACAA**TGG**
MymK.3.65: GTGGACCAGCATAAACACC**AGG**
MymK.3.97: CTGGTCCACTAATAACTGG**CGG**
MymK.4.24: CAAGAAATGTACCGATCCA**AGG**

>C. robusta MymK cDNA for riboprobes and overexpression START STOP T7
ATGGGTTTGATTTTAGTGAAATTTCTACTCCCTGCAATTAGTAGTGGCGCTTTTTTTATACCAGGAATCT
TTGCTACGAAGAAAAGGCTTTTCACGTTAGCATTTCTTTACATTTTCACTGCTTTCTTCCAACTGTTTTT
CCACCTCTGCACAACTCCATTGTTGTCTCTTCTGTTTTGCTTGATGGGAAAAAAGCTTTTGACATTCTTC
TCCACGTATGGACTAGTGCTATCGATATACTCCACACTTACACAATTAACAAGATATACCGATGATCGAA
AGCACTCAGCTGTGGTATGTGGTGGCTTGCTTATTGGCGTGAGGATATTCCAGGAAAATGAAGGACCTGG
TGTTTATGCTGGTCCACTAATAACTGGCGGCCTACTGCTTGCGATATCCTGGGGTCAAGAAATGTACCGA

TCCAAGGCTTTATATCCGGATAAAGAGAAATGGTTGAAGATAATTTTACCATCGTTTGCACTTGGAGCTG
TTTCTCTACTTCTACTTTGTGTTTTTCAAAACAGTTGGAATTATGCTTTTGTTCATTCCATCCACCACTT
ATTAATGTCAGCTGCCATCACAATTATCCTTCGGCTCGTGGAAGACGGAGAGAAGCAAGATAAATGTTGT
GGTTTATCCATTGCATGTTGTATATGTTGAgaattcCCTATAGTGAGTCGTATTA

>C. robusta MymK gene model, based on HT genome assembly[1]
EXONS START STOP (UTRs not determined)
ATGGGTTTGATTTTAGTGAAATTTCTACTCCCTGCAATTAGTAGTGGCGCTTTTTTTATACCAGGAATCT
TTGCTACGAAGAAAAGGCTTTTCACGTTAGCATTTCTTTACATTTTCACTGCTTTCTTCCAACTGgtaag
tagattactttagaaattacaatagggcagaattaagtccaacgaatagtgttctataacagatatattt
ttcaccatttttattattatcttgtttttgattatgttgtatagtaaggtgggggaagatgggacacct
ttagcacgtaatatccaaatatcctgatcgtcctttaaataacaacggtctaagggagtcgtaagaatat
aggtaattttataattcattgtatttccttggtttattcctaagtctgacaaaaaatagagttaaaaggt
ttcctatcttctcccaccctactgtatagtacaatggggggagatgggacacttaagcacatattgccaa
atattgccaaaatcaaaatagatgacggttttgggtaataccacgaattagtaccatcattcctttatta
attgacatcaatttaataaaatataataaaacaaacgaggagttatttagcgatccgcacaacaggtgaa
attttacaaatttgaaaacacacaggataagatgggacactttaaaatcattgttaattttgattattaa
gtgtatttataaatatgaatatatgaaactagtttaatcgttcacacacgcaaactaaaaaattcattgg
ctttactatgattcagcgagcattaaaaaatagcatcttactatttcgcatttttttagctgttcaaagta
tactgtttctatttacatattttttaatatttaaaaacagtactcagctttgacggacgttttataaa
gtcgtgataatactgtcgaataattctgtaaccttattttcctaattatcattaaatgggggtccgtaaa
aagaaaattaaaaaaagtcttgtctgacaaagtgtcccatcttccaaagtttacaataaacagtttcgag
tgcacacttatgtaggatttagaatcaagtgcattacattaatacacatcacagtataaataaacattgg
aatagaaaataaaacaaaaaaaaacaatcataataacaaatagataagaaacaaaatggaaaaggggcag
caagtataatttcggatttgttgcgaattgtttgaatttgtccattgttgttgtggatcaatccttgtga
tgcctggagctcgggttcatgaagggttccttcgattttggtttacacctgcaataaagtatttgtgggc
agtaaatctgtttaaggtttgctttggaattgaaaagttaggaagaataagatgaaaaagtaaggaaaaa
atatataaacaaatattaaaaagaagagacaatcaacagaattgaataaataaaaaaaaaaataaaaaaa
tatatataaaaaatgagcaagactgagtgtcatgcatatgtgtgttatataacaataatgatgtgagcta
caaaaatagtagcttgatggacagagttctattagggccggcacaaggataattaggaaattaaacaaac
aaaacacaagtgtgccaacggcttgccaagtaatagtactcatgtcggtgtgaacactgaaataatgtgc
aacaaattatggtgatagtgaatgtacatgctgcaataaagtaaaagtcggaccatgcaaatgatacaaa
agtaaaaaattgggagttatggtgtttcacagtggtcaatttaaaataaatcaaagtataaagtgcaaca
atgtaaaaatagtgctgtgtgacattgtagttatttataaaggaaaactttaaagcacaatattatgtgg
tttgacatagaaaagagagccagagggttttgggttaaagggaagctataaacatactgtttacgaatcc
attctatacctactatactatatcataatatactgaagtgagtatactattactacgcgatcacaaatga
cgaaacctaggctactttgctagttatatatctcttatttgtttttgataaagcattttccgccgtggta
aggagagagttgttaattgttacgtttaaaccgcagTTTTTCCACCTCTGCACAACTCCATTGTTGTCTC
TTCTGTTTTGCTTGATGGAAAAAAGCTTTTGACATTCTTCTCCACGTATGGACTAGTGCTATCGTTATA
CTCCACACTTACACAATTAACAAGATATACCGATGATCGAAAgtaagttgttagttctagggaaaaatat
aacttattttactccagggtggcgggacaacgacagtcgttataaaacggggggttctgtttcatacacct
cgtataccttcttactaattaacccgtttatgtattttaagggtgatttagtatagactgtagtaatcga
ttgtttttatttgctgctaccactgtaggtggtccacagagttacggttatatagttctaaaaatgttct
tttattacgaaattgggcaagaataaggggaatgaaaatttgcccatatatccccactaccatacagaat
tgtagaagcgaaacattgtactttcgcttagcccaatttttgaaataaccgcattacagGCACTCAGCTGT
GGTATGTGGTGGCTTGCTTATTGGCGTGAGGATATTCCAGGATGAAGGACCTGGTGTTTATGCTGGTCCA
CTAATAACTGGCGGCCTACTGCTTGCGATATCCTGGGTTTGTAGCAAATTTTTGGTTTCTACGTTTGAAT
GTTCAATGGCGATTTTTGTACATAAAGGAGACCACGTCACGTATTTTAACCCGGCCGGTGGCACAATAGT

TAGCGCACCGGCATCTGGTTCAGAGgttcagacacgcaaaatcgtgaagcgttccatatttttaaaaaa
aggttcctgaaaattaacgccaatattgctcccagtaattacttgtgagctgtttaaattgttagctata
aatacagaaacaataaccctacaaaaaaacgattttacctaccaaaaggcgggcataaggtgctgtatga
aacagattacccgtattataatgtcgtttccccgccacgcgagtataaggcacgcaagttacattcattc
gtttattatcgcgtttcaaacacagGGTCAAGAAATGTACCGATCCAAGGCTTTATATCCAGATAAAGAG
AAATGGTTGAAGATAATTTTACCATCGTTTGCACTTGGAGCTGTTTCTCTACTTCTACTTTGTGTTTTTC
AAAACAGgtaaaacgttatacttttattgatcggactattgacgctgtatgattttaacaacatcaatat
gtttaatgtagaactataccaatgcttataaaacctagcatgttgtaattgcgggatattattctctaaa
cagTTGGAATTATGCTTTTGTTCATTCCATCCACCACTTATTAATGTCAGCTGCCATCACAATTATCCTT
CGGCTCGTGGAAGACGGAGAGAAGCAAGATAAATGTTGTGGTTTATCCATTGCATGTTGTATATGTTGA

>MymK(-509/-1) promoter
TGCTCTGGAAAATTTACCAAGGGAAACTCCCTCACGTGGTAACATAGAAATAGTACAAAAAATGCTTAAA
ATTGTTTTTCACCAAAACTAACGGTACACATACCGTTGCCTAAACTAGGCAAAACACATATTAGATTGCA
TTTAAACTGGTAAATGTCCTAATAATTTAAAACTTGCATTAACAATAAATTTAGTTGCCAATAATATAAC
TTTTAAATTGCGTATTTTAAGTATGTACATAGTTAAGAATATTTATTCGGGCCATAAATGTGTATTTTAA
AGTTGTTTATATTTAGACCAAATACTGATATTTTAATAAACGTTTTGGGTCGTGAGAAATATCACATTAT
TGTGTCCAGCTGAATCAGACAACAGCTGAAGACCACATGCCAATTCCCAAGATAATGGTGCAGCACGCGC
AAAACAACCGAGAGTACCGTAGTACATTTGGTATGTCGCCTTTAGAGCATTCATTGCATAGTGTTAAACC
GTAGATTAAATCATAGTT

>MRF(-2685/-1) promoter
gcaagctcctttggggtttggccactcgatcggcaagcgaaatatcatttaagtttttaaatgtcgttta
aatggcagcagccgtaacaggtggactgccacgtgtcgcccggtgtgatcgtgtttctatcccaagggtg
gcagtgcaggcatgccgacgctcatatgagccattcataataatttggcgcatgtaagaggaaaatgcac
cagtaagatggtgccagcacattgtaattaaaatgtggtgctacggggtaagatggaaatcgttaggaaa
taaaatcccttattcccaatcgcttttaccaattaaccacgttattttatatttgcgaagttacggttac
ataattatagtagggtgggggaagatggaacaactttagcacaaaatatataaatatcctggtcgtgttt
taaacaattaacaatggtctatgggagttgcgaggatacggttttatatttctttgaatgttctttgttt
actaccaaaatgtacgagaaaatagaatgaaaaggtgtcccatcttcccctatcctactatatgtaaata
ctcctttttttgctaccaaacgggacgagaaaagaaattaaagcatgaaccattttacttcaacctactac
tacagtatctaagtaatttcttttcgaaacagatttaacttctttgcgatttcaattagcgaatcccagt
ttgaccgttaaagctatagcttagagtacgtggtgctaaatttcgttgattcgcgcccctgaagcgagca
acttattagatagaaatcccagctggtaacaacatgtgttcatcgccctactgatgtctgtggttgccga
actctctttacgagactgagagccgacgtctcgataaacgaggttgtctattatgacttatttaggaaat
aaaaagtaagaaaataaaaatttaattaagtaaaatacaaacgtttacaaaaattataattttttttcac
caattttaattttttttcaagtatccgacgacgtttaagaagtttaaaatccaaatcctttgctgcttcc
tagaattatactgtggggtaagatggaaaccgtaaatattctgtgtttacaaccaaataggacgataaaa
gagaatgaaacgtggacaatttgccctaacacattacacggctaaaaatattaggtttcctcttaaaata
gcagcgacgaccataaggttattcgaagatgaaaggccaaagtaacggatgcaagtttctgcaagacaat
gctgtaattgttacgtcacagaggaattccacaacttattccagagcagacgagcgtcgcgtttactgtt
tgctaaaaatggcgactcgaggtgacaattacaattcgcgctgcgaatgacgtaataatgctattcgctg
tgatgaaataatagaagtggaaattcgaacaataccaacccacgatgaagcatatgtagcgaatccggaa
ttaaacctctagacttgcttgggaaatgcatggcgtatatgagtgtattacgtgatcgttaaagcgtctt
atacagtcgcattagaagctattttaagaaaatccacggcccagcatgcaaatatagctgtgatctaata
ggttaggttagtttacattaagtttaataatgtcgatcacttattgcatgtttcatgatcttgcaaatat
gatgacgttgtttattgaggttgctgctttcagttttatacaatcttccaacaagttaaaactggcgaat
ttagacaacgattcagtcataaaataaaggccttacgcatctcgagcgaaccagtacgagtgaaaattat

aacttcaaagtctgcccggcacgtgacttagagtctcccgcgagaccggcgctccggagttctggaatgc
agaagaagaagaagcattgtcacctaccgtgacgtcataaacgagtgtcttaaatagcgcctgcatcgag
ccatgcattctataacatcgctagccagccacgcatggtaaagttatcaagcgatagccagtcaatgata
tatagtagccaatgtctgttataaccaggcatggtcacatagcgtgttttaatacaagtagatctacatg
gtatatagttagcggcgaactaaagttttaaactgcagcaaattgcagacataaggtcttgcaaaagtag
cctacgagtattacattagccattcggtgatcaagttatttcacacaggaatcatgcttaatgttgagtt
tggaagttcgtaataatatgttagttgcaattaaactgcaagccttatcgttgcacgaagtatattatat
accagactttactatatatagaatcagaaagaatacagttacgaatccccgacgaaacaagtttctaata
ttaaatcgtgaaacattataaaagtgaatggattaaacttatcacgcttaaattatatcaagtcttcagg
gtatatacttttcgccggttctaaattatttttcgcaagacttgtttttttatacaaaatcttaacctaaa
cgaatgtgcagttataggtttaatattatgtgttacttgcaattacagtgcaaggagaaccgttgcaaaa
ttacaaccataagttgcaagatattaagattatgtattgacctcatatattttgtatttcagaaatctag
ccggtagtttgacatatttatacg

**Previously published sequences (hypothetical sequences given, not actual verified sequences):**

>CD4::GFP (Tolkin et al. 2016)[2] Homo sapiens CD4 GFP
ATGAACCGGGGAGTCCCTTTTAGGCACTTGCTTCTGGTGCTGCAACTGGCGCTCCTCCCAGCAGCCACTC
AGGGAAAGAAAGTGGTGCTGGGCAAAAAAGGGGATACAGTGGAACTGACCTGTACAGCTTCCCAGAAGAA
GAGCATACAATTCCACTGGAAAAACTCCAACCAGATAAAGATTCTGGGAAATCAGGGCTCCTTCTTAACT
AAAGGTCCATCCAAGCTGAATGATCGCGCTGACTCAAGAAGAAGCCTTTGGGACCAAGGAAACTTCCCCC
TGATCATCAAGAATCTTAAGATAGAAGACTCAGATACTTACATCTGTGAAGTGGAGGACCAGAAGGAGGA
GGTGCAATTGCTAGTGTTCGGATTGACTGCCAACTCTGACACCCACCTGCTTCAGGGGCAGAGCCTGACC
CTGACCTTGGAGAGCCCCCCTGGTAGTAGCCCCTCAGTGCAATGTAGGAGTCCAAGGGGTAAAAACATAC
AGGGGGGGAAGACCCTCTCCGTGTCTCAGCTGGAGCTCCAGGATAGTGGCACCTGGACATGCACTGTCTT
GCAGAACCAGAAGAAGGTGGAGTTCAAAATAGACATCGTGGTGCTAGCTTTCCAGAAGGCCTCCAGCATA
GTCTATAAGAAAGAGGGGGAACAGGTGGAGTTCTCCTTCCCACTCGCCTTTACAGTTGAAAAGCTGACGG
GCAGTGGCGAGCTGTGGTGGCAGGCGGAGAGGGCTTCCTCCTCCAAGTCTTGGATCACCTTTGACCTGAA
GAACAAGGAAGTGTCTGTAAAACGGGTTACCCAGGACCCTAAGCTCCAGATGGGCAAGAAGCTCCCGCTC
CACCTCACCCTGCCCCAGGCCTTGCCTCAGTATGCTGGCTCTGGAAACCTCACCCTGGCCCTTGAAGCGA
AAACAGGAAAGTTGCATCAGGAAGTGAACCTGGTGGTGATGAGAGCCACTCAGCTCCAGAAAAATTTGAC
CTGTGAGGTGTGGGGACCCACCTCCCCTAAGCTGATGCTGAGCTTGAAACTGGAGAACAAGGAGGCAAAG
GTCTCGAAGCGGGAGAAGGCGGTGTGGGTGCTGAACCCTGAGGCGGGGATGTGGCAGTGTCTGCTGAGTG
ACTCGGGACAGGTCCTGCTGGAATCCAACATCAAGGTTCTGCCCACATGGTCCACCCCGGTGCAGCCAAT
GGCCCTGATTGTGCTGGGGGGCGTCGCCGGCCTCCTGCTTTTCATTGGGCTAGGCATCTTCTTCTGTGTC
AGGactgtgtctagcaccATGGTGAGCAAGGGCGAGGAGCTGTTCACCGGGGTGGTGCCCATCCTGGTCG
AGCTGGACGGCGACGTAAACGGCCACAAGTTCAGCGTGTCCGGCGAGGGCGAGGGCGATGCCACCTACGG
CAAGCTGACCCTGAAGTTCATCTGCACCACCGGCAAGCTGCCCGTGCCCTGGCCCACCCTCGTGACCACC
CTGACCTACGGCGTGCAGTGCTTCAGCCGCTACCCCGACCACATGAAGCAGCACGACTTCTTCAAGTCCG
CCATGCCCGAAGGCTACGTCCAGGAGCGCACCATCTTCTTCAAGGACGACGGCAACTACAAGACCCGCGC
CGAGGTGAAGTTCGAGGGCGACACCCTGGTGAACCGCATCGAGCTGAAGGGCATCGACTTCAAGGAGGAC
GGCAACATCCTGGGGCACAAGCTGGAGTACAACTACAACAGCCACAACGTCTATATCATGGCCGACAAGC
AGAAGAACGGCATCAAGGTGAACTTCAAGATCCGCCACAACATCGAGGACGGCAGCGTGCAGCTCGCCGA
CCACTACCAGCAGAACACCCCCATCGGCGACGGCCCCGTGCTGCTGCCCGACAACCACTACCTGAGCACC
CAGTCCGCCCTGAGCAAAGACCCCAACGAGAAGCGCGATCACATGGTCCTGCTGGAGTTCGTGACCGCCG
CCGGGATCACTCTCGGCATGGACGAGCTGTACAAGTAA

>H2B::mCherry (Stolfi et al. 2010)[3] Helobdella (leech) H2B mCherry

ATGCCACCAAAGCCTGCCAGCAAGGGAGCTAAGAAGGCCGCCAGCAAGGCGAAAGCTGCTCGCAGCACGG
ACAAGAAGCACAAGAGAAGGCGAAAGGAAAGCTACTTTATATACATATACAAAGTGCTGAAGCAGGTTCA
CCCGGACACGGGCATCAGCGGCAAAGCCATGTCAATAATGAACTCGTTCGTCAATGACATCTTTGAACGA
ATCGCAGCCGAAGCTTCTCGCCTCGCCCACTACAACAAGAGATCCACAATCACCAGCAGAGAAATCCAGA
CGGCCGTCAGGCTTCTGTTACCCGGCGAGTTGGCCAAGCACGCCGTCAGCGAAGGCACCAAGGCCGTCAC
CAAGTACACCAGCTCAAAGgtcgacaggccaatctggccgcgggtcgacggtaccgcgggcccgggatcc
atcgccaccATGGTGAGCAAGGGCGAGGAGGATAACATGGCCATCATCAAGGAGTTCATGCGCTTCAAGG
TGCACATGGAGGGCTCCGTGAACGGCCACGAGTTCGAGATCGAGGGCGAGGGCGAGGGCCGCCCCTACGA
GGGCACCCAGACCGCCAAGCTGAAGGTGACCAAGGGTGGCCCCCTGCCCTTCGCCTGGGACATCCTGTCC
CCTCAGTTCATGTACGGCTCCAAGGCCTACGTGAAGCACCCCGCCGACATCCCCGACTACTTGAAGCTGT
CCTTCCCCGAGGGCTTCAAGTGGGAGCGCGTGATGAACTTCGAGGACGGCGGCGTGGTGACCGTGACCCA
GGACTCCTCCCTGCAGGACGGCGAGTTCATCTACAAGGTGAAGCTGCGCGGCACCAACTTCCCCTCCGAC
GGCCCCGTAATGCAGAAGAAGACCATGGGCTGGGAGGCCTCCTCCGAGCGGATGTACCCCGAGGACGGCG
CCCTGAAGGGCGAGATCAAGCAGAGGCTGAAGCTGAAGGACGGCGGCCACTACGACGCTGAGGTCAAGAC
CACCTACAAGGCCAAGAAGCCCGTGCAGCTGCCCGGCGCCTACAACGTCAACATCAAGTTGGACATCACC
TCCCACAACGAGGACTACACCATCGTGGAACAGTACGAACGCGCCGAGGGCCGCCACTCCACCGGCGGCA
TGGACGAGCTGTACAAGTAA

>Unc-76::GFP (Imai et al. 2009)[4] C. elegans Unc-76 tag GFP

ATGGCGGATCTGCGAGTACCGGACATTCCGCTCGCCTCGTGTGATGATGATGATATCGATAGTAATAAGA
ATTTGAGCAACCATTCATCAGACGAGAAACATCACTGCAACAGCAACAGCGACGAGGAACGTCTTCATGA
CGAGTTCTCTGGATCCCTTGAGGACCTTGTCGGCAACTTTGACGAAAAAATTGCGGCATGCCTGAAGGAC
CACGAGGTGACGACAGCGGATATTGCACCTGTGCAGATACGTACTCAAGAGGAAGTTATGAATGAAAGCC
AAACATGGTGGACATTAACCGGAAACTTTGGAAACATTCAACCTCTCGACTTTGGAACCTCTTCGATATG
TAAAAAGATGGCCGCAGCTCTGGACAGTGATTCATTGAAAGACGACGCATCTACACGCCGAAGTATGACA
AATTCCGATGATGAGGATCTTTTACGACAACAAATGGATGTTCATCAAATGATTGGACATCATCATGGAT
CTACGGATACTGGTGGTGAAACACCTCCACAGACTGCTGATCAAGTTATCGAAGAAATTGATGAAATGTT
ACAGgtaccggtcgccaccATGGTGAGCAAGGGCGAGGAGCTGTTCACCGGGGTGGTGCCCATCCTGGTC
GAGCTGGACGGCGACGTAAACGGCCACAAGTTCAGCGTGTCCGGCGAGGGCGAGGGCGATGCCACCTACG
GCAAGCTGACCCTGAAGTTCATCTGCACCACCGGCAAGCTGCCCGTGCCCTGGCCCACCCTCGTGACCAC
CCTGACCTACGGCGTGCAGTGCTTCAGCCGCTACCCCGACCACATGAAGCAGCACGACTTCTTCAAGTCC
GCCATGCCCGAAGGCTACGTCCAGGAGCGCACCATCTTCTTCAAGGACGACGGCAACTACAAGACCCGCG
CCGAGGTGAAGTTCGAGGGCGACACCCTGGTGAACCGCATCGAGCTGAAGGGCATCGACTTCAAGGAGGA
CGGCAACATCCTGGGGCACAAGCTGGAGTACAACTACAACAGCCACAACGTCTATATCATGGCCGACAAG
CAGAAGAACGGCATCAAGGTGAACTTCAAGATCCGCCACAACATCGAGGACGGCAGCGTGCAGCTCGCCG
ACCACTACCAGCAGAACACCCCCATCGGCGACGGCCCCGTGCTGCTGCCCGACAACCACTACCTGAGCAC
CCAGTCCGCCCTGAGCAAAGACCCCAACGAGAAGCGCGATCACATGGTCCTGCTGGAGTTCGTGACCGCC
GCCGGGATCACTCTCGGCATGGACGAGCTGTACAAGTAA

>Cas9 (Stolfi et al. 2014)[5] START NLS STOP

ATGGCTAGCCCCAAAAAGAAGAGGAAAGTGGACAAGAAGTATTCTATCGGACTGGACATCGGGACTAATA
GCGTCGGGTGGGCCGTGATCACTGACGAGTACAAGGTGCCCTCTAAGAAGTTCAAGGTGCTCGGGAACAC
CGACCGGCATTCCATCAAGAAAAATCTGATCGGAGCTCTCCTCTTTGATTCAGGGGAGACCGCTGAAGCA
ACCCGCCTCAAGCGGACTGCTAGACGGCGGTACACCAGGAGGAAGAACCGGATTTGTTACCTTCAAGAGA
TATTCTCCAACGAAATGGCAAAGGTCGACGACAGCTTCTTCCATAGGCTGGAAGAATCATTCCTCGTGGA
AGAGGATAAGAAGCATGAACGGCATCCCATCTTCGGTAATATCGTCGACGAGGTGGCCTATCACGAGAAA

TACCCAACCATCTACCATCTTCGCAAAAAGCTGGTGGACTCAACCGACAAGGCAGACCTCCGGCTTATCT
ACCTGGCCCTGGCCCACATGATCAAGTTCAGAGGCCACTTCCTGATCGAGGGCGACCTCAATCCTGACAA
TAGCGATGTGGATAAACTGTTCATCCAGCTGGTGCAGACTTACAACCAGCTCTTTGAAGAGAACCCCATC
AATGCAAGCGGAGTCGATGCCAAGGCCATTCTGTCAGCCCGGCTGTCAAAGAGCCGCAGACTTGAGAATC
TTATCGCTCAGCTGCCGGGTGAAAAGAAAAATGGACTGTTCGGGAACCTGATTGCTCTTTCACTTGGGCT
GACTCCCAATTTCAAGTCTAATTTCGACCTGGCAGAGGATGCCAAGCTGCAACTGTCCAAGGACACCTAT
GATGACGATCTCGACAACCTCCTGGCCCAGATCGGTGACCAATACGCCGACCTTTTCCTTGCTGCTAAGA
ATCTTTCTGACGCCATCCTGCTGTCTGACATTCTCCGCGTGAACACTGAAATCACCAAGGCCCCTCTTTC
AGCTTCAATGATTAAGCGGTATGATGAGCACCACCAGGACCTGACCCTGCTTAAGGCACTCGTCCGGCAG
CAGCTTCCGGAGAAGTACAAGGAAATCTTCTTTGACCAGTCAAAGAATGGATACGCCGGCTACATCGACG
GAGGTGCCTCCCAAGAGGAATTTTATAAGTTTATCAAACCTATCCTTGAGAAGATGGACGGCACCGAAGA
GCTCCTCGTGAAACTGAATCGGGAGGATCTGCTGCGGAAGCAGCGCACTTTCGACAATGGGAGCATTCCC
CACCAGATCCATCTTGGGGAGCTTCACGCCATCCTTCGGCGCCAAGAGGACTTCTACCCCTTTCTTAAGG
ACAACAGGGAGAAGATTGAGAAAATTCTCACTTTCCGCATCCCCTACTACGTGGGACCCCTCGCCAGAGG
AAATAGCCGGTTTGCTTGGATGACCAGAAAGTCAGAAGAAACTATCACTCCCTGGAACTTCGAAGAGGTG
GTGGACAAGGGAGCCAGCGCTCAGTCATTCATCGAACGGATGACTAACTTCGATAAGAACCTCCCCAATG
AGAAGGTCCTGCCGAAACATTCCCTGCTCTACGAGTACTTTACCGTGTACAACGAGCTGACCAAGGTGAA
ATATGTCACCGAAGGGATGAGGAAGCCCGCATTCCTGTCAGGCGAACAAAAGAAGGCAATTGTGGACCTT
CTGTTCAAGACCAATAGAAAGGTGACCGTGAAGCAGCTGAAGGAGGACTATTTCAAGAAAATTGAATGCT
TCGACTCTGTGGAGATTAGCGGGGTCGAAGATCGGTTCAACGCAAGCCTGGGTACCTACCATGATCTGCT
TAAGATCATCAAGGACAAGGATTTTCTGGACAATGAGGAGAACGAGGACATCCTTGAGGACATTGTCCTG
ACTCTCACTCTGTTCGAGGACCGGGAAATGATCGAGGAGAGGCTTAAGACCTACGCCCATCTGTTCGACG
ATAAAGTGATGAAGCAACTTAAACGGAGAAGATATACCGGATGGGGACGCCTTAGCCGCAAACTCATCAA
CGGAATCCGGGACAAACAGAGCGGAAAGACCATTCTTGATTTCCTTAAGAGCGACGGATTCGCTAATCGC
AACTTCATGCAACTTATCCATGATGATTCCCTGACCTTTAAGGAGGACATCCAGAAGGCCCAAGTGTCTG
GACAAGGTGACTCACTGCACGAGCATATCGCAAATCTGGCTGGTTCACCCGCTATTAAGAAGGGTATTCT
CCAGACCGTGAAAGTCGTGGACGAGCTGGTCAAGGTGATGGGTCGCCATAAACCAGAGAACATTGTCATC
GAGATGGCCAGGGAAAACCAGACTACCCAGAAGGGACAGAAGAACAGCAGGGAGCGGATGAAAAGAATTG
AGGAAGGGATTAAGGAGCTCGGGTCACAGATCCTTAAAGAGCACCCGGTGGAAAACACCCAGCTTCAGAA
TGAGAAGCTCTATCTGTACTACCTTCAAAATGGACGCGATATGTATGTGGACCAAGAGCTTGATATCAAC
AGGCTCTCAGACTACGACGTGGACCACATCGTCCCTCAGAGCTTCCTCAAAGACGACTCAATTGACAATA
AGGTGCTGACTCGCTCAGACAAGAACCGGGGAAAGTCAGATAACGTGCCCTCAGAGGAAGTCGTGAAAAA
GATGAAGAACTATTGGCGCCAGCTTCTGAACGCAAAGCTGATCACTCAGCGGAAGTTCGACAATCTCACT
AAGGCTGAGAGGGGCGGACTGAGCGAACTGGACAAAGCAGGATTCATTAAACGGCAACTTGTGGAGACTC
GGCAGATTACTAAACATGTCGCCCAAATCCTTGACTCACGCATGAATACCAAGTACGACGAAAACGACAA
ACTTATCCGCGAGGTGAAGGTGATTACCCTGAAGTCCAAGCTGGTCAGCGATTTCAGAAAGGACTTTCAA
TTCTACAAAGTGCGGGAGATCAATAACTATCATCATGCTCATGACGCATATCTGAATGCCGTGGTGGGAA
CCGCCCTGATCAAGAAGTACCCAAAGCTGGAAAGCGAGTTCGTGTACGGAGACTACAAGGTCTACGACGT
GCGCAAGATGATTGCCAAATCTGAGCAGGAGATCGGAAAGGCCACCGCAAAGTACTTCTTCTACAGCAAC
ATCATGAATTTCTTCAAGACCGAAATCACCCTTGCAAACGGTGAGATCCGGAAGAGGCCGCTCATCGAGA
CTAATGGGGAGACTGGCGAAATCGTGTGGGACAAGGGCAGAGATTTCGCTACCGTGCGCAAAGTGCTTTC
TATGCCTCAAGTGAACATCGTGAAGAAAACCGAGGTGCAAACCGGAGGCTTTTCTAAGGAATCAATCCTC
CCCAAGCGCAACTCCGACAAGCTCATTGCAAGGAAGAAGGATTGGGACCCTAAGAAGTACGGCGGATTCG
ATTCACCAACTGTGGCTTATTCTGTCCTGGTCGTGGCTAAGGTGGAAAAAGGAAAGTCTAAGAAGCTCAA
GAGCGTGAAGGAACTGCTGGGTATCACCATTATGGAGCGCAGCTCCTTCGAGAAGAACCCAATTGACTTT
CTCGAAGCCAAAGGTTACAAGGAAGTCAAGAAGGACCTTATCATCAAGCTCCCAAAGTATAGCCTGTTCG
AACTGGAGAATGGGCGGAAGCGGATGCTCGCCTCCGCTGGCGAACTTCAGAAGGGTAATGAGCTGGCTCT
CCCCTCCAAGTACGTGAATTTCCTCTACCTTGCAAGCCATTACGAGAAGCTGAAGGGGAGCCCCGAGGAC

AACGAGCAAAAGCAACTGTTTGTGGAGCAGCATAAGCATTATCTGGACGAGATCATTGAGCAGATTTCCG
AGTTTTCTAAACGCGTCATTCTCGCTGATGCCAACCTCGATAAAGTCCTTAGCGCATACAATAAGCACAG
AGACAAACCAATTCGGGAGCAGGCTGAGAATATCATCCACCTGTTCACCCTCACCAATCTTGGTGCCCCT
GCCGCATTCAAGTACTTCGACACCACCATCGACCGGAAACGCTATACCTCCACCAAAGAAGTGCTGGACG
CCACCCTCATCCACCAGAGCATCACCGGACTTTACGAAACTCGGATTGACCTCTCACAGCTCGGAGGGGA
TGAGGGAGCTCCCAAGAAAAAGCGCAAGGTAGGTTAATGA

>U6 promoter (Nishiyama et al. 2008)[6]
TGGCGGGTGTATTAAACCACTAAACAAACAATTGCCCCAAGCTCTCTTCACAATTATAAACACTATAATG
TTTGGACAAGAGATTAGCGTGGCTGTGACGAGAACTCTCAAAGGCTTGGTGTAATTGATATTTTATAAGA
AGCAGATTAAACTTCAATACAGTTTACACCTCATTTACAAAAAATTGGCTGCCAAAATCGCTAATTAACA
CATATTTAAAACAATTTCTACAGATATACACAGTATAGTATGATTACTAACTGCATAATAAACAAACATA
TCCAACAGACACTCACTAATCTGCCATAACAAGCTTCAAAAACTTAAACTCGAAATTTTAGTGAATCTTT
TTTTTTAAATGAAGATTTTATTTAAAAAGTTAAAAATATTACAGTTCAGGTATAGGTTTACACCTAATCT
TTAATAATCCGAACTAAATTTTAACTATTTAGAAACTTTTTCAACCAAAGTTTAAAAAAATAGATTCTTC
GCACGCTAAAACTATCATTTACACAAAAAAATGCAACAAAATGCAGAAAAAAATTACATTAGAGTTTAGG
TTAGTTACCTGCTAATCAATATAAACTAACTTCCCGCATAATATTCATCTAAAATTAGCAATAATCACGT
TTTACGCTAAAATTTGTGTAAAACTAAACTTCGTCCTTTGTCAAGGAGAAAATTTGACTCAAAAGCTGCG
CGCGCAGGGGAGATCCCCAAGCGAGTGTTTGTTACATCATAATCATGTGGAAAAATCCCCTAATAAGTAA
AAATACATATTTTTTAATTTTGGGGGCAAATAAACCGCTTTTTATGTCTAAAAACGCCAAAAATGGATCG
CGCGAGCCCAAAAACGCACAAATAACGTACAGACAGTGTCTCTGCGTACACAGACGGTATTTCCCCTTTA
AATTGAGAACTAGACTTAAGCACGCTTATAAGTCTGGAAGGCATCCGATGGTATAGAT

>Tbx6-r.b promoter (-1249/+12, Christiaen et al. 2009)[7]
caacggagtacgcgtgtcaagtttaatggcgtaattaccgaacaactgttgataagtaatgaggaccccg
ctgcggtaacctttcgaaattgctggttgcaagcggtgttggtgcgataataaggaaaagcggtaagcgt
tattttttgatccaccacaaagaccaaacctatattgatacactaaactaaatcaaattcgaacctatcc
attaaagtgcgaaatatacatagaatgaacaataatccgagctatattatgggcaatattttggctagtt
ttctgcttacaataatacataagaggcctacactaaatacggcggtatataaaactatgcaagacatacc
catatattagttttaatcaaattgatttaaaatatttttaaaactattaggcaactttgagtagaataag
ggtttaacctaaacgtatgtggttttagccatcgatggaaacgttttacaattatcgttactttttgaa
gacctttttcgttctatttttaagaagaacattcaaagaaatggaaaaccgtttcttacgaatcctatat
accgttgttaattgtttaaaaacacgataaggatattatgttctgaatgtgtcccatcttaccccacagc
actatataataaaattttagtttttgtgacagacttatgtcgcattcttaaacggatgatccccaactgg
ttaacaacgtaacaagcttgcaaaagcgaaggtgacattggtaacaacgtacgataactattgtctacta
ataacaatagacttaacacataacatatttaggaaatggcttcatatggcggactgtcaacttagttgtc
aaaattacctttaaatctctataaacgaagctgtttaataaaaaaaactaacaaggctattctacatcaa
accataataatgaaattatcaaacacattttaaagcgatttcatcaaaccaacgcgccacatgcaagac
ggtatgcgtcacactgagttttggagtgttctgcatgcgctagacttgaatcagcaggagagttcggagg
cttatcaggaagcagttgtccttgttaatgactcgttaagatcaaagtggcatcgaaaacgagtctcgct
ataaaacgggtttagtttcacagtacctcattccgctttctgttctcattggatatacaccaaactgaaa
gtacgagttaaatcgaaagagagaggaaattgtaagttttattggaccagacaagactatggcgaatatg

>Mesp promoter (-1916/-1, Davidson et al. 2005)[8]
cggttcaacgtgacgtcccatgccgatatcgtaaccatccggaacctctgatgctttttcaatatcatct
tttttgaaatccttcatttccgtttcatcgtcatttttgaaagccgggttctcatactcgtttttggtgc
tgtaaccgttttctgacatttttatctcatccaagtgcgaaccacttcaaagatggatagatacaccaga
ttttattacaaacaattagcaattcacgaaagttaaaaaaacgacataaaactataaaataaaattatac

ttattaaatatgaagaaaaatatgcattttaatcattctattgcaacaaatcggtattttcgtattct
cttatacgaagattgcatacaagcttaacgtttcatctgtttccattggaataaaatagaaaacgtccca
ccgcctgccctaacttttataagcttcacaaattatcaaagtttaccgttgaactcctgataaatgatt
attcagctattttaaacccaacttcgtttataagtgatgccgtctccttcttcgtatgaccccgctctac
taaatacatgtttgcacactgatcctaatgaagaccgcacctaaacctttacatagccgatgcattcaca
tataatgttctttgtgcaaccgaagtcgtatttgagcaaaccaaatatccaacttatataagtagacctt
taaaaatggacggtgaacaaaactttgcgaagtcatcataattgaaggtttatttagctctcctattgta
gggaaagatcctatggatgctccaaaacgataaacccgacttaccgaaaacgcggcggtcgttagaccat
gcttattgataaaccacatacatacaaactttagcgcaaactcggctatatctttacaacattagacaaa
ttctctttcatgtattttcatgggattataaagtgtctattttaggtaacaagactttaataaaaatatc
agtgtcccatattgttggcggctttgtttgcgagtgtctctgtattgacaacgtttataattaacttgga
agtaatgagtatacagcagcagacacagaagtttgtcgggggtatcctatgattgtacatcatgtgagca
aatatacttcaactgctagtgaataatatttgatatacgcacaagtcgtacaaatacacgactacaaata
agtcatacttgtcgaagtttgtattcccctgttttcgatcaatcttatcagccacaaaaaatggaaaaat
tccaaaaacgtagacacccacaaagtaacatacatggtaactcgtaagctaatacagagcacccgtgtga
aaacgactgtcgttttccggccacgcgagaataaagtaagttacattgattcattcactgacagtttccc
accaaaactgaagtattaatcaattatgaatcggcagtgctttcagtgggaagatttcggaaaatgtggg
tctaatgtcacagtaaactttgtgtttgttttaaaaacaatcatcgttaaatcaacatcgcaattctgtc
aatcactccacaataaactatatggctaatggaaacaggctgtttgtttgtatatcataaaatacttgcg
tcgtattatctgaccaaacaaaagcgttacaggaacggtccagcttcaaaatttgctgatgcatcagatt
ttacatttgaaatgtgattaattacgaaaatccagcgaatagaattgtcacaacaagtcattagcgacgg
atatttcgcctttgaaacttaaaggcgataatgactttgcccgtttcatgcggcgataaacgaactaatt
agacacctcctacagatataatggtaattcagaatcgtgtggttatgtaattcacaaaaacattttaaca
aaacaggttgatttgaaacttgtatt


1 Satou, Y. *et al.* A nearly complete genome of Ciona intestinalis Type A (C. robusta) reveals the contribution of inversion to chromosomal evolution in the genus Ciona. *Genome biology and evolution* **11**, 3144-3157 (2019).

2 Tolkin, T. & Christiaen, L. Rewiring of an ancestral Tbx1/10-Ebf-Mrf network for pharyngeal muscle specification in distinct embryonic lineages. *Development* **143**, 3852-3862 (2016).

3 Stolfi, A. *et al.* Early chordate origins of the vertebrate second heart field. *Science* **329**, 565 (2010).

4 Imai, K. S., Stolfi, A., Levine, M. & Satou, Y. Gene regulatory networks underlying the compartmentalization of the Ciona central nervous system. *Development* **136**, 285-293 (2009).

5 Stolfi, A., Gandhi, S., Salek, F. & Christiaen, L. Tissue-specific genome editing in Ciona embryos by CRISPR/Cas9. *Development* **141**, 4115-4120 (2014).

6 Nishiyama, A. & Fujiwara, S. RNA interference by expressing short hairpin RNA in the Ciona intestinalis embryo. *Development, growth & differentiation* **50**, 521-529 (2008).

7 Christiaen, L., Stolfi, A., Davidson, B. & Levine, M. Spatio-temporal intersection of Lhx3 and Tbx6 defines the cardiac field through synergistic activation of Mesp. *Developmental biology* **328**, 552-560 (2009).

8 Davidson, B., Shi, W. & Levine, M. Uncoupling heart cell specification and migration in the simple chordate *Ciona intestinalis*. *Development* **132**, 4811-4818 (2005).
