## Supplemental Table 1 for "Evolution of a chordate-specific mechanism for myoblast fusion"

**Supplementary Table 1**. sgRNA and oligonucleotide sequences.

| Purpose | Species | Gene | Sequence 5' → 3' | Amplicon (bp) | Application |
|---|---|---|---|---|---|
| CRISPR sgRNA | Human | MymK | sgRNA1: CTCACAGCTACAGAAGATGA | N/A | Fig. 1d |
| | | | sgRNA2: AAAGAAGAAGCGTAGCATCA | | |
| | Human | MymX | sgRNA1: GGCTCCCAGGACATGCGAG | | Fig. 4a |
| | | | sgRNA2: ACCTCTCCCTCCTCTCCAGG | | |
| | Mouse | MymK | sgRNA1: TAGCGATGCTCACTGTCGGG | | Extended Data Fig. 5a |
| | | | sgRNA2: GCGTCCTTACCATCGCTGTG | | |
| | | | sgRNA3: AGACAAACCAGGCCCATCAC | | |
| | Mouse | MymX | sgRNA1: GCTGCTGCCTGTTGCCCGCC | | Extended Data Fig. 11a |
| | | | sgRNA2: GAGGCCTCTCCAGAATCCGG | | |
| | | | sgRNA3: CCTCTGGGAGTGGTCCACTC | | |
| | Lizard | MymK | sgRNA1: CACGCCAAACATCACCAAGG | | Extended Data Fig. 5f |
| | | | sgRNA2: TTTGACGGTGATCATGAGGA | | |
| | *Ciona* | MymK | sgRNA2.6: gCAATGGAGTTGTGCAGAGG | | Fig. 2h |
| | | | sgRNA4.68: gCAGCTCCAAGTGCAAACGA | | |
| Genotype Analysis | Human | MymK | F: CTTCCTTCCCAGCCATCCAG<br>R: GGGCTAGTGAGCAGGGACTA | 492 | Fig. 1d |
| | Human | MymX | F: AACTGAAGGGAGGGGGAACT<br>R: TGGAGGACAGAGGGGCAATA | 599 | Fig. 4a |
| | Mouse | MymK | F: GCCTTTACCACCTTCTCCCC<br>R: CCCACCTCACACCTTCCTTC | 6,278 | Extended Data Fig. 5a |
| | Mouse | MymX | F: AGTTCAGGCTTCAGGTCAGAG<br>R: GCTAGGGGAGTGGGAACTGT | 743 | Extended Data Fig. 11a |
| | Lizard | MymK | F: CTTCATGTATGTTGGAAGGAGGCT<br>R: GTTGAGTGTAGACGCTTTTCTCTG | 395 | Extended Data Fig. 5f, g |
| | Lamprey | MymX | F: ATTGGAGGCACAGATGGTGG<br>R: GGGGTTTCACGACAAGCAAC | 498 | Fig. 3e |
| | *Ciona* | MymK | F: CGCGATCACAAATGACGAAAC<br>R: CCCGCAATTACAACATGCTAG | 1400 | Extended Data Fig. 8a |
| RT-PCR | Lamprey | 18s | F:CGCTTTGGTGACTCTGGATAA<br>R: GCCTGCCATCGTAAGTTGATA | 100 | Fig. 4e, f |
| | Lamprey | Myh1 | F: GTGAAGCAGAAGCTGGAGAA<br>R: ACGAGTTTCACCTTGGACTTT | 100 | Fig. 4f |
| | Lamprey | MymX | F: TTCTCTCCTGCGTGCTGTG<br>R: GCGTTCTTGTCATCGCCATC | 106 | Fig. 4e, f |
| | Lamprey | MymK | F: AGAATTCGAGGAGCCGCAG<br>R: AGGAACGCCCTCAGTTGGAT | 174 | Fig. 6c |
| | Lamprey | MymK | F1: CGGGGTGCGTACATAGACAG | 275 | Extended |

| | | | | | |
|---|---|---|---|---|---|
| Analyze mRNA isoforms | Lamprey | MymK | F2: GCACGTGGTTCCTCACCAA | 130 | Data Fig. 15 |
| | Lamprey | MymK | F3: GAGTGAGCGAACTGAAACACG | 187 | |
| | Lamprey | MymK | F4: TGGTCTGCGTGGCACG | 131 | |
| | Lamprey | MymK | R: TAGACCATGGCCTCGATGTG | N/A | |
| | Lamprey | MymK | F: AGTTAATTAAGGATCCGCCACC<br><u>ATGGGCTCCATCCTGGTC</u><br>R: CTGGCGGCCGCTCGAG<br><u>TCACACGCAGCAGCAGA</u> | 701 | Fig. 6b |
